# MOFA+: a probabilistic framework for comprehensive integration of structured single-cell data

**DOI:** 10.1101/837104

**Authors:** Ricard Argelaguet, Damien Arnol, Danila Bredikhin, Yonatan Deloro, Britta Velten, John C Marioni, Oliver Stegle

## Abstract

Technological advances have enabled the joint analysis of multiple molecular layers at single cell resolution. At the same time, increased experimental throughput has facilitated the study of larger numbers of experimental conditions. While methods for analysing single-cell data that model the resulting structure of either of these dimensions are beginning to emerge, current methods do not account for complex experimental designs that include both multiple views (modalities or assays) and groups (conditions or experiments). Here we present Multi-Omics Factor Analysis v2 (MOFA+), a statistical framework for the comprehensive and scalable integration of structured single cell multi-modal data. MOFA+ builds upon a Bayesian Factor Analysis framework combined with fast GPU-accelerated stochastic variational inference. Similar to existing factor models, MOFA+ allows for interpreting variation in single-cell datasets by pooling information across cells and features to reconstruct a low-dimensional representation of the data. Uniquely, the model supports flexible group-level sparsity constraints that allow joint modelling of variation across multiple groups and views.

To illustrate MOFA+, we applied it to single-cell data sets of different scales and designs, demonstrating practical advantages when analyzing datasets with complex group and/or view structure. In a multi-omics analysis of mouse gastrulation this joint modelling reveals coordinated changes between gene expression and epigenetic variation associated with cell fate commitment.

## Introduction

Single-cell methods have provided unprecedented opportunities to assay cellular heterogeneity. This is particularly important for studying complex biological processes, including the immune system, embryonic development and cancer^1–4^.

Following the establishment of the first scalable single-cell RNA sequencing (scRNA-seq) methods, other molecular layers are increasingly receiving attention, including single-cell assays for DNA methylation^5–9^ and chromatin accessibility^10–12^. More recently, technological advances enabled multiple biological layers to be probed in parallel in the same cells^12,13^, including: single-cell genome and transcriptome (G&T-seq)^14^, single-cell DNA methylation and transcriptome (scM&T-seq)^15^, single-cell chromatin accessibility and transcriptome (sci-CAR)^16^ and single-cell Nucleosome, Transcriptome and Methylation (scNMT-seq)^17^, among others^18–24^. These experimental techniques provide the basis for studying regulatory dependencies between transcriptomic and (epi)-genetic diversity at the single-cell level.

However, from a computational perspective, the integration of single-cell assays remains challenging owing to high degrees of missing data, inherent assay noise, and the scale of modern single-cell datasets, which can potentially span millions of cells. Previously, we introduced Multi-Omics Factor Analysis (MOFA), a statistical framework that addresses some of these challenges. However, the inference framework of MOFA is not designed to cope with increasingly-large scale datasets. Moreover, while MOFA is already devised to account for multiple views, the model makes strong assumptions about the dependencies across cells and in particular cannot account for additional structure between cells, e.g. batch, donors or conditions. By pooling and contrasting information across studies or experimental conditions, it would be possible to obtain more comprehensive insights into the complexity underlying biological systems^25–29^.

Other methods that have been proposed for integrating different data modalities include Seurat (v3) and LIGER, two strategies based on dimensionality reduction and manifold alignment^30,31^. Both methods anchor independent datasets from related populations of cells by exploiting the existence of a common feature space (for example matching gene expression and corresponding promoter accessibility). MOFA+, in contrast, is aimed at a different problem and allows for integrating data modalities via a common sample space (i.e. measurements derived from the same set of cells), where the features may be distinct across views.

## Model description

In a previous study, we developed Multi-Omics Factor Analysis (MOFA), a statistical framework for the integrative analysis of multiple data modalities^32^. Using a Bayesian Group Factor Analysis framework, MOFA infers a low-dimensional representation of the data in terms of a small number of (latent) factors that capture the global sources of variability (**Figure 1a**). While the model is applicable to single-cell assays, MOFA and related factor models have critical limitations, including their scalability and simplistic assumptions on the structure of the data. In particular, these models do not provide a principled approach for integrating multiple groups and views within the same inference framework.

**Figure 1:**
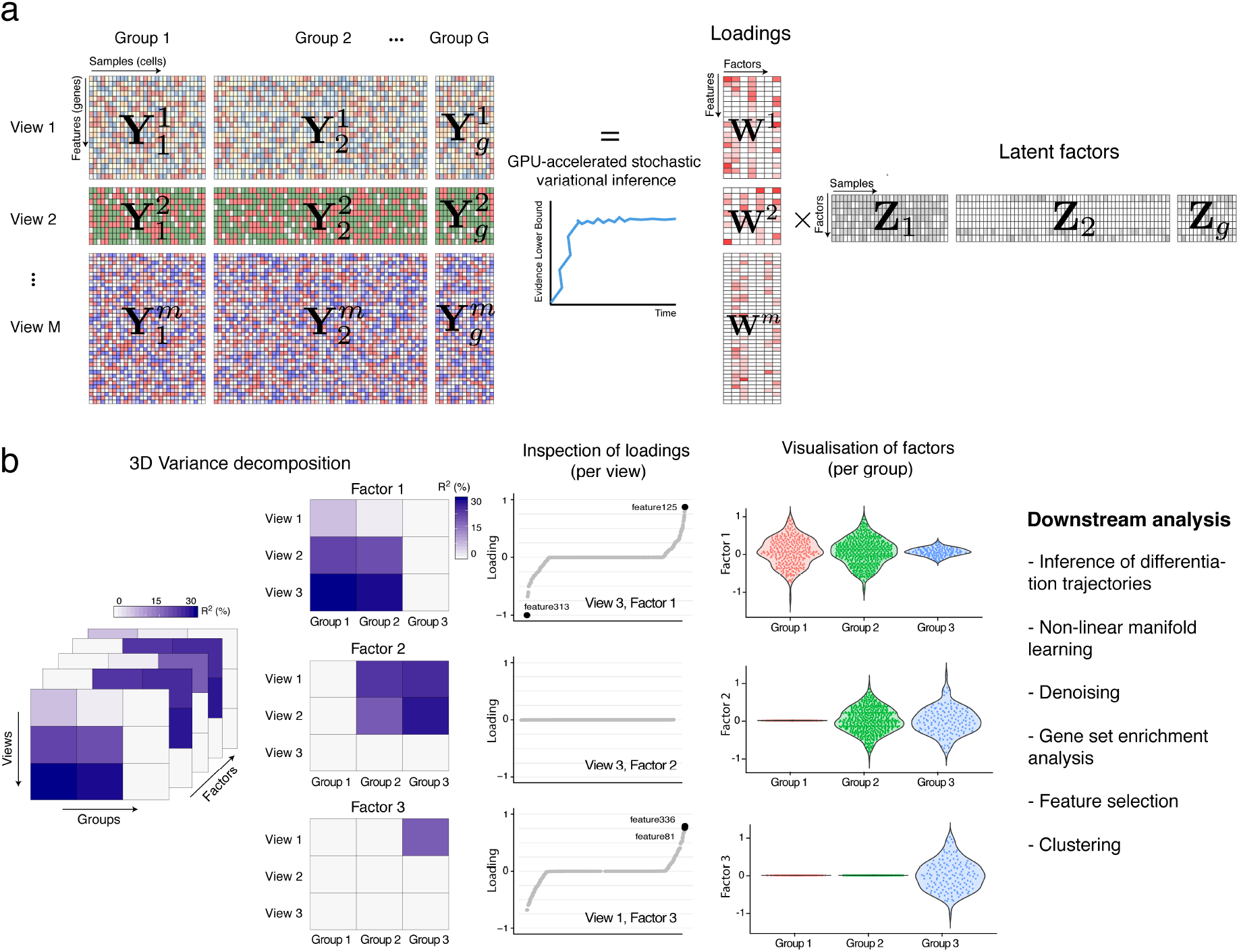
Multi-Omics Factor Analysis v2 (MOFA+) provides an unsupervised framework for the integration of multi-group and multi-view single-cell data. (a) Model overview: the input consists of multiple data sets structured into M views and G groups. Views consist of non-overlapping sets of features that can represent different assays. Analogously, groups consist of non-overlapping sets of samples that can represent different conditions or experiments. Missing values are allowed in the input data. MOFA+ exploits the dependencies between the features to learn a low-dimensional representation of the data (Z) defined by K latent factors that capture the global sources of molecular variability. For each factor, the weights (W) link the highdimensional space with the low-dimensional manifold and provide a measure of feature importance. The sparsity-inducing priors on both the factors and the weights enable the model to disentangle variation that is unique to or shared across the different groups and views. Model inference is performed using GPU-accelerated stochastic variational inference. (b) The trained MOFA+ model can be queried for a range of downstream analyses: 3D variance decomposition, quantifying the amount of variance explained by each factor in each group and view, inspection of feature weights, visualisation of factors and other applications such as clustering, inference of non-linear differentiation trajectories, denoising and feature selection.

In MOFA+ we address these challenges by i) developing a stochastic variational inference framework amenable to GPU computations, enabling inference with potentially millions of cells and ii) incorporating priors for flexible, structure regularisation, thus enabling joint modeling of groups of samples and multiple views.

Briefly, the inputs to MOFA+ are multiple datasets where features have been aggregated into non-overlapping sets of views and where cells have been aggregated into non-overlapping sets of groups (**Figure 1a**). Views generally correspond to different data modalities or omics (i.e. RNA expression, DNA methylation and chromatin accessibility), and groups to different experiments, batches or conditions. Importantly, the probabilistic framework underlying MOFA+ naturally handles missing values. During model training, MOFA+ infers K latent factors (per group) with associated feature weight matrices (per view) that explain the major axes of variation across the datasets. Importantly, the model provides sparsity-inducing priors to account for the structure of the data and to encourage sparse solutions to deliver interpretable solutions. After training, the model output enables a wide range of downstream analyses (**Figure 1b**), including variance decomposition, inspection of feature weights, inference of differentiation trajectories, and clustering, among others.

For technical details and mathematical derivations, we refer the reader to the **Methods** and the **Appendix.** A comparison of the model features with other factor analysis models is provided in **Table S1**.

## Model validation using simulated data

First, we validated the new features of MOFA+ using simulated data drawn from its generative model. The simulated data represented a range of dataset sizes with differing numbers of views and groups.

First, to assess the utility of stochastic variational inference, we trained different models either using conventional (deterministic) variational inference (VI), or using stochastic variational inference (SVI). Across a wide range of hyperparameter settings (see **Methods**) we observed that SVI yields Evidence Lower Bounds (i.e., values of the objective function) that are consistent with those obtained from conventional inference (**Figure S1**). However, the GPU-accelerated SVI implementation in MOFA+ achieved up to a ~20 fold increase in speed compared to VI, with the most dramatic speedups observed for large datasets. This inference scheme facilitates the application of MOFA+ to datasets comprising hundreds of thousands of cells using commodity hardware (**Figure S2**).

Next, we assessed the group-wise sparsity prior, by assessing to what extent it facilitates the identification of factors with simultaneous differential activity between groups and views. Indeed, when simulating data where factors explain differing amounts of variance across groups and across views, MOFA+ was able to more accurately reconstruct the true factor activity patterns than MOFA v1 or standard Bayesian Factor analysis (**Figure S3**).

## Integration of a heterogeneous time-course single-cell RNA-seq dataset

To illustrate the ability of MOFA+ to model multiple groups, we considered a time course scRNA-seq dataset (a single view), consisting of 16,152 cells that were isolated from multiple mouse embryos at embryonic days (E) 6.5, E7.0 and E7.25 (two biological replicates per stage). In this dataset, embryos are expected to contain similar subpopulations of cells but also to exhibit transcriptional differences due to variation in the rate of the developmental progression. As a proof of principle, we used MOFA+ to disentangle stage-specific variation from variation that is shared across all samples. For this purpose, we considered the six batches of cells (two replicates for each of the three embryonic stages) as different groups in the MOFA+ model.

MOFA+ identified 10 robust factors (Methods, **Figure S4**), capturing between 35% and 55% of the total transcriptional variance per embryo (**Figure 2a**). Some factors recapitulate the existence of post-implantation developmental cell types, including extra-embryonic (ExE) cell types (Factor 1 and Factor 2) and the transition of epiblast cells to nascent mesoderm via a primitive streak transcriptional state (Factor 4; **Figure 2b-c and Figure S5**). Consistently, the top weights for these factors are enriched for lineage-specific gene expression markers, including *Ttr* and *Apoa1* for ExE endoderm, *Rhox5* and *Bex3* for ExE ectoderm, and *Mesp1* and *Phlda2* for nascent mesoderm^33^. Other factors correspond to the formation of mesoderm-derived subpopulations that emerge from the caudal primitive streak after E7.0, including mesenchymal cells (Factor 9, **Figure S6**). MOFA+ also detects technical variation due to metabolic stress that affects all batches in a similar fashion (Factor 3, **Figure S7**).

**Figure 2:**
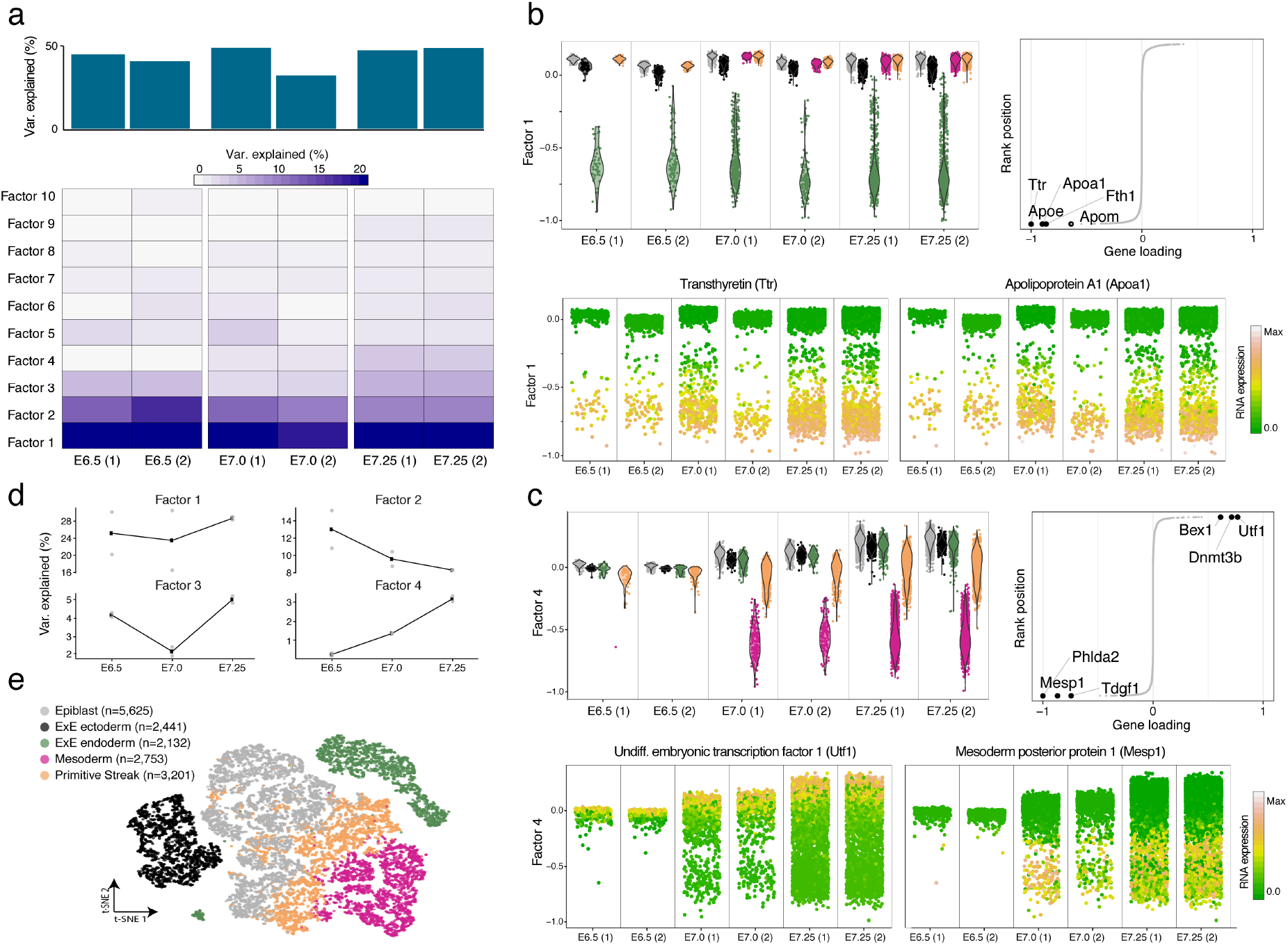
Integration of heterogeneous scRNA-seq experiments reveals stage-specific tran-scriptomic signatures associated with cell type commitment in mammalian development. (a) The heatmap displays the fraction of variance explained for each factor (rows) in each group (pool of mouse embryos at a specific developmental stage, columns). The bar plots show the variance explained per group with all factors. (b-c) Characterisation of Factor 1 as extra-embryonic (ExE) endoderm formation (b) and Factor 4 as Mesoderm commitment (c). In each panel the top left plot shows the distribution of factor values for each batch of embryos. Cells are coloured by cell type. Line plots (top right) show the distribution of gene weights, with the top five genes with largest (absolute) weight highlighted. The bottom beeswarm plots represent the distribution of factor values, with cells coloured by the expression of the genes with highest weight. (d) Line plots show the amount of variance explained (%, averaged across the two biological replicates) for each factor as a function of time. The value of each replicate is shown as grey dots. (e) Dimensionality reduction using t-SNE on the 10 inferred factors. Cells are coloured by cell type.

When inspecting the factor activity across development, we observe that the percentage of variance explained by Factor 1 is not correlated with developmental progression, indicating that commitment to ExE endoderm fate occurs early in the embryo and that the proportion of this cell type remains relatively constant from E6.5 to E7.25. In contrast, the amount of variance explained by Factor 4 increases over time (**Figure 2d**), consistent with a higher proportion of cells committing to mesoderm after ingression through the primitive streak.

All together, this application shows how MOFA+ can identify biologically meaningful structure in scRNA-seq datasets with multiple groups. Interpretability is achieved at the expense of limited information content per factor (due to the linearity assumption). Nevertheless, the MOFA factors can be used as input to other methods that learn compact nonlinear manifolds that discriminate cell types (**Figure 2e**) and enable the reconstruction of pseudotime trajectories^34,35^.

## Identification of context-dependent methylation signatures associated with cellular diversity in the mammalian cortex

As a second use case, we considered how MOFA+ can be used to investigate variation in epigenetic signatures between populations of neurons. This application illustrates how a multi-group and multi-view structure can be defined from seemingly uni-modal data, which can then be used to test specific biological hypotheses.

We analysed 3,069 cells isolated from the frontal cortex of young adult mice, where DNA methylation was profiled using single-cell bisulfite sequencing^7^. Recent studies have demonstrated that neurons contain significant levels of non-CpG methylation (mCH), an epigenetic mark that has been historically dismissed as a methodological artifact of incomplete bisulfite conversion^36–39^.

Here we used MOFA+ to dissect the degree of coordination between mCH and mCG signatures in different regions of the brain. As input data we quantified mCH and mCG levels at gene bodies, promoters and putative enhancer elements (Methods). Each combination of genomic and sequence context (e.g., mCG at enhancer elements) was defined as a separate view. To explore the influence of the neuron’s location we grouped cells according to their cortical layer: Deep, Middle or Superficial (**Figure S8**).

The sparseness of DNA methylation data results in large amounts of missing values, which hampers the use of conventional dimensionality reduction techniques such as PCA or NMF^34,35,40^. By contrast, the probabilistic framework underlying MOFA+ naturally accounts for missing values^32^.

MOFA+ identifies 5 robust factors (Methods; **Figure S9**) that explain the structured heterogeneity across genomic contexts and cortical layers. Factor 1, the major source of variation, is linked to the existence of inhibitory and excitatory neurons. This factor shows significant mCG activity across all cortical layers, mostly driven by coordinated changes in enhancer elements, but to some extent also gene bodies (**Figure 3a,b**). Consistently, the top weights in the mCG:gene body view are enriched for genes whose RNA expression has been shown to discriminate between the two classes of neurons, including *Neurod6* and *Nrgn^7^*. In addition, this analysis identifies novel genes with differential gene body mCG levels that may have yet unknown roles in defining the epigenetic landscape of neuronal diversity, including *Vsig2, Taar3* and *Cort* (**Figure S10**).

**Figure 3:**
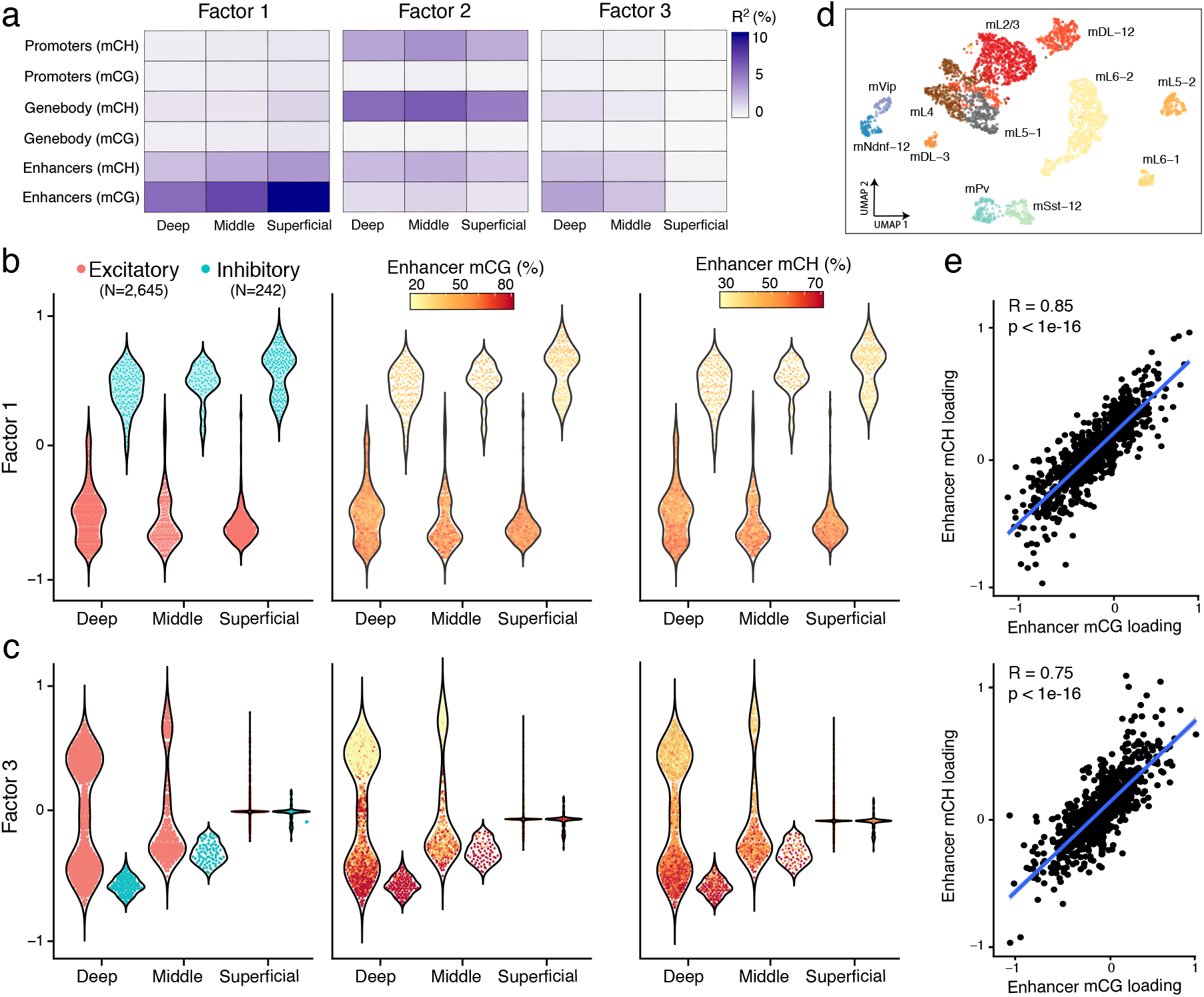
MOFA+ reveals context-dependent DNA methylation signatures associated with cellular diversity in the mammalian cortex. (a) Variance explained for each factor across the different groups (cortical layer, x-axis) and views (genomic context, y-axis). For simplicity, only the first three factors are shown. (b-c) Characterisation of (b) Factor 1 as the two major neuron populations and (c) Factor 3 as increased cellular diversity of excitatory neurons in deep cortical layers. The beeswarm plots show the distribution of factor values for each group, defined as the neuron’s cortical layer. In the left plot, cells are coloured by neuron class. In the middle and right plots the cells are coloured by average mCG and mCH levels (%), respectively, of the top 100 enhancers with the largest weights. (d) UMAP projection of the MOFA factors. Each dot represents a cell, coloured by maximally resolved cell type assignments. (e) Correlation of enhancer mCG weights (x-axis) and mCH weights (y-axis) for Factor 1 (top) and Factor 3 (bottom).

Factor 2 captures genome-wide differences in global mCH levels (R=0.99), which is moderately correlated with differences in global mCG levels (R=0.32) (**Figure S11**). Factor 3 captures heterogeneity linked to the increased cellular diversity along cortical depth, with the Deep layer displaying significantly more diversity of excitatory cell types than the Superficial layer (**Figure 3a,c**). Again, we observe that the MOFA+ factors are a suitable input to learn non-linear manifolds and reveal the existence of subpopulations of both excitatory and inhibitory cell types (**Figure 3d**). Notably, the MOFA+ factors are significantly better at identifying subpopulations than the conventional approach of using Principal Component Analysis with imputed measurements (**Figure S12**).

Interestingly, in addition to the dominant mCG signal, MOFA+ connects Factor 1 and Factor 3 to variation in mCH, which could suggest a role of mCH in cellular diversity. We hypothesise that this can be supported if the genomic regions that show mCH signatures are different than the ones marked by the conventional mCG signatures. To investigate this, we correlated the mCH and mCG feature weights for each factor and genomic context (**Figure 3e and Figure S13**). In all cases we observe a strong positive dependency, indicating that mCH and mCG signatures are spatially correlated and target similar loci.

Taken together, this result supports the hypothesis that mCH and mCG tag the same genomic loci and are associated with the same sources of variation, suggesting that the presence of mCH may be the result of non-specific *de novo* methylation as a by-product of the establishment of mCG^36^.

## MOFA+ reveals molecular signatures of lineage commitment during mammalian embryogenesis

As a final application, we considered a complex dataset with multiple sample groups and views. The dataset consists of a multi-omic atlas of mouse gastrulation where scNMT-seq was used to simultaneously profile RNA expression, DNA methylation and chromatin accessibility in 1,828 cells at multiple stages of development^41^. In this dataset MOFA+ provides a method for delineating coordinated variation between the transcriptome and the epigenome and for detecting at which stage(s) of development it occurs.

As input to the model we quantified DNA methylation and chromatin accessibility values over two sets of regulatory elements: gene promoters and enhancer elements (distal H3K27ac sites^41–43^). RNA expression was quantified over protein-coding genes. After data processing (**Methods**), separate views were defined for the RNA expression and for each combination of genomic context and epigenetic readout. Cells were grouped according to their developmental stage (E5.5, E6.5 and E7.5), reflecting the underlying experimental design (**Figure S14**). Notably, the epigenetic readouts are extremely sparse, with, on average, only 18% and 10% of cells having recorded data at a gene promoter for DNA methylation and chromatin accessibility, respectively. In this context, methods that pool information across cells and features are essential for robust inference.

MOFA+ identifies 8 robust factors with a minimum variance explained of 1% in the gene expression view. The first factor captured the formation of ExE endoderm, a cell type that is present across all stages (**Figure 4a**), in agreement with our previous results using the independently generated transcriptomic atlas of mouse gastrulation (**Figure 2**). MOFA+ links Factor 1 to changes across all molecular layers. Notably, the distribution of weights for DNA methylation are skewed towards negative values (at both enhancers and promoters), indicating that ExE endoderm cells are characterised by a state of global demethylation, consistent with previous studies^44^.

**Figure 4:**
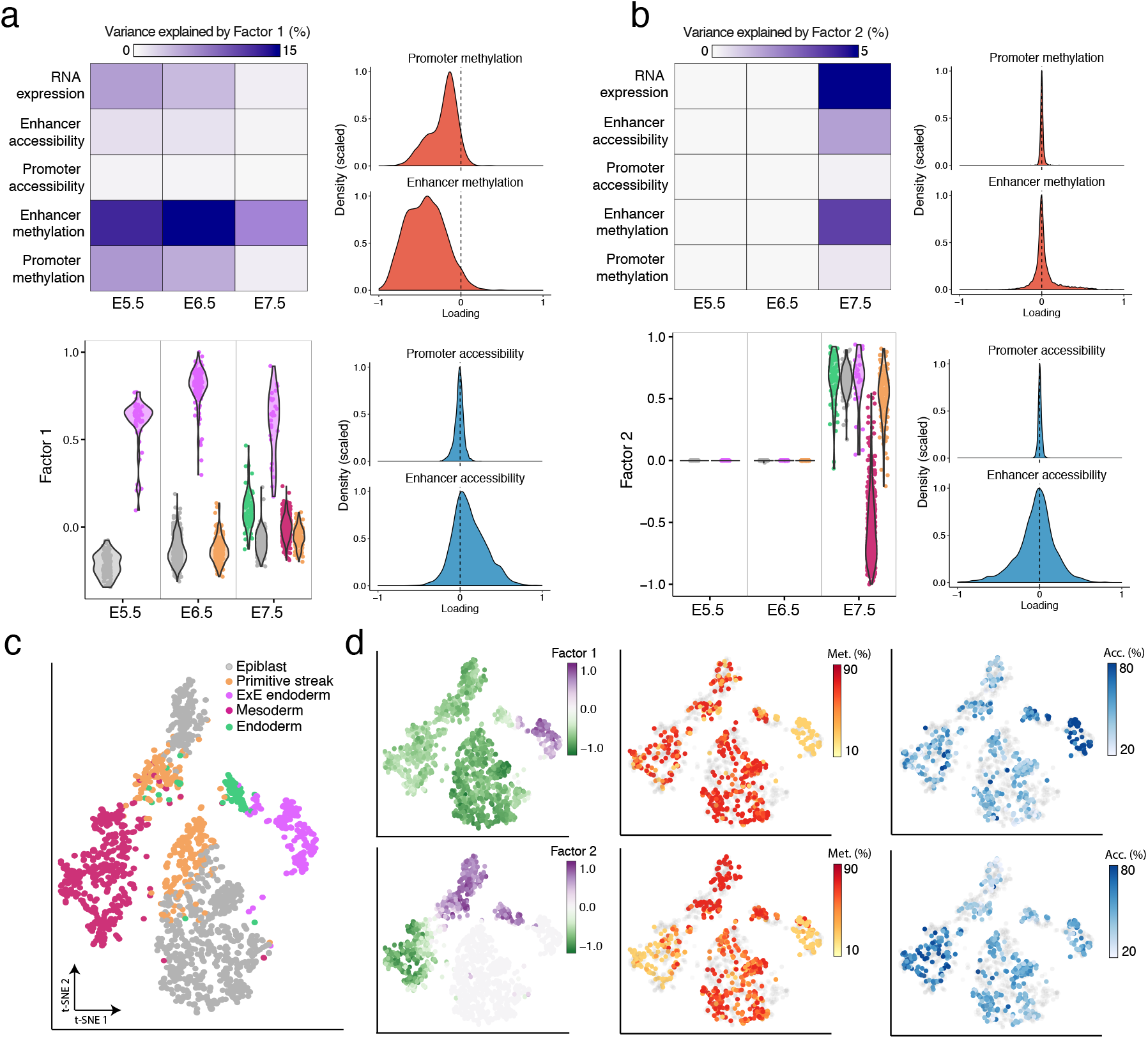
MOFA+ integrates multi-modal scNMT-seq experiments to reveal epigenetic signatures associated with lineage commitment during mammalian embryogenesis. (a-b) Characterisation of Factor 1 as ExE endoderm formation and Factor 2 as Mesoderm commitment. Top left plot shows the variance explained by the factor across the different views (rows) and groups (embryonic stages, as columns). Bottom left plot shows the distribution of factor values for each stage, coloured by cell type assignment. Histograms display the distribution of DNA methylation and chromatin accessibility weights for promoters and enhancer elements. (c) Dimensionality reduction using t-SNE on the inferred MOFA factors. Cells are coloured by cell type. (d) Same as (c), but cells are coloured by Factor 1 values (top left) and Factor 2 values (bottom left); by the DNA methylation levels of the enhancers with the largest weight in Factor 1 (top middle) and Factor 2 (bottom middle); by the chromatin accessibility levels of the enhancers with the largest weight in Factor 1 (top right) and Factor 2 (bottom right).

The next factors captured the molecular variation associated with the emergence of the primary germ layers at E7.5: mesoderm (Factor 2, **Figure 4b**), and embryonic endoderm (Factor 4, **Figure S15**). Again, for both factors, MOFA+ connects the transcriptome variation to changes in DNA methylation and chromatin accessibility. Yet, in striking contrast to Factor 1, the variance decomposition analysis and the distribution of weights indicate that the epigenetic dynamics are mostly driven by enhancer elements. Little coordinated variation is observed in promoters (**Figure 4b**), even for genes that show strong differential expression between germ layers (**Figure S16**). These results are in agreement with other studies that pinpointed distal regulatory elements as a major target of epigenetic modifications during embryogenesis^45–47^.

The remaining factors capture variation that is mostly driven by the RNA expression, whose etiology can be related to the existence of morphogenic gradients (Factor 8, **Figure S17**), the emergence of other cellular subpopulations during gastrulation (Factor 7, **Figure S18**) and cell cycle (Factor 6, **Figure S19**).

In conclusion, the MOFA+ output suggests that independent cell fate commitment events undergo different modes of epigenetic variation. While some lineages manifest global changes in the epigenetic landscape (ExE endoderm, Factor 1), other cell types are associated with the emergence of local epigenetic patterns that are driven by specific genomic contexts (embryonic endoderm and mesoderm, Factors 2 and 4).

## Discussion

As single-cell technologies mature, they are applied to generate data sets of increasing complexity, with highly structured and sparse measurements^16,17,24,48,49^. Consequently, there is a need for integrative computational frameworks that can robustly and systematically interrogate the data generated in order to reveal the underlying sources of variation^25^.

In this study we introduced MOFA+, a generalisation of the MOFA framework^32^ that facilitates analysis of large-scale datasets with complex multi-group and/or multi-view experimental designs. From a technical perspective, MOFA+ provides two major features: first, GPU-accelerated stochastic variational inference ensures scalability to potentially millions of cells; second, structured sparsity priors provide a principled inference framework to jointly analyse multiple data sets. Additionally, MOFA+ inherits all the features from its predecessor, including a natural approach for handling missing values as well as the capacity to perform inference with non-Gaussian readouts^32^.

Although MOFA+ represents an important step forward in the analysis of single-cell omics data, it has some limitations. First, it requires multi-modal measurements from the same set of cells. This contrasts with other integrative frameworks such as Seurat^31^ or LIGER^30^, which anchor data sets based on the assumption of a common feature space (e.g. matching gene expression and promoter accessibility). Second, the model is not currently able to capture strong non-linear relationships (**Figure S20**). We speculate that this could be addressed by combining MOFA with concepts from variational autoencoders, as recently proposed for the analysis of scRNA-seq data^50–52^. Third, the model assumes independence between features in its prior distributions, despite the fact that genomic features are known to interact via complex regulatory networks^53^.

To facilitate adoption of the method, we deploy MOFA+ as open-source software with multiple tutorials and a web-based analysis workbench, to support for a large variety of downstream analysis, enabling a user-friendly in-depth characterisation of structured single-cell data.

## Methods

### Multi-Omics Factor Analysis v2 model (MOFA+)

The input to MOFA+ is a list of matrices, each matrix containing a predefined group of cells (group) and a predefined set of features (view, see Figure 1 for a visual representation).

We introduce the following notation: M for the number of views, D_m_ for the number of features in the *m*-th view, G for the number of groups, N_g_ for the number of samples in the *g*-th group and K for the number of factors.

As in the original version of MOFA^32^, the underlying master equation is the standard matrix factorisation framework:

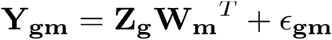

- *Y_gm_* denotes the matrix of observations for the *m*-th view and the *g*-th group.
- *W_m_* denotes the weight matrix for the *m*-th view
- *Z_g_* denotes the factor matrix for the *g*-th group
- *ε_gm_* denotes the residual noise for the *m*-th view and the *g*-th group. The specific form of the noise can be tailored to the nature of the data type^32^

The factor matrix *Z_g_* has dimensionality (N_g_,K) and contains the low-dimensional representation of the samples from the *g*-th group. The weight matrix *W_m_* has dimensionality (D_m_,K) and contains an association score for each feature with each factor. The noise matrix *ε_gm_* contains the unexplained variance (i.e. noise) for each feature in each group.

The model is formulated in a probabilistic Bayesian setting. We introduce prior distributions on all unobserved variables of the model in order to induce specific regularisation criteria, as described below.

### Interpretation of the factors

The MOFA+ Factors capture global sources of variability in the data. The factors matrices express how much the MOFA+ Factors are active within the various groups of cells. Mathematically, each factor orders cells along a one-dimensional axis centered at zero. Samples with different signs have opposite effects along the inferred axis of variation. Cells that remain centered at zero represent either an intermediate phenotype or no phenotype at all associated with the factor under consideration.

### Interpretation of the weights

The weights matrices provide a score for how strong each feature relates to each factor, hence allowing a biological interpretation of the MOFA+ Factors. Genes with no association with the factor have values close to zero, while genes with strong association with the factor have large absolute values. The sign of the loading indicates the direction of the effect: a positive loading indicates that the feature has higher levels in the cells with positive factor values, and vice versa.

### Model regularisation

The regularisation of the weights and the factors is critically important for enabling MOFA to perform inference with structured data sets.

In the original version of MOFA, structured priors were applied to the weights to enable inference and interpretable outputs of multi-view data sets (i.e. structured features but not samples were facilitated). In MOFA+ we generalised this by introducing a symmetric regularisation for both the factors and weights, hence accounting for structure in both the sample and the feature space (see appendix for mathematical details). The main purpose is to enable the identification of factors that are active in different subsets of both groups and views.

The first level of sparsity uses an Automatic Relevance Determination prior to explicitly model differential activity of factors across views and/or across groups. The second level of sparsity uses a spike-and-slab prior to simultaneously push individual weights and factors to zero. This effectively encourages sparse solutions where factors are (potentially) associated with a small number of active features and/or active within small subsets of samples. Using feature-wise sparsity priors helps disentangling technical and biological sources of variability^54^.

### Noise model

MOFA+ supports a variety of different likelihood models to enable integration of diverse combinations of data types. These include a Gaussian noise model for continuous data, a Poisson model for count data and a Bernoulli model for binary data. This feature is inherited from MOFA^32^.

### Statistical variational inference

In MOFA, inference was performed using mean-field variational Bayes^55–57^. While this framework is typically faster than sampling-based Monte Carlo approaches, it becomes prohibitively slow when applied to large single-cell datasets. In MOFA+ we implemented a stochastic version of the algorithm^57,58^ that can be accelerated by performing computation using GPUs.

The use of stochastic variational inference comes at the cost of introducing additional hyperparameters: batch size (number of cells used to compute the gradients), learning rate (step size) and forgetting rates (rate of decay of the learning rate). While we find the hyperparameters to be robust across a variety of simulated data (**Figure S1**), their optimisation is likely to be important in some contexts. By default we use GPU-powered standard variational inference if the full data set fits in the GPU. Otherwise, we perform stochastic variational inference using a batch size of 50%, a learning rate of 0.5 and a forgetting rate of 0.25.

### Variance decomposition

Once the model is trained, we can quantify how much of the observed variance is explained by each factor *k* in each group *g* and in each view *m.* This is estimated using a coefficient of determination:

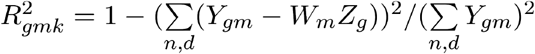

### Gene set enrichment analysis

Gene set enrichment analysis was performed using the Reactome gene sets^59^. For every gene set G, we evaluate its significance via a parametric t-test, where we contrast the weights of the foreground set (features that belong to the set G) versus the background set (the weights of features that do not belong to the set G). Resulting P-values were adjusted for multiple testing for each factor using the Benjamini–Hochberg procedure^60^. Significant enrichments were at a false discovery rate of 1%.

### 10x: data processing

The gastrulation scRNA-seq atlas was obtained from^33^ and is available in the Gene Expression Omnibus under accession GSE87038. Cells were subset to stages E6.5, E7.0 and E7.25. Cells from stage E6.75 were not included in the analysis because they consist of a single biological replicate. Gene expression counts were normalised using *scran^61^.* The 5,000 most overdispersed genes after regressing out the stage effect were selected prior to fit the model. Details on the quality control and data preprocessing can be found in^33^.

### Ecker: data processing

The mouse brain DNA methylation data was obtained from^7^ and is available in the Gene Expression Omnibus under accession GSE97179. DNA methylation was quantified over genomic features using a binomial model where the number of successes is the number of reads that support methylation (or accessibility) and the number of trials is the total number of reads. A CpG methylation rate was calculated for each genomic feature and cell using a maximum likelihood approach. The rates were subsequently transformed to M-values^62^ and modelled with a Gaussian likelihood.

As input to MOFA+ we filtered genomic features with low coverage (at least 3 CpG measurements or at least 10 CH measurements) and we selected the intersection of the top 5000 most variable sites across the different genomic and sequence contexts (see **Figure S8**). Details on the quality control and data preprocessing can be found in^7^.

### scNMT: data processing

The gastrulation scNMT-seq multi-omics data was obtained from^41^and is available in the Gene Expression Omnibus under accession GSE121708

Gene expression counts were quantified over protein-coding genes using featureCounts^63^ with the Ensembl gene annotation 87^64^. The read counts were log-transformed and size-factor adjusted, and modelled with a Gaussian likelihood. As input to MOFA+, we filtered genes with a dropout rate higher 90% and we subsetted the top 5,000 most variable genes (after regressing out the stage effect). In addition, batch effects and the dropout rate per cell were regressed out prior to fitting the model.

DNA methylation and chromatin accessibility data were quantified over genomic features using a binomial model where the number of successes is the number of reads that support methylation (or accessibility) and the number of trials is the total number of reads. A CpG methylation or GpC accessibility rate for each genomic feature and cell was calculated by maximum likelihood. The rates were subsequently transformed to M-values^62^ and modelled with a Gaussian likelihood. As input to MOFA+ we filtered genomic features with low coverage (at least 3 CpG and 5 GpC measurements) and we selected the top 2500 most variable sites per combination of genomic context and data modality (see **Figure S14**) Details on the quality control and data preprocessing can be found in^41^.

### Software availability

An open-source implementation of MOFA+ is available from https://github.com/bioFAM/MOFA2, which includes vignettes to reproduce the analyses presented in this article.

Also, we deploy an interactive web-based platform to facilitate the exploration of MOFA+ models (**Figure S21**). This is available from https://github.com/gtca/mofaplus-shiny

### Competing interests

All authors declare no competing financial interests

## Acknowledgements

R.A. is a member of Robinson College at the University of Cambridge. We thank Florian Buettner for comments on the manuscript.

## Author contributions

R.A., D.A. and B.V. conceived the project.

R.A., D.A. D.B., Y.D and B.V. implemented the model.

R.A., D.A., J.C.M and O.S., interpreted results.

D.A. implemented the interactive web-based platform.

R.A. generated figures.

R.A., wrote the manuscript with feedback from all authors

J.C.M and O.S. supervised the project.

---

**Figure S1.**
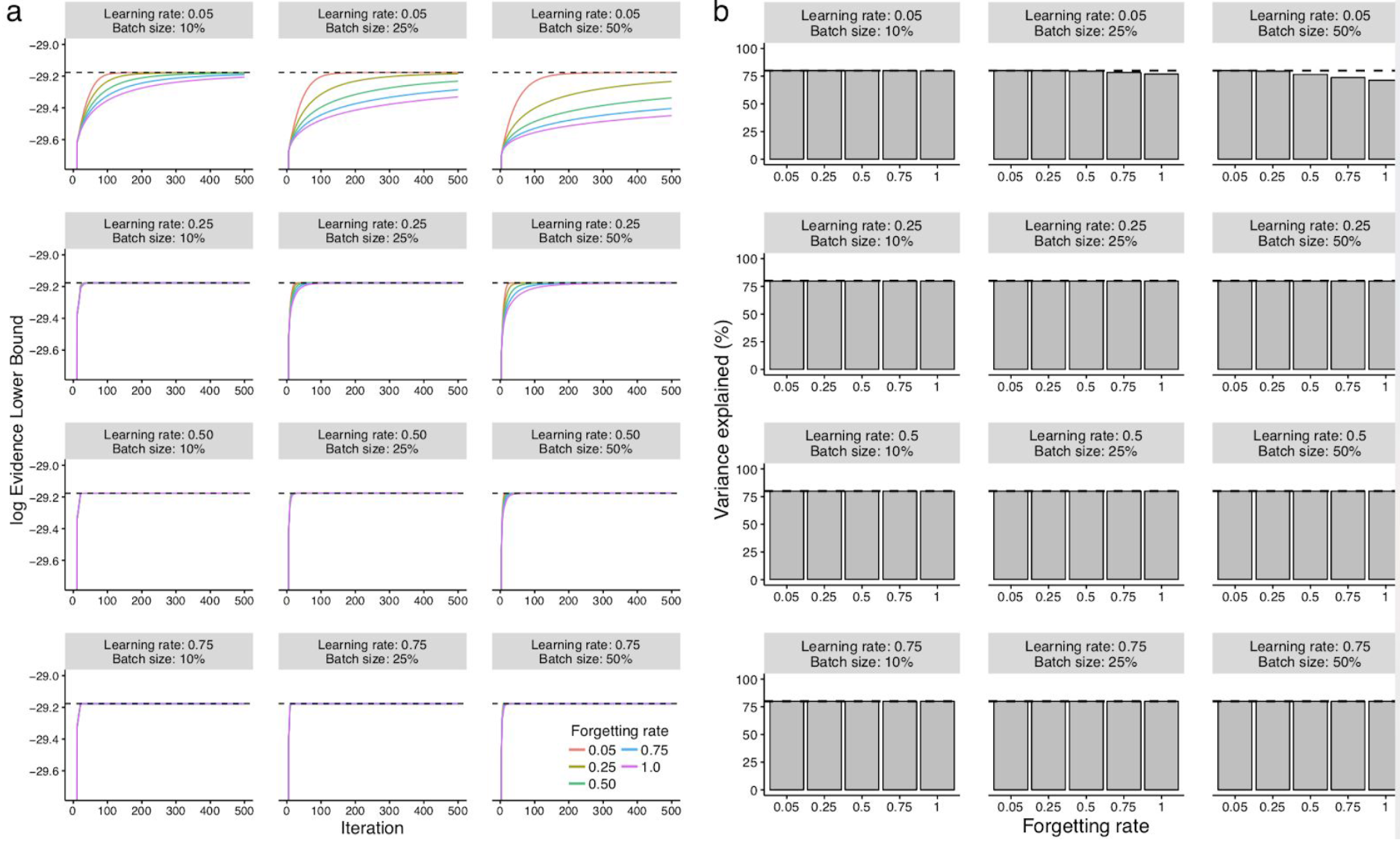
Validation of stochastic variational inference using simulated data. Data is simulated from the generative model with the following parameters: M=3 views, G=3 groups, D=1,000 features (per view), N=100,000 samples (per group) and K=25 factors. (a) Line plots display the iteration number of the inference (x-axis) and the log-Evidence Lower Bound (ELBO) on the y-axis. Panels correspond to different values of batch sizes (10%, 25%, 50% of the data) and initial learning rates (0.05, 0.25, 0.5, 0.75). Colors correspond to different forgetting rates (0.05, 0.25, 0.5, 0.75, 1.0). The dashed horizontal line indicates the ELBO achieved using standard variational inference. (b) Bar plots display the forgetting rate (x-axis) and the total variance explained (%) in the y-axis. Panels correspond to different values of batch sizes (10%, 25%, 50% of the data) and initial learning rates (0.05, 0.25, 0.5, 0.75). The dashed line indicates the variance explained (%) achieved using standard variational inference.

**Figure S2.**
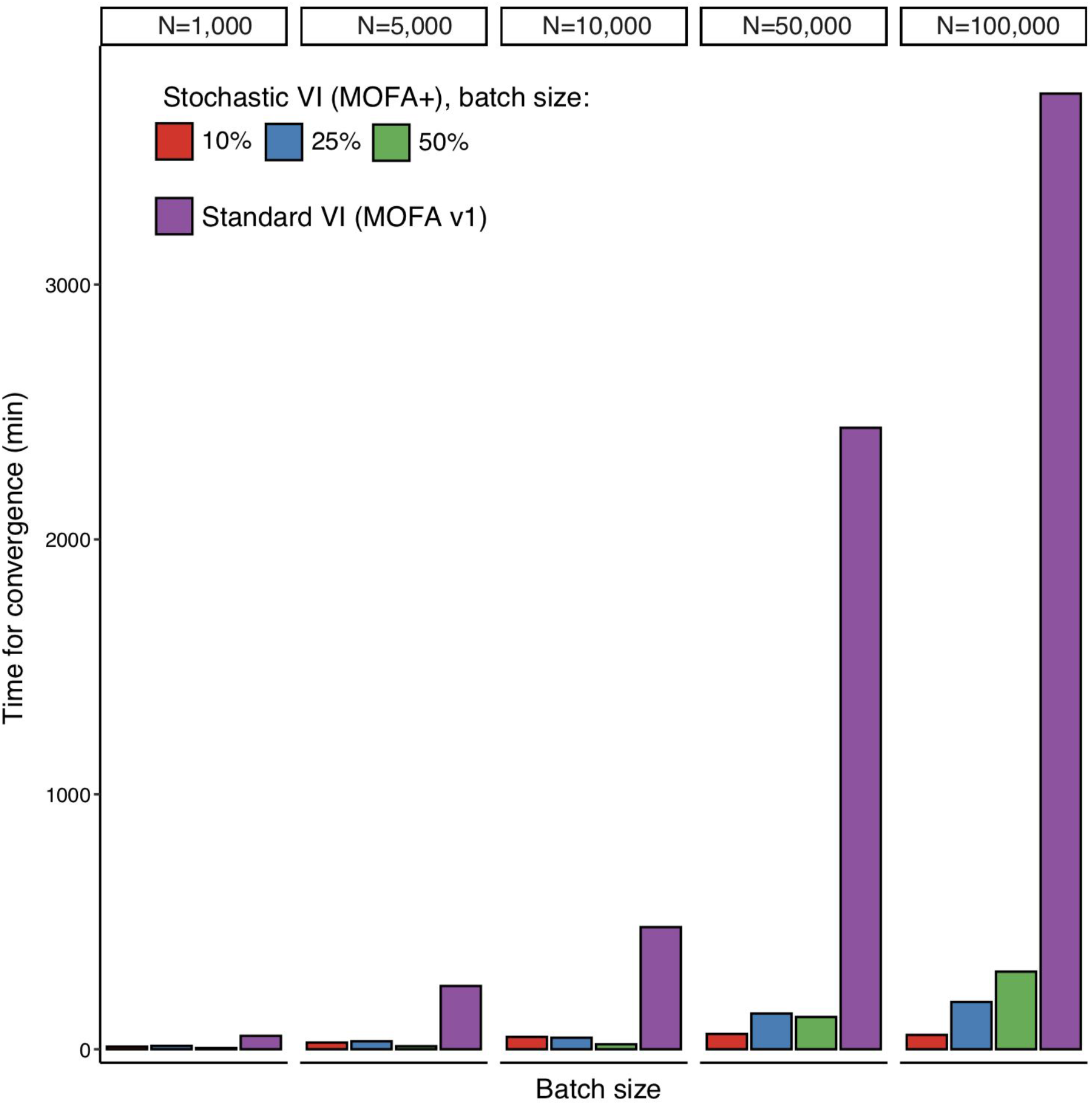
Evaluation of convergence speed for stochastic variational inference using simulated data. Data is simulated from the generative model with increasing sample size from 1,000 to 100,000. The other parameters are fixed to M=3 views, G=3 groups, D=1,000 features (per view), and K=25 factors. Bar plots show the training time for standard variational inference (VI) and for stochastic variational inference (SV). Colors represent stochastic models trained with different batch sizes (10%, 25% or 50%). Learning rate and forgetting rate hyperparameters were both fixed to 0.5. VI models were fit using a single E5-2680v3 CPU. SVI models were fit using an Nvidia GTX 1080Ti GPU.

**Figure S3.**
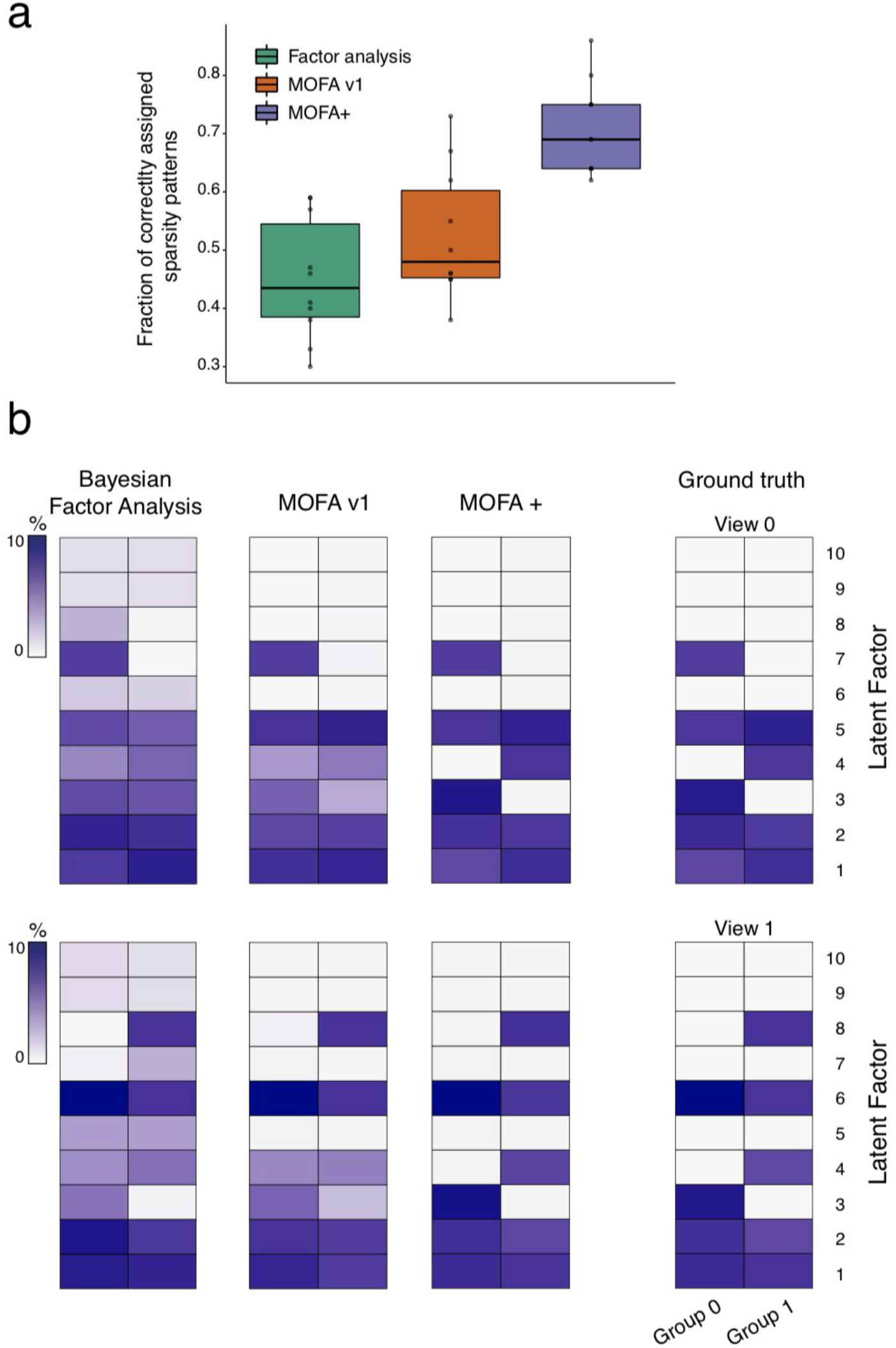
Validation of group-wise ARD prior in the factors using structured simulated data. Data is simulated from the MOFA+ generative model with the following parameters: M=2 views, G=2 groups, D=1,000 features, N=1,000 samples and K=10 factors. We incorporate structure in the simulation process by turning some factors off in random sets of views and groups. The task of MOFA+ is to recover the true factor activity structure given a random initialisation. We compared three models: Bayesian Factor Analysis (no sparsity priors), MOFA v1 (only view-wise sparsity prior) and MOFA+ (view-wise and group-wise sparsity prior). (a) Fraction of correctly assigned sparsity patterns for each model. If the difference between the simulated and the inferred variance explained pattern was less than 0.1%, the pattern was assigned as correct. The box plots show median levels and the first and third quartile out of 10 trials. Whiskers show 1.5x the interquartile range. (b) Representative example of the resulting variance explained patterns. The first row of heatmaps correspond to view 0 and the second row to view 1. In each heatmap, the first column corresponds to group 0 and the second column to group 1. Rows correspond to the inferred factors. The colour scale displays the fraction of variance explained by a given factor in a given view and group. The heatmaps displayed in columns one to three show the solutions yielded by different models (Bayesian Factor Analysis; MOFA; MOFA+). The ground truth is shown in the right panel.

**Figure S4.**
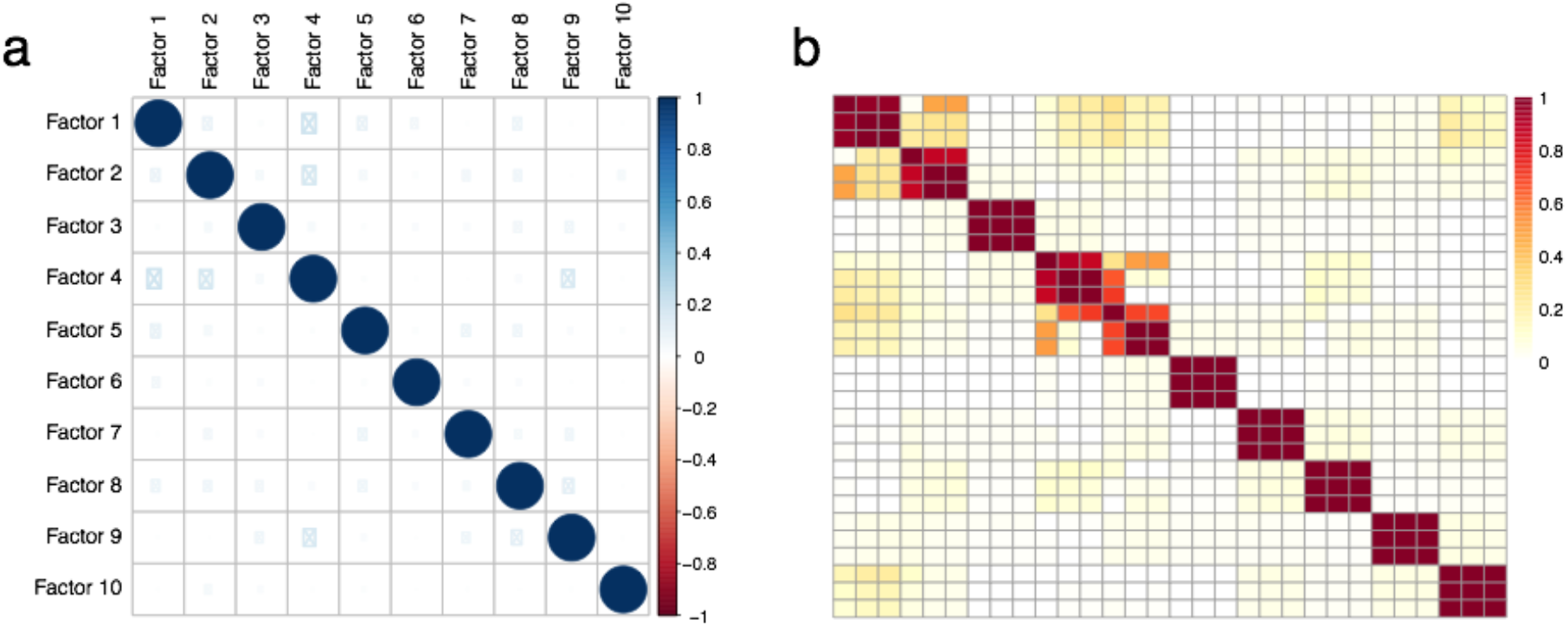
Robustness analysis of MOFA+ factors from the gastrulation scRNA-seq data set. The optimisation procedure of MOFA+ is not guaranteed to find a consistent optimal solution at every trial, and factors can vary between different model instances. To assess the consistency of factors we trained models under different random parameter initialisations and computed the Pearson correlation between factors within and between trials. (a) shows the correlations of factors within a trial. In MOFA there is no orthogonality constraints, but in order to maximise the variance explained factors are expected to be largely uncorrelated. (b) shows a heatmap of the Pearson correlation coefficients between every pair of factors in different trials. Each diagonal block represents a factor that is consistently learnt across multiple trials, suggesting that all 10 factors are robust (i.e. consistently found in all trials).

**Figure S5.**
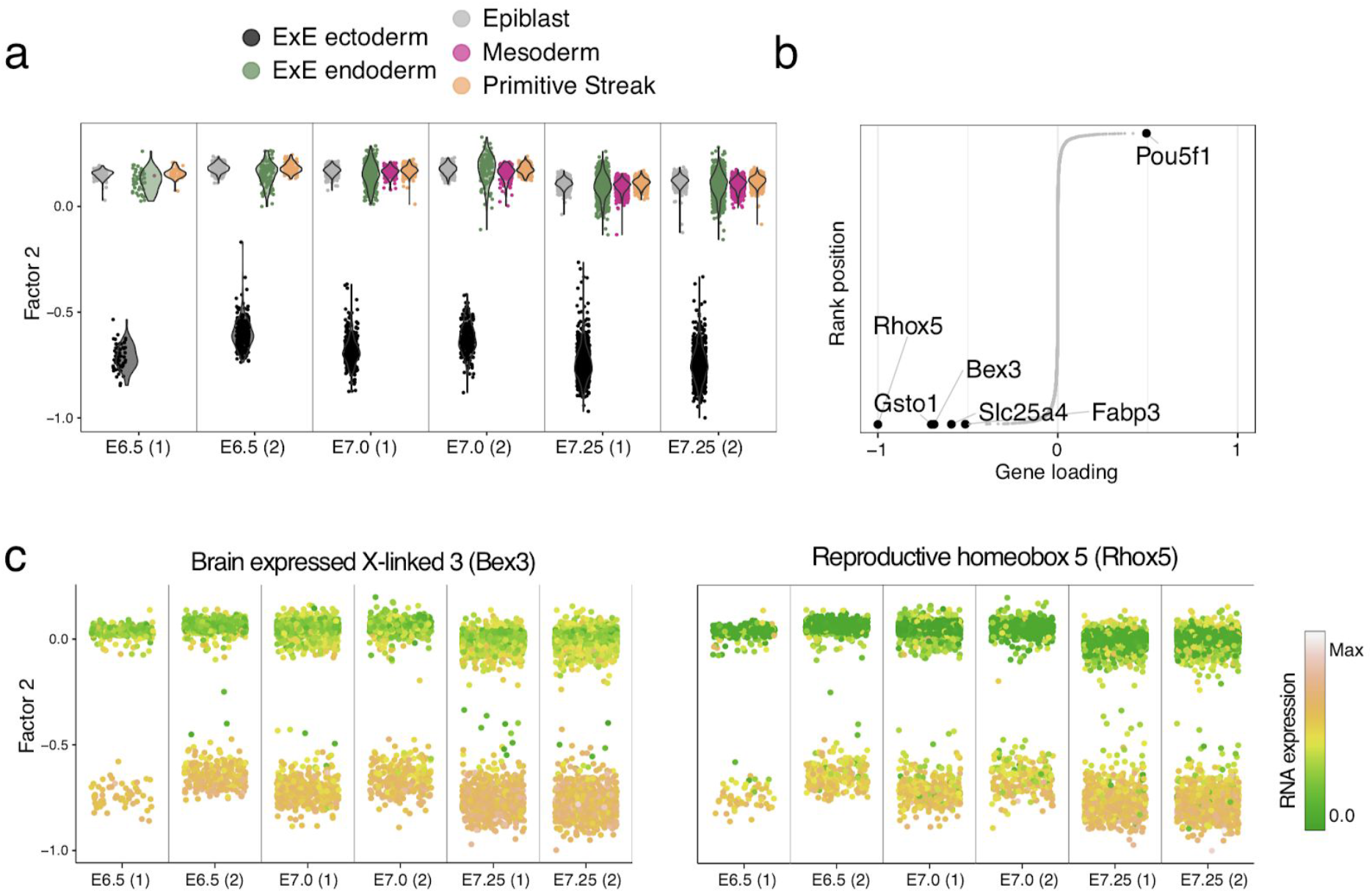
Characterisation of Factor 2 as extra-embryonic (ExE) ectoderm formation. (a) Distribution of factor values per batch of embryos, where each dot represents a single cell, coloured by cell type^1^ (b) Distribution of gene weights, with the top six genes with largest (absolute) weight highlighted. (c) Distribution of factor values per batch of embryos, with cells coloured by the expression of the genes with highest weight.

**Figure S6.**
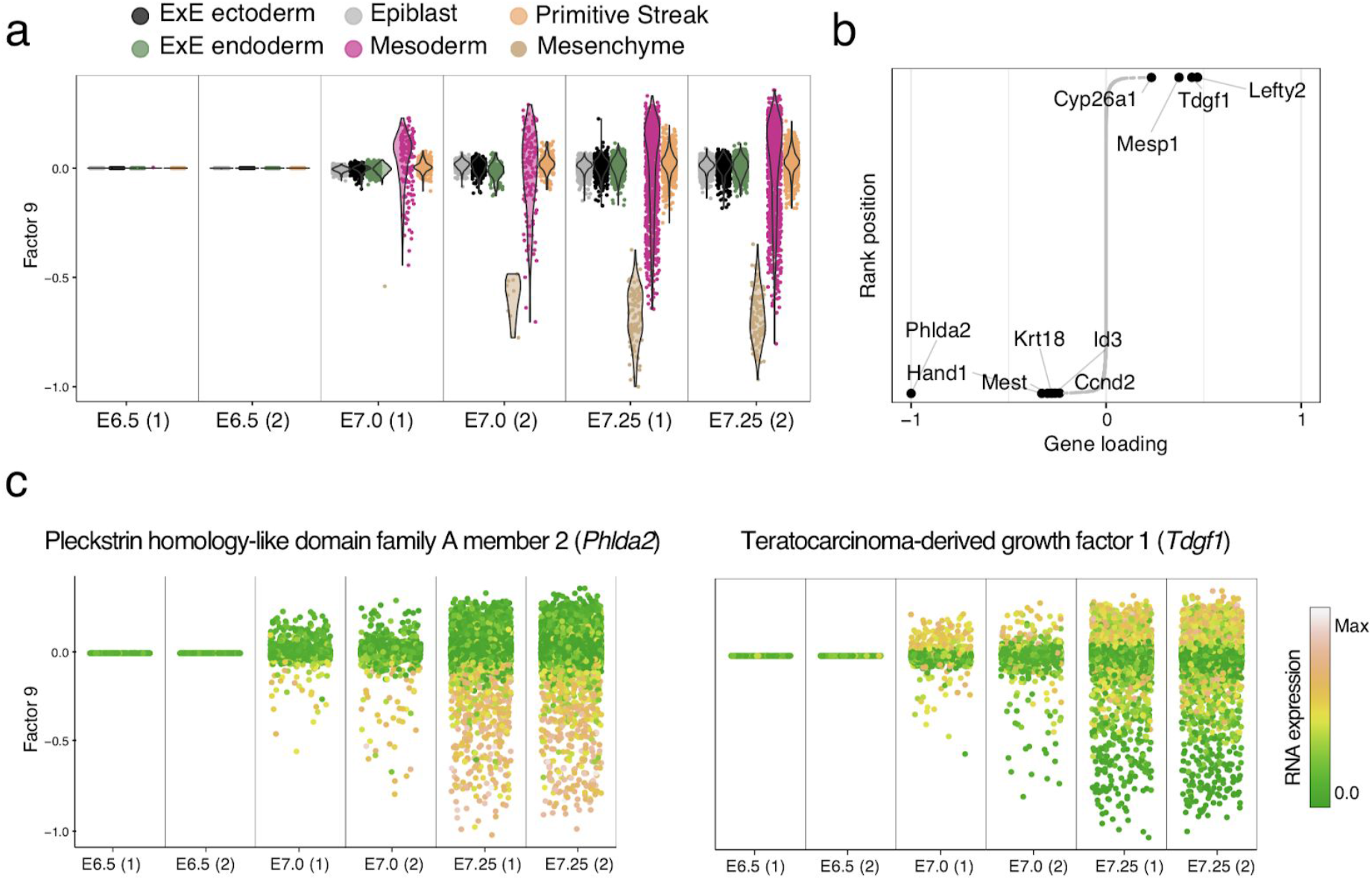
Characterisation of Factor 9 as mesenchyme formation. (a) Distribution of factor values per batch of embryos, where each dot represents a single cell, coloured by cell type^1^. (b) Distribution of gene weights, with the top ten genes with largest (absolute) weight highlighted. (c) Distribution of factor values per batch of embryos, with cells coloured by the expression of the genes with highest weight.

**Figure S7.**
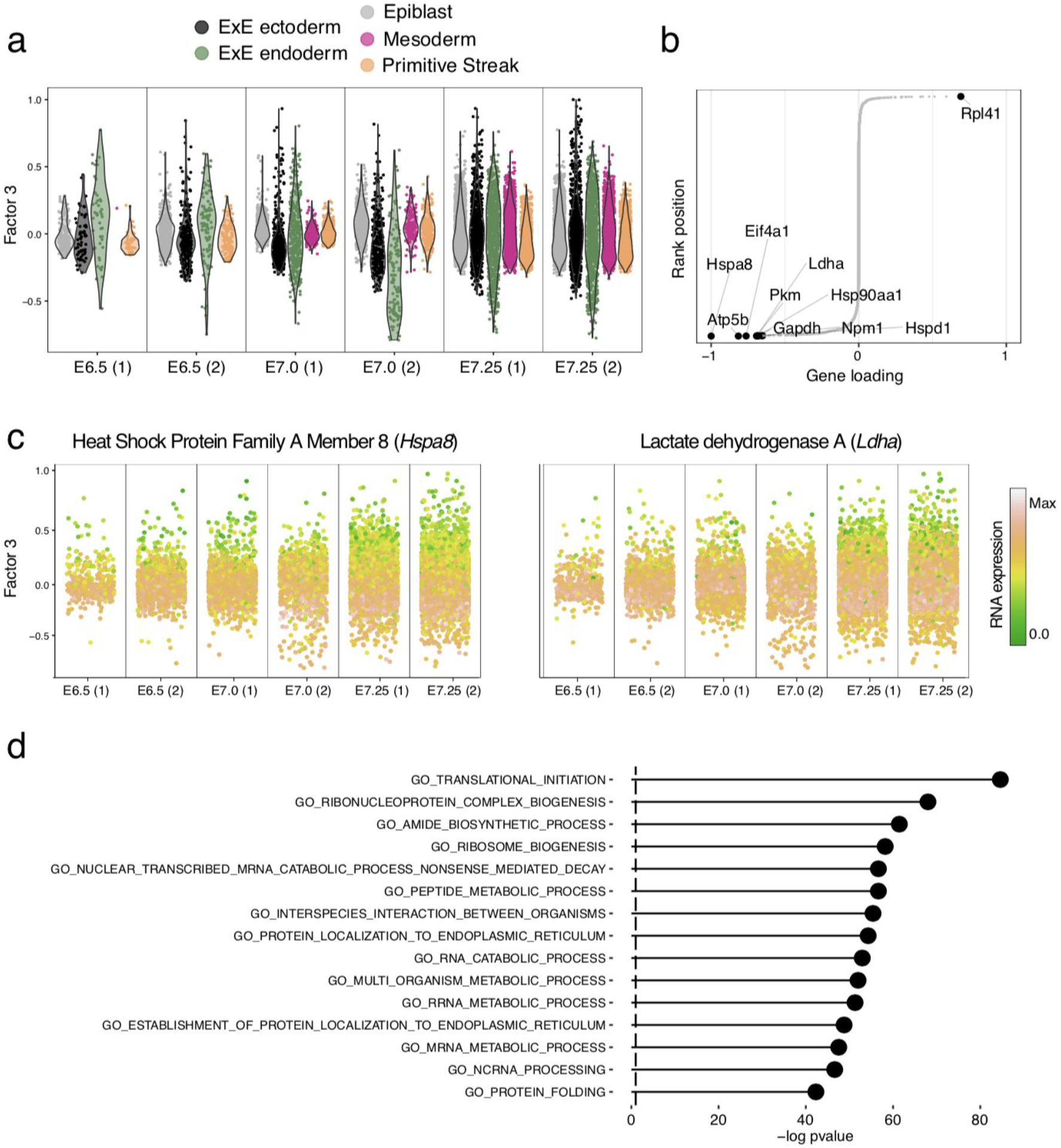
Characterisation of Factor 3 as metabolic stress. (a) Distribution of factor values per batch of embryos, coloured by cell type^1^. (b) Distribution of gene weights, with the top six genes with largest (absolute) weight highlighted. (c) Distribution of factor values per batch of embryos, with cells coloured by the expression of the genes with highest weight. (d) Gene set enrichment analysis applied to the Factor 3 weights (Methods)

**Figure S8:**
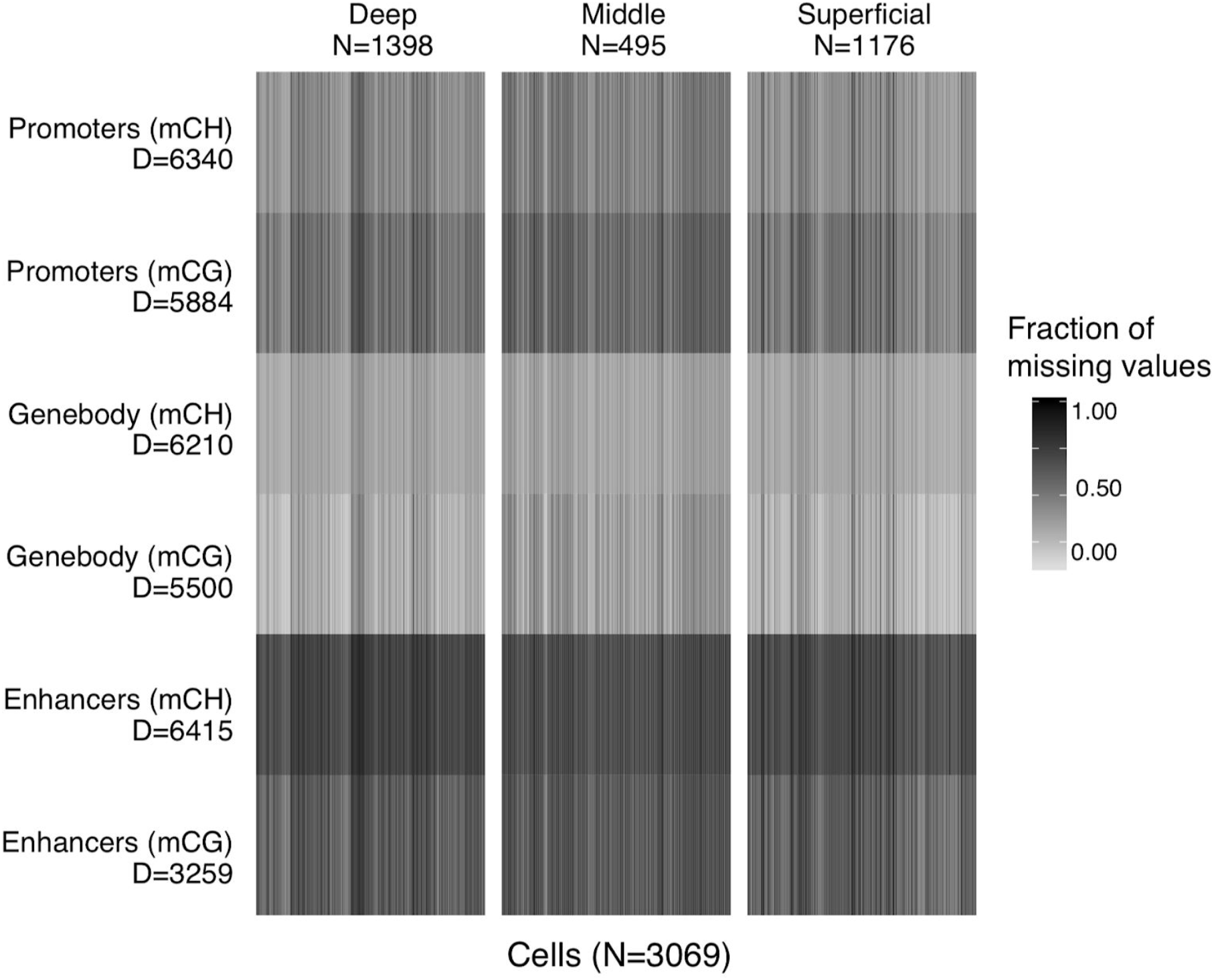
Overview of the single-cell DNA methylation data set. The tile plot shows the structure of the input data in terms of views (rows) versus groups (columns), with associated dimensionalities (D for features, N for samples). The color displays the fraction of missing values for each combination of sample and view.

**Figure S9.**
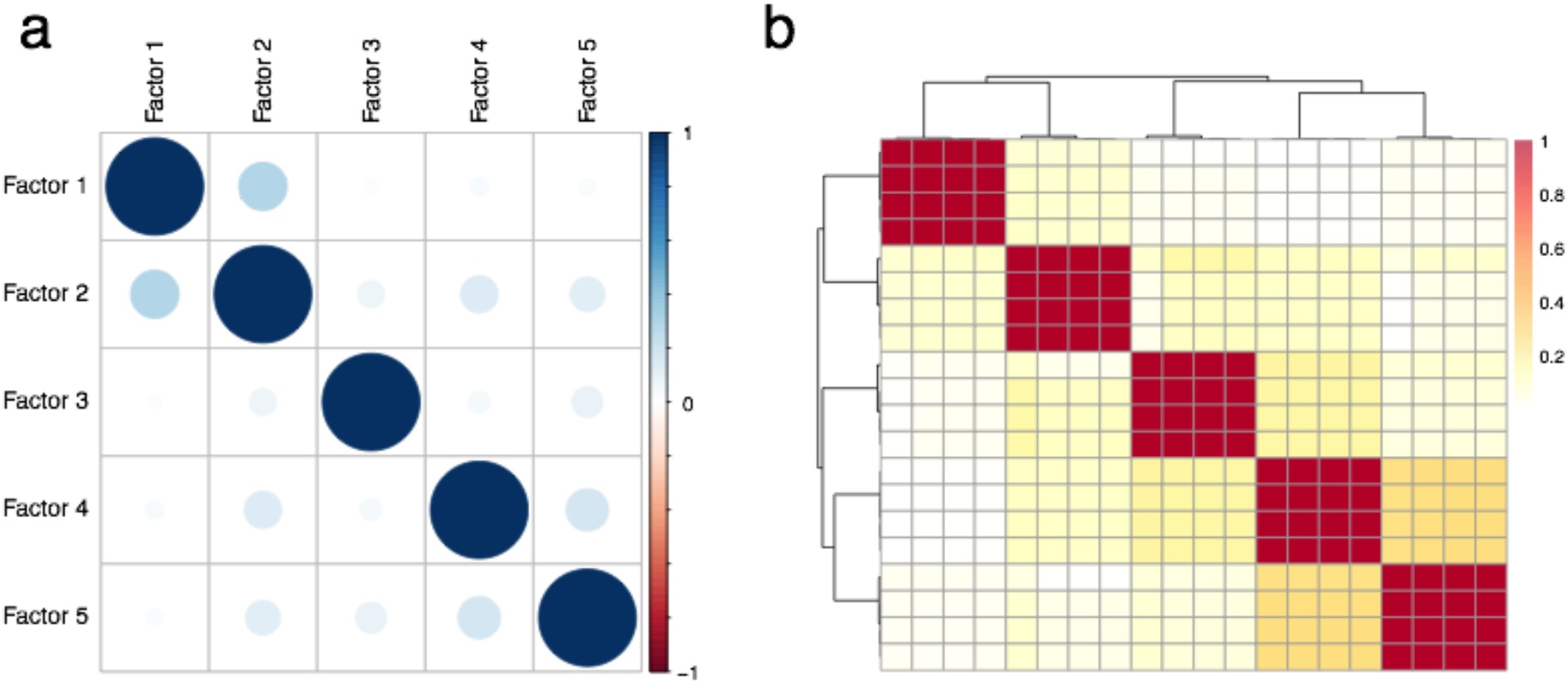
Robustness of MOFA factors from the single-cell DNA methylation data set. The optimisation procedure of MOFA is not guaranteed to find a consistent optimal solution at every trial, and factors can vary between different model instances. To assess the consistency of factors we trained models under different random parameter initialisations and computed the Pearson correlation between factors within and between trials. (a) shows the correlations of factors within a trial. In MOFA there is no orthogonality constraints, but in order to maximise the variance explained factors are expected to be largely uncorrelated. (b) shows a heatmap of the Pearson correlation coefficients between every pair of factors in different trials. Each diagonal block represents a factor that is consistently learnt across multiple trials, suggesting that all 5 factors are robust (i.e. consistently found in all trials).

**Figure S10:**
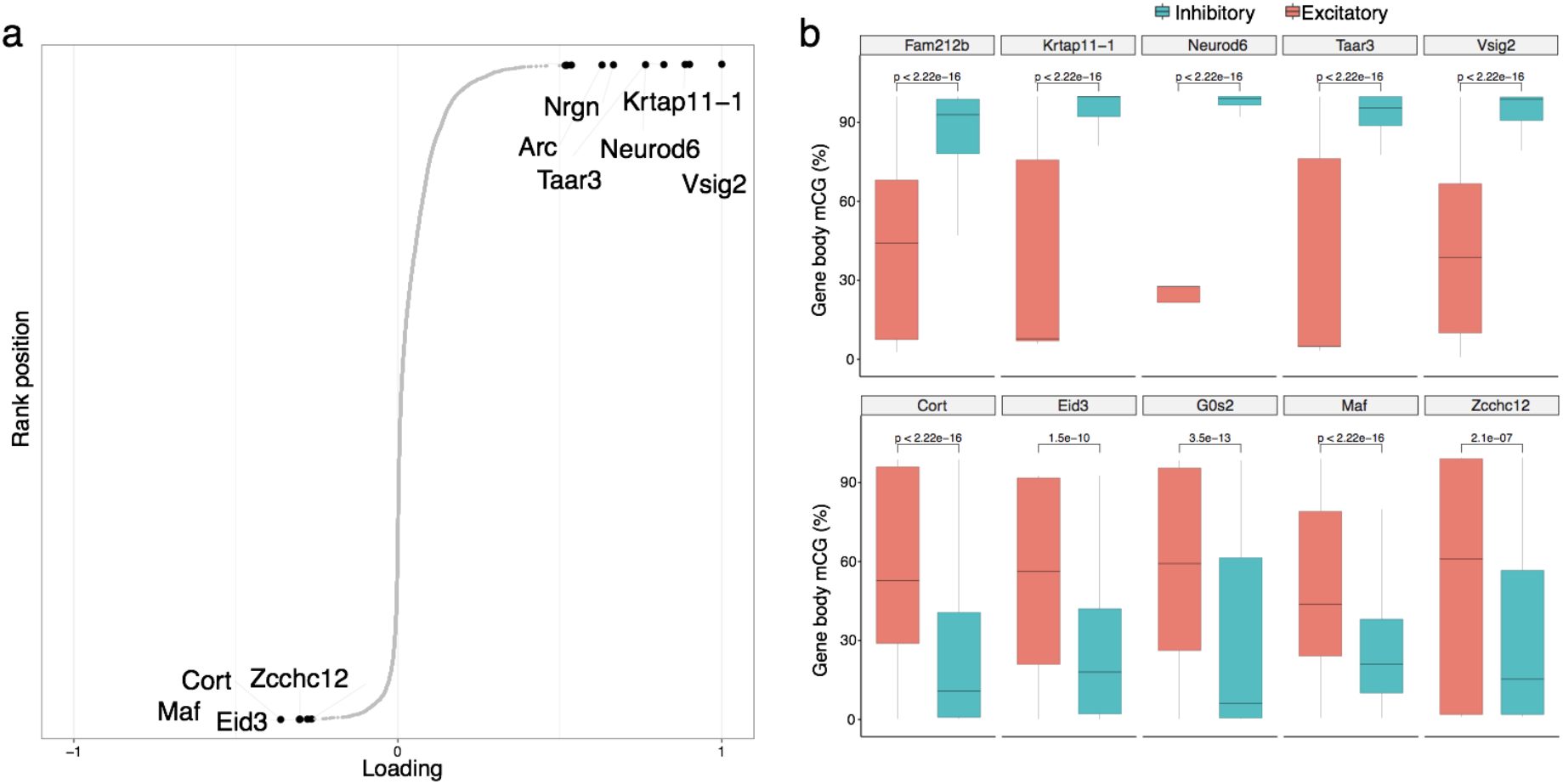
Inspection of gene body mCG weights for Factor 1. (a) Cumulative distribution of gene body mCG weights (x-axis) for Factor 1. Each dot corresponds to one gene, sorted by rank (y-axis) The weights provide a measure of feature importance, hence the higher the weight in absolute value, the higher the association between the feature and the factor (in this case, excitatory vs inhibitory neurons). The sign of the weight indicates the direction of the variability; positive weights indicate higher mCG in cells with positive Factor 1 values (inhibitory cells, see Figure 3b), whereas negative weights indicate lower mCG in cells with negative Factor 1 values (excitatory cells, see Figure 3b). (b) Box plots comparing the distribution of gene body mCG levels (%) between excitatory and inhibitory neurons for the top 5 genes with the highest positive (top) or negative (bottom) weight. For each gene, a nominal p-value is reported using a t-test.

**Figure S11:**
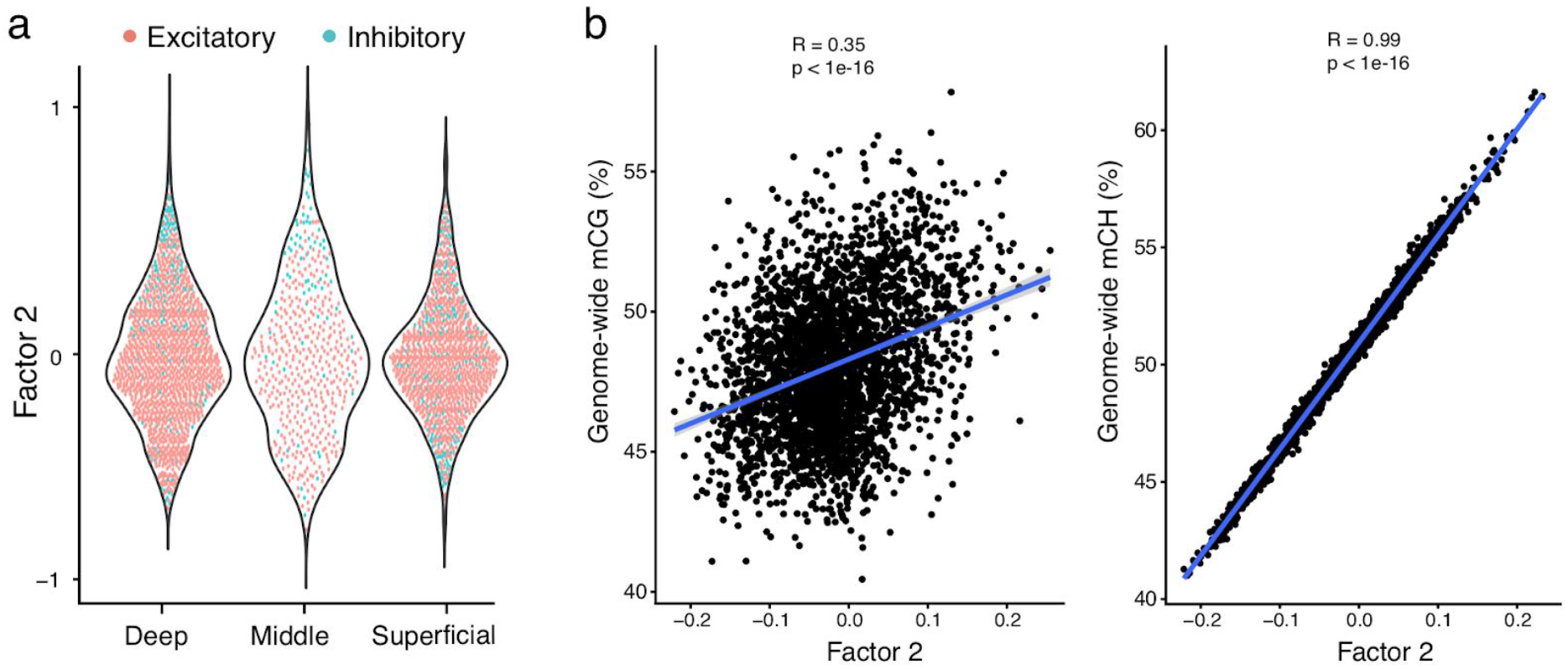
Characterisation of Factor 2 as differences in global mCH levels. (a) Beeswarm plots show the distribution of Factor 2 values for each cortical layer. Cells are coloured by neuron class. (b) Correlation of Factor 2 values (x-axis) with global mCG levels (%, left) and global mCH levels (%, right). The blue line shows the linear regression fit.

**Figure S12.**
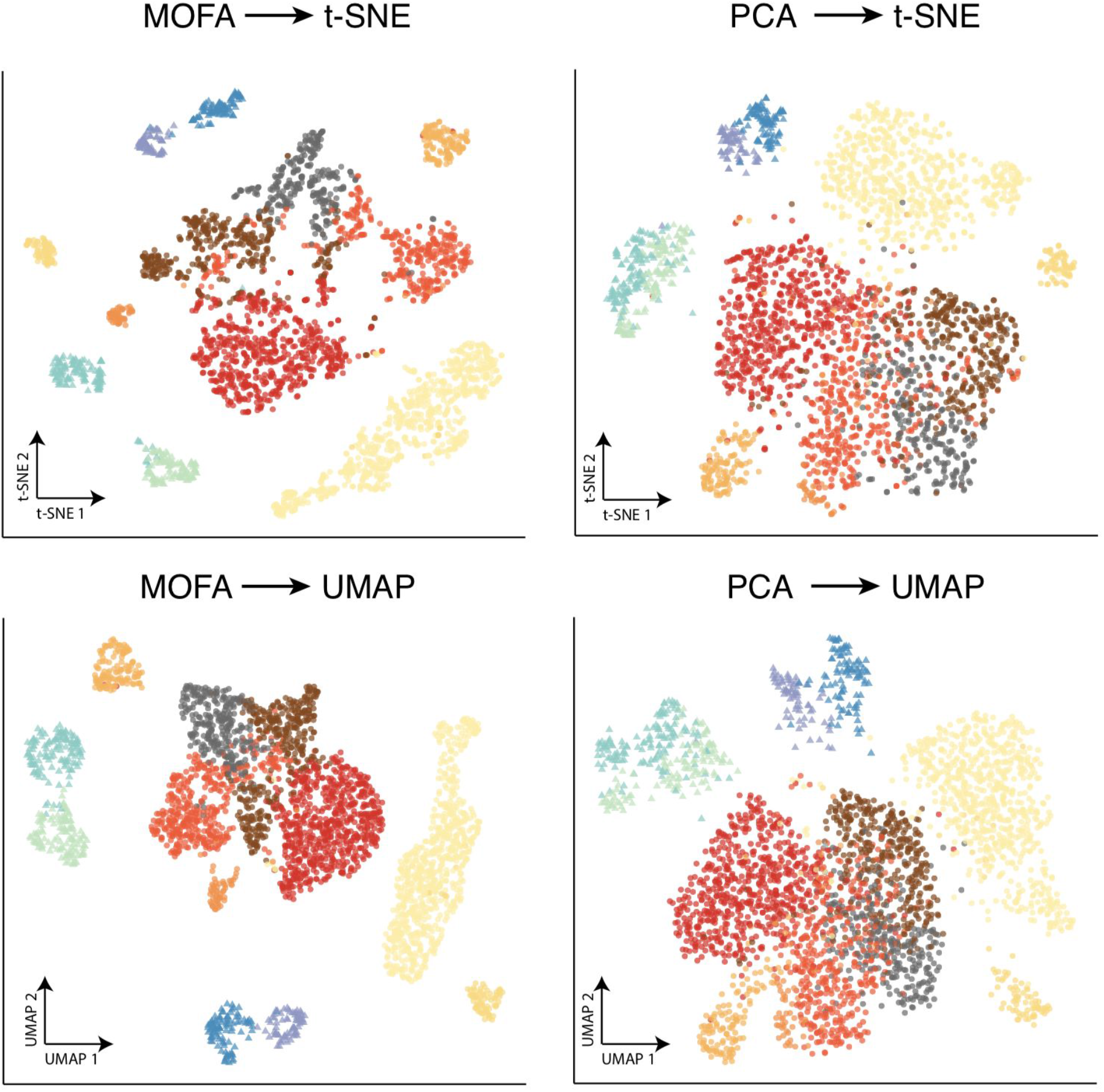
Benchmarking MOFA+ factors and Principal Components as input to non-linear dimensionality reduction methods for DNA methylation data. Plots display UMAP (bottom) or t-SNE (top) projections when using as input MOFA+ factors (left) or principal components (right). Each dot represents a cell, coloured by cell type assignments^2^. Conventional implementations of Principal Component Analysis *(irlba* R package) do not handle missing values; missing values are thus imputed using feature-wise means. To ensure a fair comparison we used the same number of PCs and MOFA+ Factors (K=15)

**Figure S13.**
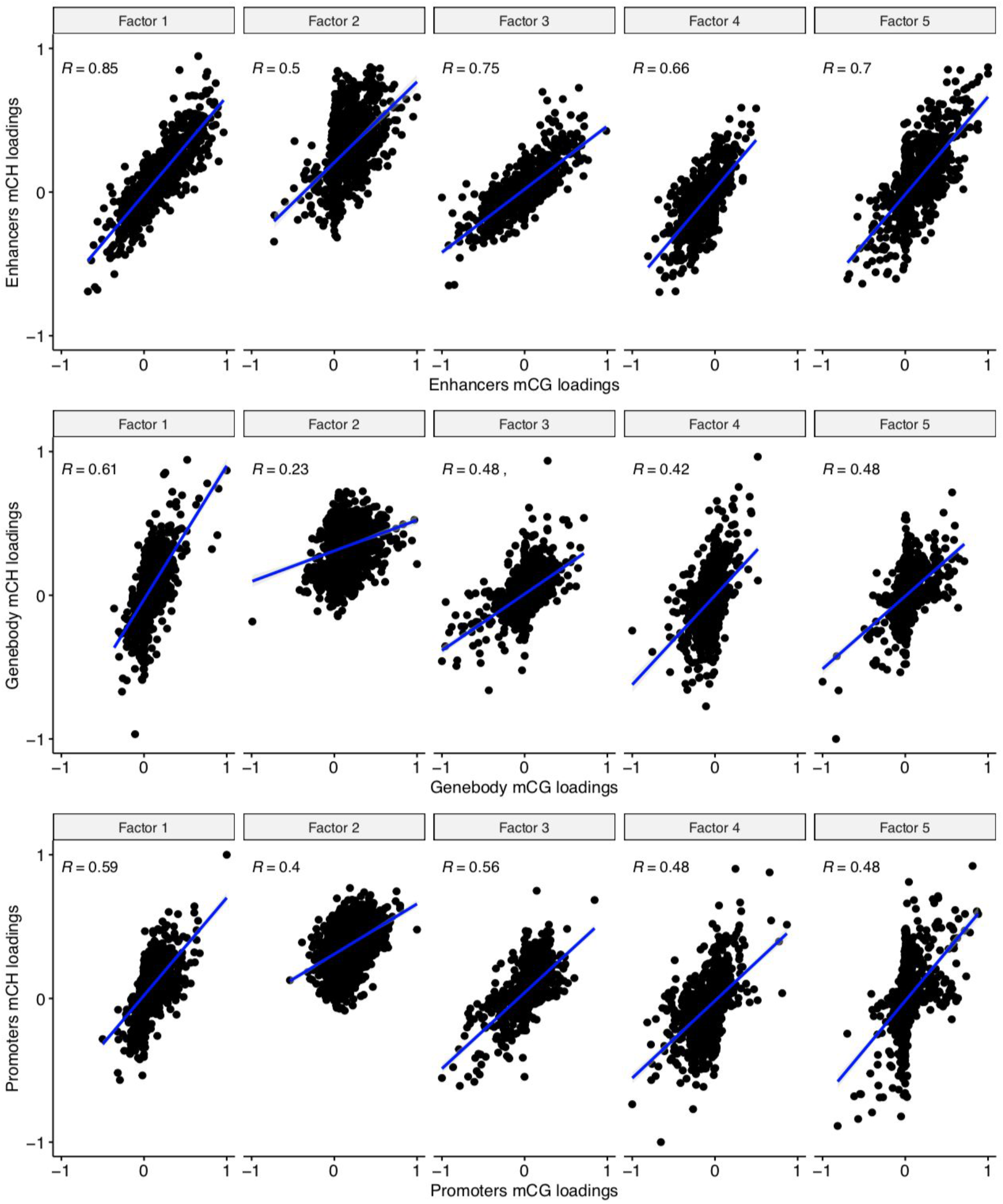
mCH signatures are redundant to mCG signatures across multiple genomic contexts. Scatter plots show the correlation between mCG weights (x-axis) and mCH weights (y-axis) for all combinations of factors (columns) and genomic contexts (rows). The blue lines display linear regression fits (all p-values<10^-16^). For each case we observe a significant positive correlation, indicating that the two DNA methylation signatures are not independent.

**Figure S14.**
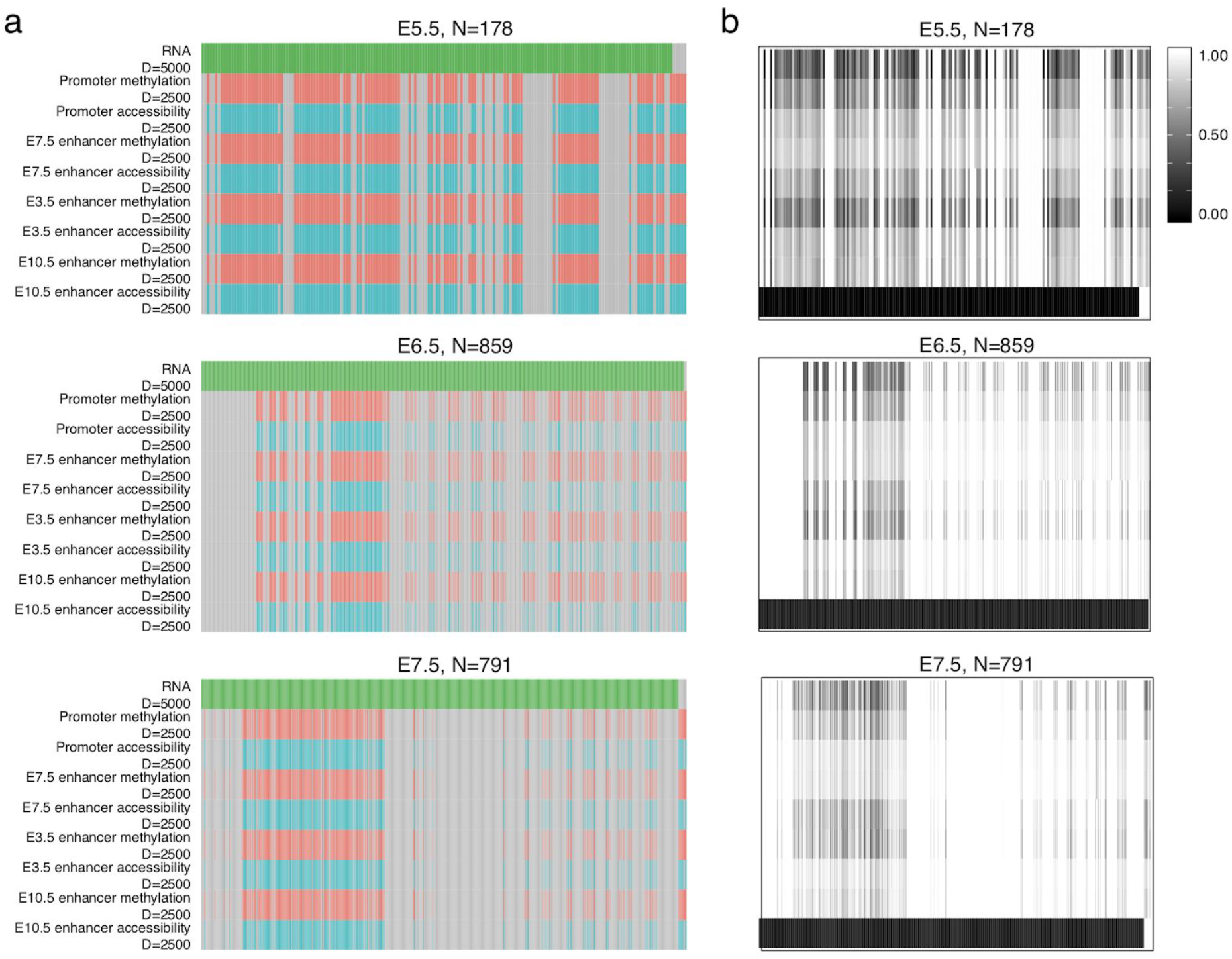
Overview of multi-omic atlas of mouse gastrulation generated using scNMT-seq. (a) Structure of the input data in terms of views (x-axis) versus samples (y-axis). Each panel corresponds to a different group (embryonic stage). Grey bars represent missing views. (b) Structure of the missing values in the data. For each cell and view, the colour displays the fraction of missing values.

**Figure S15:**
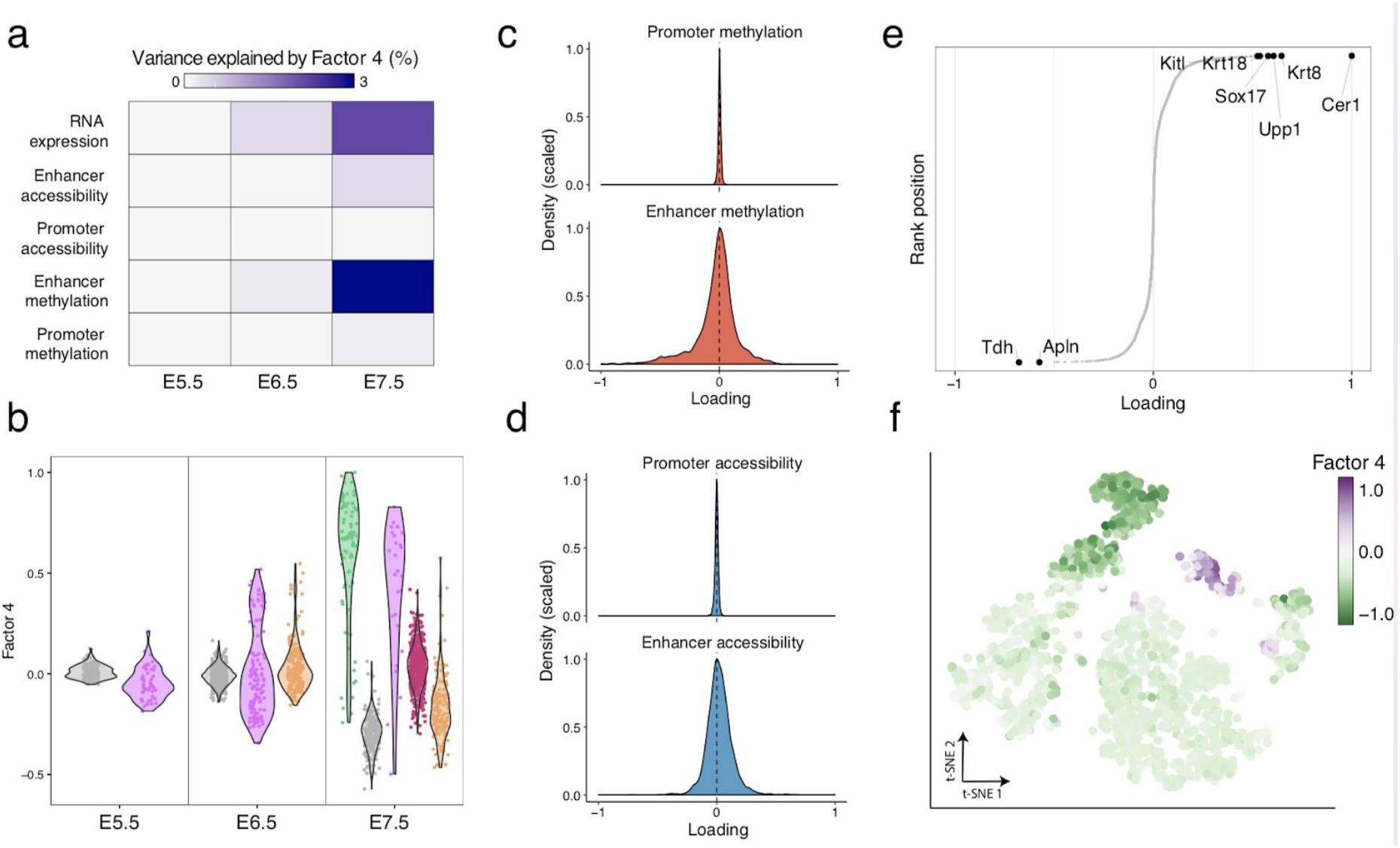
Characterisation of Factor 4 as Embryonic endoderm commitment. (a) Variance explained by Factor 4 in each group (embryonic stage, as columns) and view (rows). (b) Distribution of Factor 4 values per group (embryonic stage, x-axis), with cells coloured by cell type assignment^3^. (c-d) Histograms display the distribution of (c) DNA methylation and (d) chromatin accessibility weights for promoters and enhancer elements. (e) Distribution of RNA weights. The top genes with the highest (absolute) weight are labeled. (f) Dimensionality reduction using t-SNE on the inferred MOFA factors. Cells are coloured by Factor 4 values.

**Figure S16:**
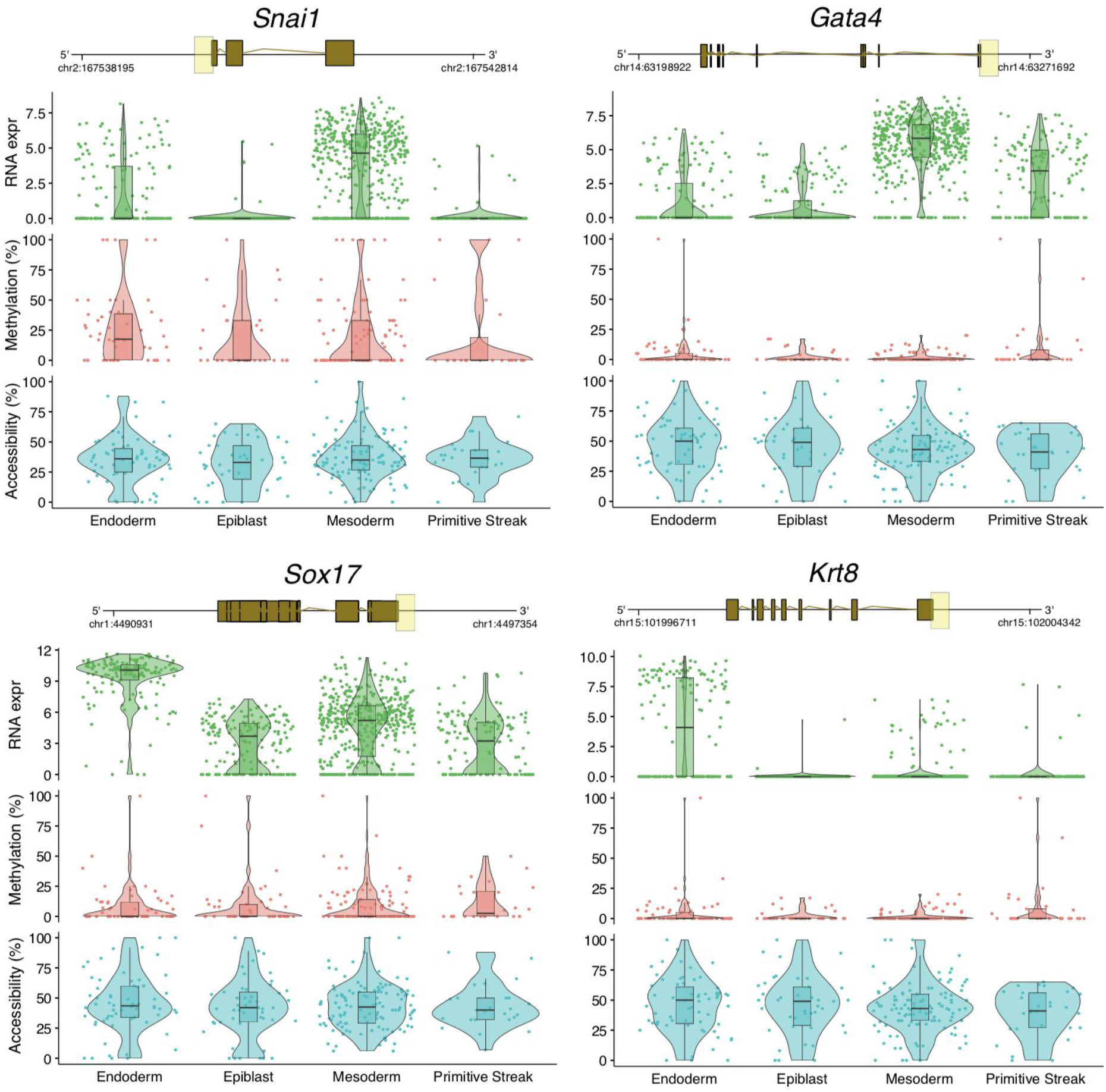
Promoters of genes that display significant differential RNA expression between germ layers do not show differential epigenetic dynamics. Box and violin plots show the distribution of RNA expression (log2 counts, green), DNA methylation (%, red) and chromatin accessibility (%, blue) levels per germ layer at E7.5. Each dot corresponds to a single cell. For each gene a genomic track is shown on the top. The promoter region that is used to quantify DNA methylation and chromatin accessibility levels (2kb upstream and downstream of the TSS) is highlighted in yellow.

**Figure S17:**
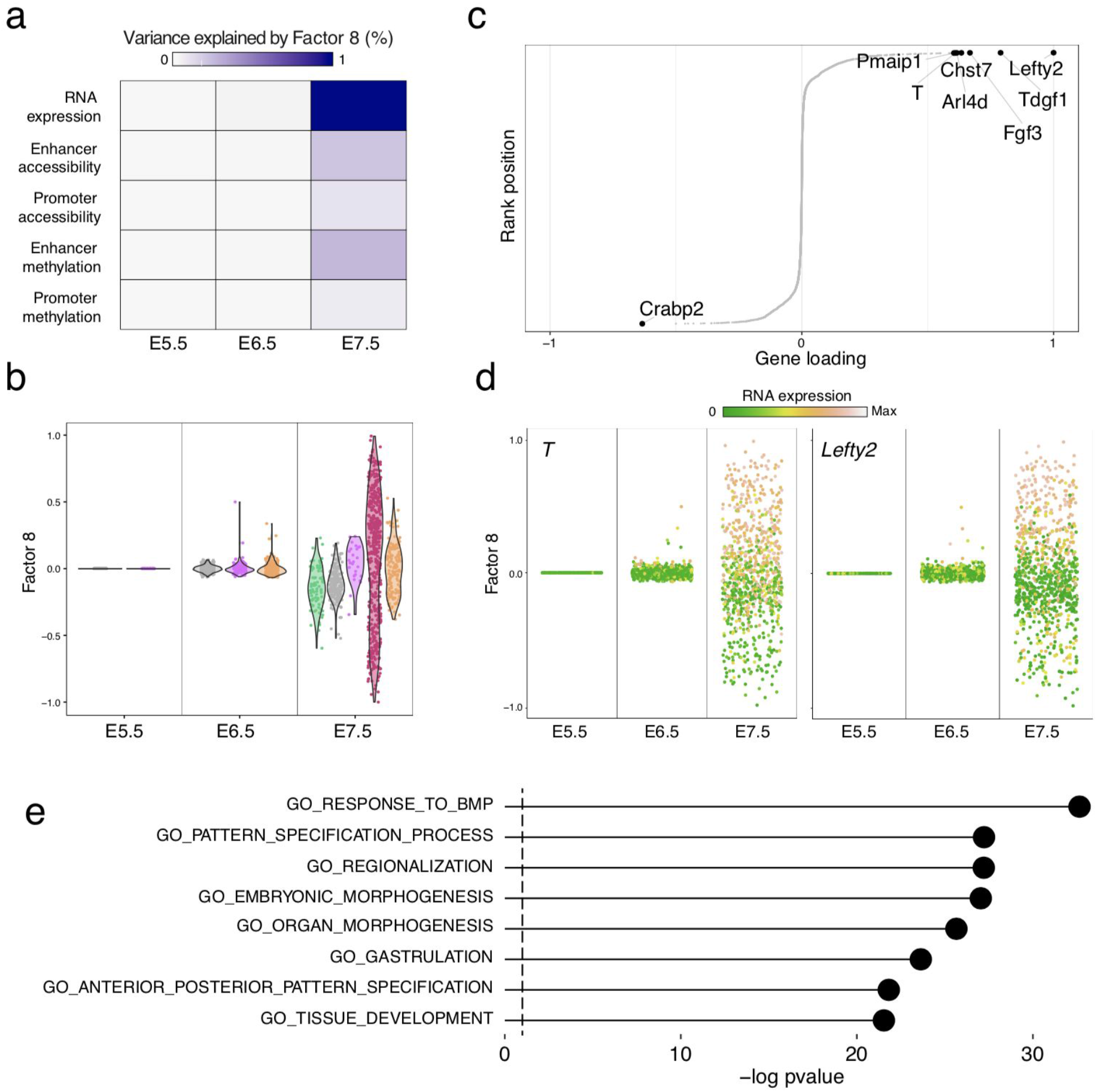
Characterisation of Factor 8 as anterior-posterior axis formation in response to signalling cues. (a) Variance explained by Factor 8 in each group (embryonic stage, columns) and view (rows). (b) Distribution of Factor 8 values per group (embryonic stage, x-axis) with cells coloured by cell type. (c) Distribution of RNA weights for Factor 8. The top genes with the highest (absolute) weight are labeled. (d) Distribution of Factor 8 values per group (embryonic stage, x-axis), with cells coloured by the expression of *T* (left) and *Lefty2* (right) (e) Gene set enrichment analysis applied to the Factor 8 weights.

**Figure S18:**
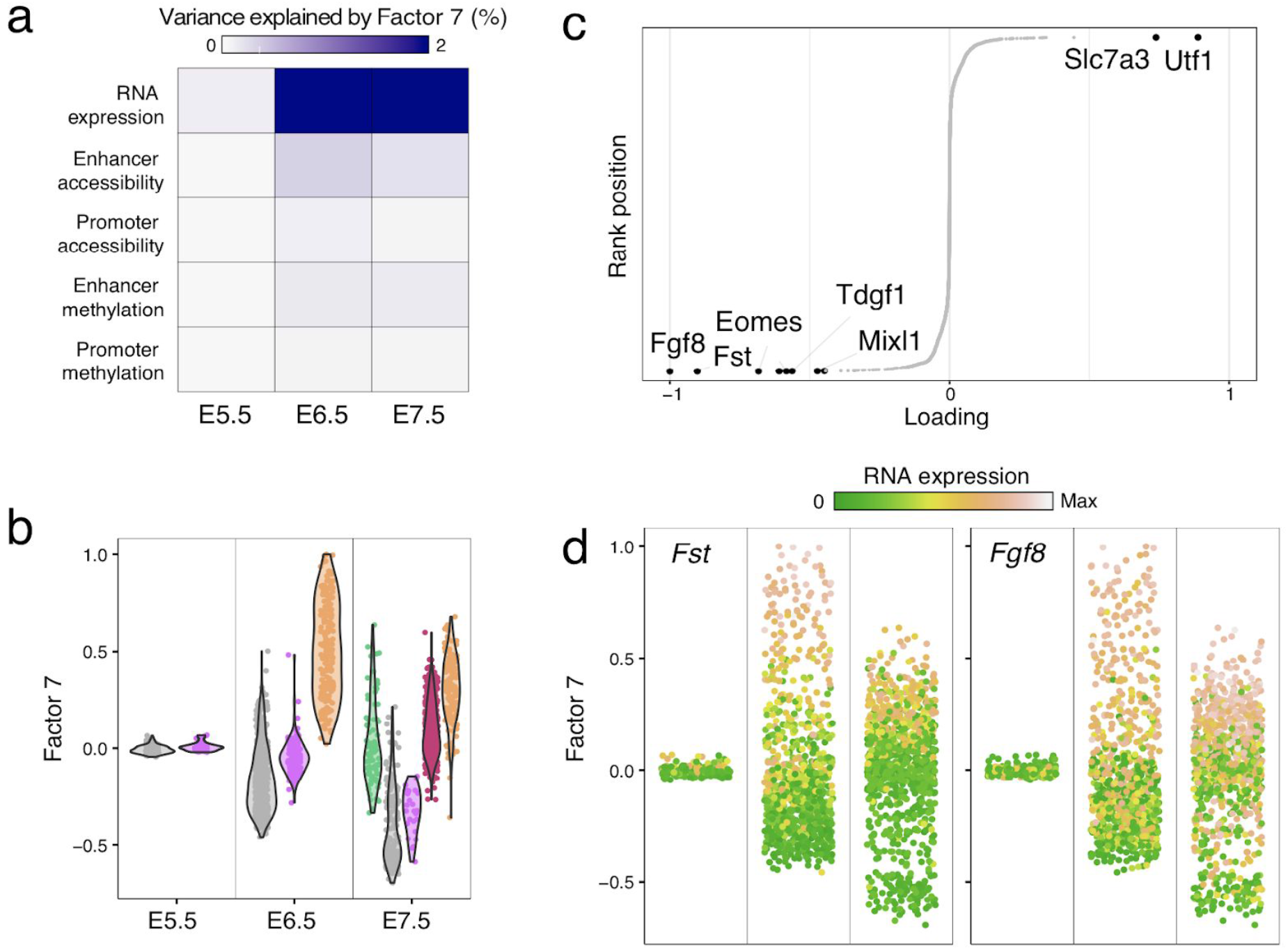
Characterisation of Factor 7 as Primitive Streak formation. (a) Variance explained by Factor 7 in each group (embryonic stage, columns) and view (rows). (b) Distribution of Factor 7 values per group (embryonic stage, x-axis), with cells coloured by cell type. (c) Distribution of RNA weights for Factor 7. The top genes with the highest (absolute) weight are labeled. (c) Distribution of Factor 7 values per stage, with cells coloured by the expression of *Fst* (left) and *Fgf8* (right)

**Figure S19:**
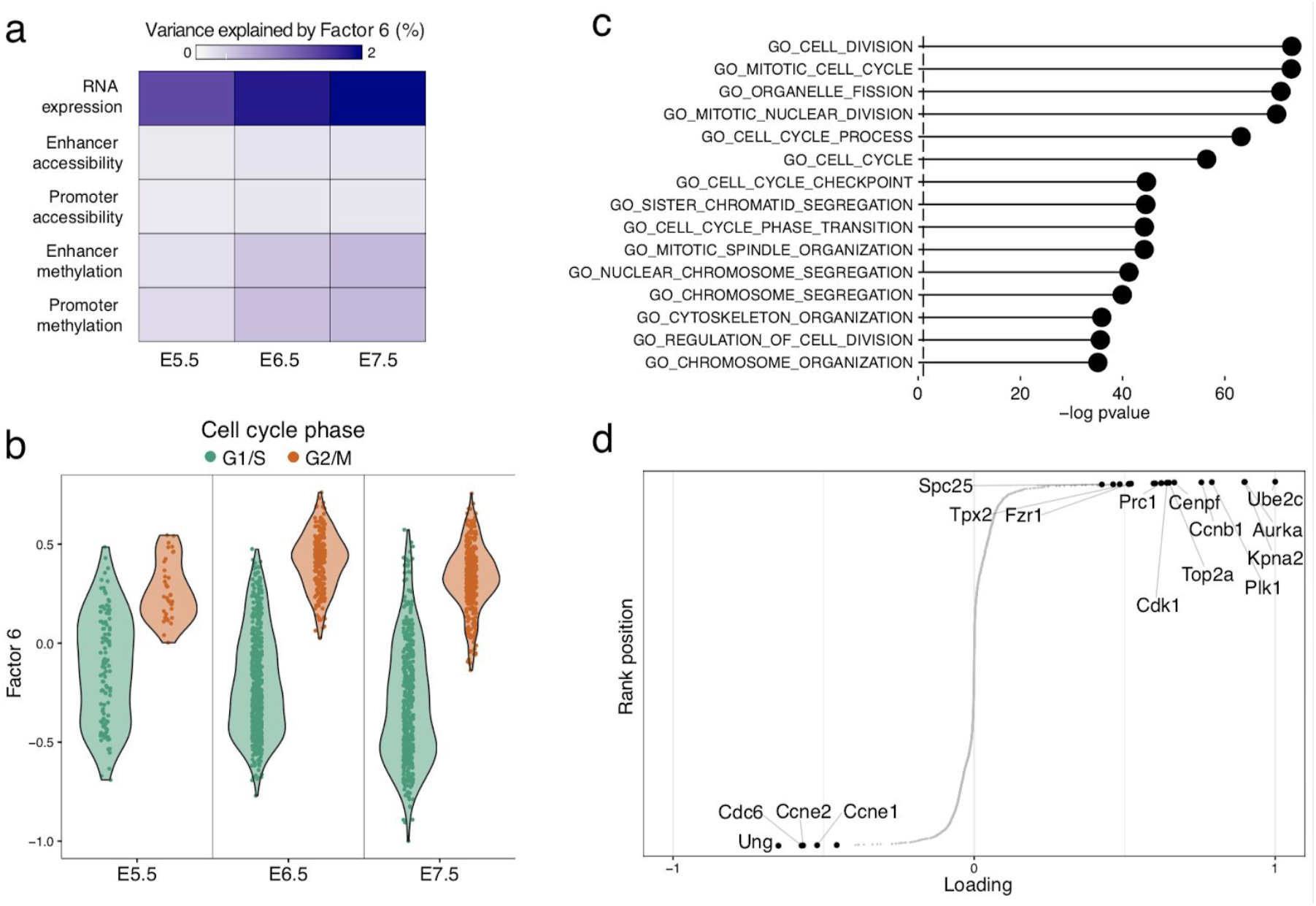
Characterisation of Factor 6 as Cell Cycle. (a) Variance explained by Factor 6 in each group (embryonic stage, columns) and view (rows). (b) Distribution of Factor 6 values per group (embryonic stage, x-axis), with cells coloured by the inferred cell cycle state using *cyclone*^4^. (c) Gene set enrichment analysis applied to the Factor 6 weights. (d) Cumulative distribution of RNA weights for Factor 6. The top genes with the highest (absolute) weight are labeled.

**Figure S20:**
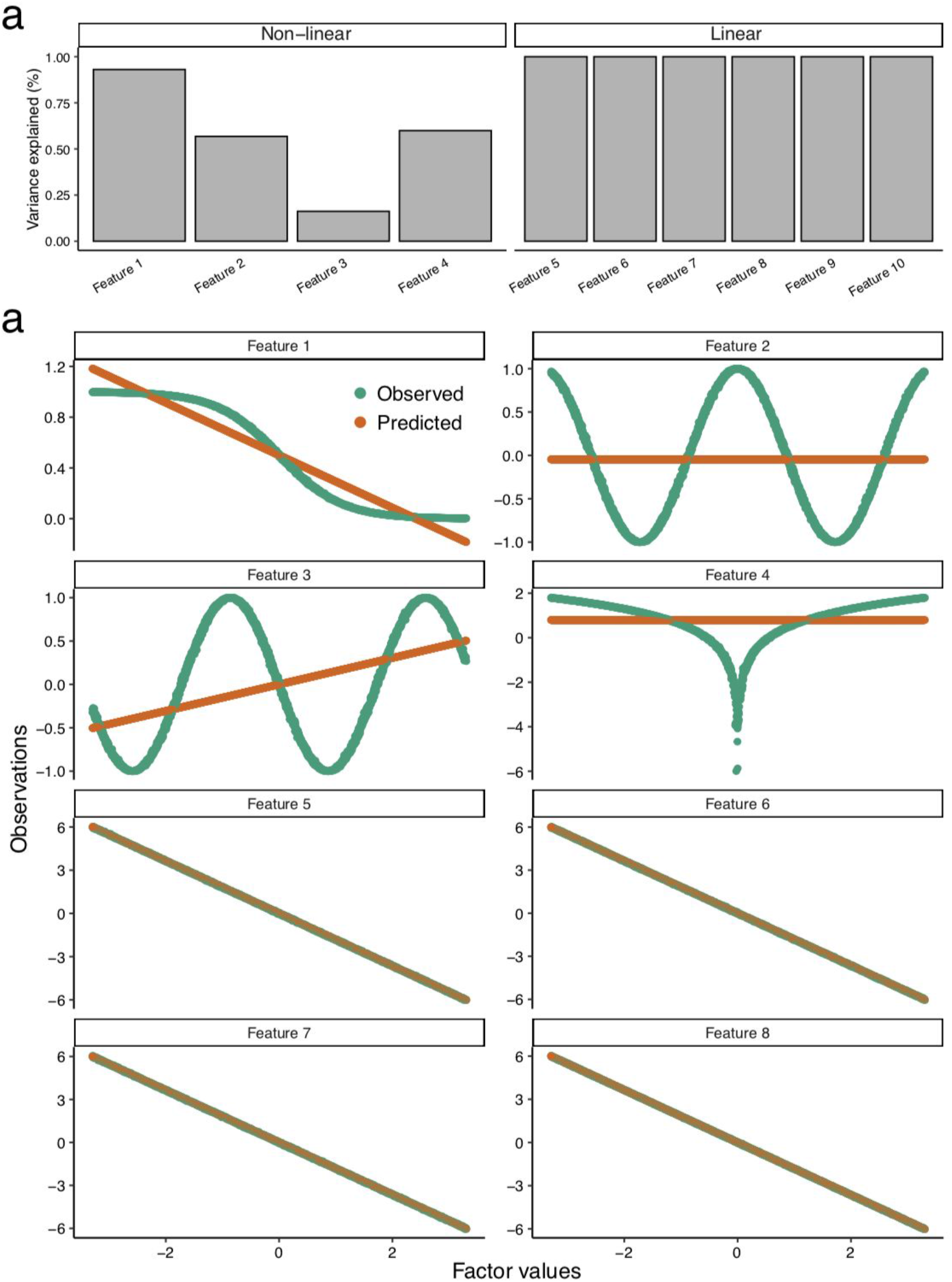
MOFA has limited ability to detect strong non-linear relationships. Data is simulated from the (linear) generative model with the following dimensionalities: M=1 views, G=1 groups, D=8 features, N=1,000 samples and K=1 factors. Data was simulated without noise, but for the first four features we introduced different classes of non-linearities: sigmoid(x), cos(x), sin(x) and log(abs(x)) functions, respectively. (a) Fraction of variance explained per feature (b) Predicted (orange) and observed (green) measurements versus Factor values.

**Figure S21:**
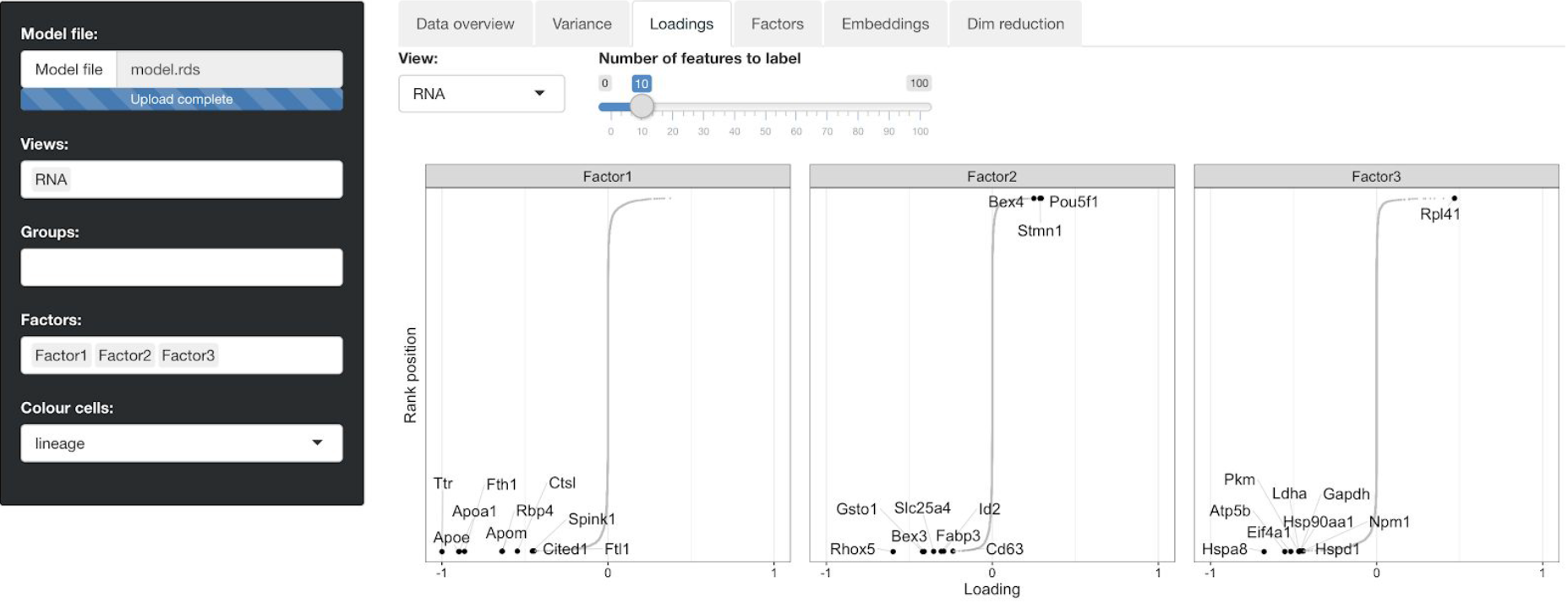
Screenshot of the interactive web-based platform to explore MOFA+ models. The platform is implemented in shiny R from Rstudio and it is available in https://github.com/gtca/mofaplus-shinv

## References

1. Griffiths, J. A., Scialdone, A. & Marioni, J. C. Using single-cell genomics to understand developmental processes and cell fate decisions. Mol. Syst. Biol. 14, e8046 (2018).

2. Papalexi, E. & Satija, R. Single-cell RNA sequencing to explore immune cell heterogeneity. Nat. Rev. Immunol. 18, 35–45 (2018).

3. Wills, Q. F. & Mead, A. J. Application of single-cell genomics in cancer: promise and challenges. Hum. Mol. Genet. 24, R74–84 (2015).

4. Patel, A. P. et al. Single-cell RNA-seq highlights intratumoral heterogeneity in primary glioblastoma. Science 344, 1396–1401 (2014).

5. Mulqueen, R. M. et al. Highly scalable generation of DNA methylation profiles in single cells. Nat. Biotechnol. 36, 428–431 (2018).

6. Guo, H. et al. Single-cell methylome landscapes of mouse embryonic stem cells and *early embryos analyzed using reduced representation bisulfite sequencing*. Genome Res. 23, 2126–2135 (2013).

7. Luo, C. et al. Single-cell methylomes identify neuronal subtypes and regulatory elements in mammalian cortex. Science 357, 600–604 (2017).

8. Clark, S. J. et al. Genome-wide base-resolution mapping of DNA methylation in single cells using single-cell bisulfite sequencing (scBS-seq). Nat. Protoc. 12, 534–547 (2017).

9. Smallwood, S. A. et al. Single-cell genome-wide bisulfite sequencing for assessing epigenetic heterogeneity. Nat. Methods 11, 817–820 (2014).

10. Buenrostro, J. D. et al. Single-cell chromatin accessibility reveals principles of regulatory variation. Nature 523, 486–490 (2015).

11. Mezger, A. et al. High-throughput chromatin accessibility profiling at single-cell resolution. Nat. Commun. 9, 3647 (2018).

12. Macaulay, I. C., Ponting, C. P. & Voet, T. Single-Cell Multiomics: Multiple Measurements from Single Cells. Trends Genet. 33, 155–168 (2017).

13. Bock, C., Farlik, M. & Sheffield, N. C. Multi-Omics of Single Cells: Strategies and Applications. Trends Biotechnol. 34, 605–608 (2016).

14. Macaulay, I. C. et al. G&T-seq: parallel sequencing of single-cell genomes and transcriptomes. Nat. Methods 12, 519–522 (2015).

15. Angermueller, C. et al. Parallel single-cell sequencing links transcriptional and epigenetic heterogeneity. Nat. Methods 13, 229–232 (2016).

16. Cao, J. et al. Joint profiling of chromatin accessibility and gene expression in thousands of single cells. Science 361, 1380–1385 (2018).

17. Clark, S. J. et al. scNMT-seq enables joint profiling of chromatin accessibility DNA methylation and transcription in single cells. Nat. Commun. 9, 781 (2018).

18. Li, L. et al. Single-cell multi-omics sequencing of human early embryos. Nat. Cell Biol. 20, 847–858 (2018).

19. Dey, S. S., Kester, L., Spanjaard, B., Bienko, M. & van Oudenaarden, A. Integrated genome and transcriptome sequencing of the same cell. Nat. Biotechnol. 33, 285–289 (2015).

20. Guo, F. et al. Single-cell multi-omics sequencing of mouse early embryos and embryonic stem cells. Cell Res. 27, 967–988 (2017).

21. Pott, S. Simultaneous measurement of chromatin accessibility, DNA methylation, and nucleosome phasing in single cells. Elife 6, (2017).

22. Cheow, L. F. et al. Single-cell multimodal profiling reveals cellular epigenetic heterogeneity. Nat. Methods 13, 833–836 (2016).

23. Bian, S. et al. Single-cell multiomics sequencing and analyses of human colorectal cancer. Science 362, 1060–1063 (2018).

24. Stoeckius, M. et al. Simultaneous epitope and transcriptome measurement in single cells. Nat. Methods 14, 865–868 (2017).

25. Stuart, T. & Satija, R. Integrative single-cell analysis. Nat. Rev. Genet. (2019). doi:10.1038/s41576-019-0093-7

26. Stuart, T., Butler, A., Hoffman, P. & Hafemeister, C. Comprehensive integration of single cell data. BioRxiv (2018).

27. Haghverdi, L., Lun, A. T. L., Morgan, M. D. & Marioni, J. C. Batch effects in single-cell *RNA-sequencing data are corrected by matching mutual nearest neighbors*. Nat. Biotechnol. 36, 421–427 (2018).

28. Barkas, N., Petukhov, V., Nikolaeva, D. & Lozinsky, Y. Wiring together large single-cell RNA-seq sample collections. bioRxiv (2018).

29. Zhang, L. & Zhang, S. Learning common and specific patterns from data of multiple interrelated biological scenarios with matrix factorization. bioRxiv (2018).

30. Welch, J. D. et al. Single-Cell Multi-omic Integration Compares and Contrasts Features of Brain Cell Identity. Cell 177, 1873–1887.e17 (2019).

31. Stuart, T. et al. Comprehensive Integration of Single-Cell Data. Cell 177, 1888–1902.e21 (2019).

32. Argelaguet, R. et al. Multi-Omics Factor Analysis-a framework for unsupervised integration of multi-omics data sets. Mol. Syst. Biol. 14, e8124 (2018).

33. Pijuan-Sala, B. et al. A single-cell molecular map of mouse gastrulation and early organogenesis. Nature 566, 490–495 (2019).

34. McInnes, L., Healy, J. & Melville, J. UMAP: Uniform Manifold Approximation and Projection for Dimension Reduction. arXiv [stat.ML] (2018).

35. Maaten, L. van der & Hinton, G. Visualizing Data using t-SNE. J. Mach. Learn. Res. 9, 2579–2605 (2008).

36. He, Y. & Ecker, J. R. Non-CG Methylation in the Human Genome. Annu. Rev. Genomics Hum. Genet. 16, 55–77 (2015).

37. Ramsahoye, B. H. et al. Non-CpG methylation is prevalent in embryonic stem cells and may be mediated by DNA methyltransferase 3a. Proc. Natl. Acad. Sci. U. S. A. 97, 5237–5242 (2000).

38. Lister, R. et al. Human DNA methylomes at base resolution show widespread epigenomic differences. Nature 462, 315–322 (2009).

39. Chen, L. et al. MeCP2 binds to non-CG methylated DNA as neurons mature, influencing *transcription and the timing of onset for Rett syndrome*. Proc. Natl. Acad. Sci. U. S. A. 112, 5509–5514 (2015).

40. Grung, B. & Manne, R. Missing values in principal component analysis. Chemometrics Intellig. Lab. Syst. 42, 125–139 (1998).

41. Argelaguet, R., Mohammed, H., Clark, S. & Stapel, C. Single cell multi-omics profiling reveals a hierarchical epigenetic landscape during mammalian germ layer specification. bioRxiv (2019).

42. Creyghton, M. P. et al. Histone H3K27ac separates active from poised enhancers and predicts developmental state. Proc. Natl. Acad. Sci. U. S. A. 107, 21931–21936 (2010).

43. Calo, E. & Wysocka, J. Modification of enhancer chromatin: what, how, and why? Mol. Cell 49, 825–837 (2013).

44. Zhang, Y. et al. Dynamic epigenomic landscapes during early lineage specification in mouse embryos. Nat. Genet. 50, 96–105 (2018).

45. Daugherty, A. C. et al. Chromatin accessibility dynamics reveal novel functional enhancers in C. elegans. Genome Res. 27, 2096–2107 (2017).

46. Lee, H. J. et al. Developmental enhancers revealed by extensive DNA methylome maps of zebrafish early embryos. Nat. Commun. 6, 6315 (2015).

47. Cusanovich, D. A. et al. The cis-regulatory dynamics of embryonic development at single-cell resolution. Nature 555, 538–542 (2018).

48. Chen, S., Lake, B. B. & Zhang, K. High-throughput sequencing of the transcriptome and chromatin accessibility in the same cell. Nat. Biotechnol. (2019). doi:10.1038/s41587-019-0290-0

49. Chappell, L., Russell, A. J. C. & Voet, T. Single-cell (multi) omics technologies. Annu. Rev. Genomics Hum. Genet. 19, 15–41 (2018).

50. Lopez, R., Regier, J., Cole, M. B., Jordan, M. I. & Yosef, N. Deep generative modeling for single-cell transcriptomics. Nat. Methods 15, 1053–1058 (2018).

51. Grønbech, C. H. et al. scVAE: Variational auto-encoders for single-cell gene expression data. bioRxiv 318295 (2018). doi:10.1101/318295

52. Lotfollahi, M., Wolf, F. A. & Theis, F. J. scGen predicts single-cell perturbation responses. Nat. Methods 16, 715–721 (2019).

53. Delgado, F. M. & Gómez-Vela, F. Computational methods for Gene Regulatory Networks reconstruction and analysis: A review. Artif. Intell. Med. 95, 133–145 (2019).

54. Gao, C., Brown, C. D. & Engelhardt, B. E. A latent factor model with a mixture of sparse and dense factors to model gene expression data with confounding effects. arXiv [stat.AP] (2013).

55. Saul, L. K., Jaakkola, T. & Jordan, M. I. Mean field theory for sigmoid belief networks. J. Artif. Intell. Res. 4, 61–76 (1996).

56. Zhang, C., Butepage, J., Kjellstrom, H. & Mandt, S. Advances in Variational Inference. IEEE Trans. Pattern Anal. Mach. Intell. 41, 2008–2026 (2019).

57. Blei, D. M., Kucukelbir, A. & McAuliffe, J. D. Variational Inference: A Review for Statisticians. arXiv [stat.CO] (2016).

58. Hoffman, M. D. Stochastic Variational Inference. J. Mach. Learn. Res. 14, 1303–1347 (2013).

59. Fabregat, A. et al. The Reactome pathway Knowledgebase. Nucleic Acids Res. 44, D481–7 (2016).

60. Benjamini, Y. & Hochberg, Y. Controlling the False Discovery Rate: A Practical and Powerful Approach to Multiple Testing. J. R. Stat. Soc. Series B Stat. Methodol. 57, 289–300 (1995).

61. Lun, A. T. L., McCarthy, D. J. & Marioni, J. C. A step-by-step workflow for low-level analysis of single-cell RNA-seq data with Bioconductor. F1000Res. 5, 2122 (2016).

62. Du, P. et al. Comparison of Beta-value and M-value methods for quantifying methylation levels by microarray analysis. BMC Bioinformatics 11, 587 (2010).

63. Liao, Y., Smyth, G. K. & Shi, W. featureCounts: an efficient general purpose program for assigning sequence reads to genomic features. Bioinformatics 30, 923–930 (2014).

64. Yates, A. et al. Ensembl 2016. Nucleic Acids Res. 44, D710–6 (2016).

## References

1. Pijuan-Sala, B. et al. A single-cell molecular map of mouse gastrulation and early organogenesis. Nature 566, 490–495 (2019).

2. Luo, C. et al. Single-cell methylomes identify neuronal subtypes and regulatory elements in mammalian cortex. Science 357, 600–604 (2017).

3. Argelaguet, R., Mohammed, H., Clark, S. & Stapel, C. Single cell multi-omics profiling reveals a hierarchical epigenetic landscape during mammalian germ layer specification. bioRxiv (2019).

4. Scialdone, A. et al. Computational assignment of cell-cycle stage from single-cell transcriptome data. Methods 85, 54–61 (2015).

