## Appendix for "MOFA+: a probabilistic framework for comprehensive integration of structured single-cell data"

November 4, 2019

#### Overview

Multi-Omics Factor Analysis (MOFA) is a statistical framework aimed at disentangling the sources of variation across data modalities (i.e. a multi-omics dataset).
In this new study we improve the first model formulation (introduced in [2]) with the aim of performing integrative analysis of large-scale datasets across multiple data modalities (views) and across multiple conditions or studies (groups). In this appendix we provide a brief introduction to the original MOFA model, followed by a detailed description of the key model innovations.

#### Mathematical notation

- Matrices are denoted with bold capital letters: $\mathbf{W}$
- Vectors are denoted with bold non-capital letters: $\mathbf{w}$. If the vector comes from a matrix, we will use a single index to indicate the row that it comes from. If two indices are used, the first one corresponds to the row and the second one to the column. The symbol ':' denotes the entire row/column. For instance, $\mathbf{w}_i$ refers to the $i$th row from the $\mathbf{W}$ matrix, whereas $\mathbf{w}_{:,j}$ refers to the $j$th column.
- Scalars are denoted with non-bold and non-capital letters: $w$. If the scalar comes from a 1-dimensional array (a vector), a single subscript will indicate its position in the vector. If the scalar comes from a 2-dimensional array, two indices will be shown at the bottom: the first one corresponding to the row and the second one to the column. For instance, $w_{i,j}$ refers to the value from the $i$th row and the $j$th column of the matrix $\mathbf{W}$, and $w_i$ to the $i$th value of the vector $\mathbf{w}$. In some cases, higher dimensional arrays (tensors) are used, and the use of multiple indices follows the same rationality.
- $\mathbf{0}_k$ is a zero vector of length $K$.
- $\mathbf{I}_k$ is the identity matrix with rank $K$.
- $\mathbb{E}_q[x]$ denotes the expectation of $x$ under the distribution $q$. When the expectations are taken with respect to the same distribution many times, we will avoid cluttered notation and we will instead use $\langle x \rangle$.
- $\mathcal{N}(x \mid \mu, \sigma)$: $x$ follows a univariate normal distribution with mean $\mu$ and variance $\sigma$.
- $\mathcal{G}(x \mid a, b)$: $x$ follows a gamma distribution with shape and rate parameters $a$ and $b$.
- $\text{Beta}(x \mid a, b)$: $x$ follows a beta distribution with shape and rate parameters $a$ and $b$.
- $\text{Ber}(x, \theta)$: $x$ follows a Bernoulli distribution with parameter $\theta$.
- $\mathbb{1}_0$: Dirac delta function centered at 0.
- $Tr(\mathbf{X})$: Trace of the matrix $\mathbf{X}$

### 1 Multi-Omics Factor Analysis v1

This section is reproduced from [2] with some modifications.

Factor analysis models are a probabilistic modelling approach which aim to reduce the dimensionality of a (big) dataset to generate a compressed and denoised representation of the data that is easier to interpret and visualise. Formally, given a dataset $\mathbf{Y} \in \mathbb{R}^{N \times D}$ of $N$ samples and $D$ features, the aim is to explain the dependencies between the observed features by means of a small set of $K$ unobserved (or latent) variables, called factors. The factors capture the global sources of variation in the dataset. Intuitively, this is similar to Principal Component Analysis.
MOFA is a multi-view generalisation of traditional Factor Analysis to $M$ input matrices (or views) $\mathbf{Y}^m \in \mathbb{R}^{N \times D_m}$. Each view consists of non-overlapping features that often represent different assays. However, there is flexibility in the definition of views, depending on the hypothesis of the user. As we demonstrate in the manuscript, one can formulate multi-view problems even from unimodal data. For example, if one has DNA methylation data, one could define as a single view the matrix with all genome-wide CpG measurements, but one could also split this matrix into different views, either by chromosome or by genomic context (i.e. promoters, enhancers, etc.). This formulation could enable the detection of patterns that are shared or specific to different genomic contexts.

Formally, in MOFA, the input data is factorised using the master equation of latent variable models:

$$\mathbf{Y}^m = \mathbf{Z}\mathbf{W}^{mT} + \boldsymbol{\epsilon}^m, \tag{1}$$

where $\mathbf{Z} \in \mathbb{R}^{N \times K}$ is a matrix that contains the factor values and $\mathbf{W}^m \in \mathbb{R}^{D_m \times K}$ are a set of $M$ matrices that define the loadings that relate the high-dimensional space to the low-dimensional latent representation. $\boldsymbol{\epsilon}^m \in \mathbb{R}^{D_m}$ captures the residuals, or the noise.

The structure of the MOFA 1.0 model is depicted in the following figure:

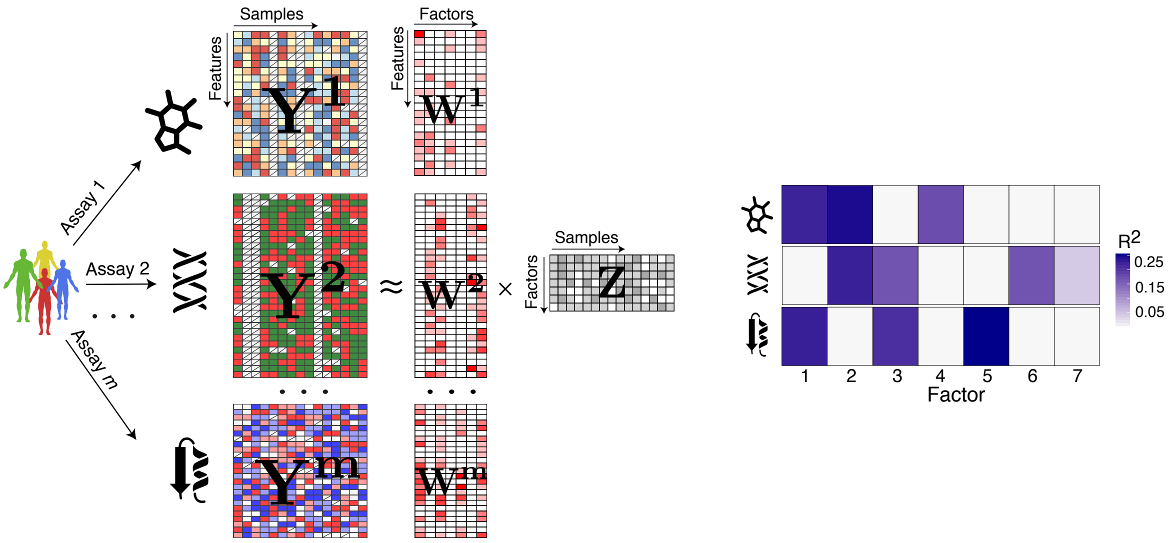

Figure 1: MOFA 1.0 overview. (a) The model takes $M$ data matrices as input ($\mathbf{Y}^1, \cdots, \mathbf{Y}^M$), one or more from each data modality, with co-occurrent samples but features that are not necessarily related and can differ in numbers. MOFA decomposes these matrices into a matrix of factors ($\mathbf{Z}$) and $M$ weight matrices, one for each data modality ($\mathbf{W}^1, \cdots, \mathbf{W}^M$). White cells in the weight matrices correspond to zeros, i.e. inactive features, whereas the cross symbol denotes missing values. (b) The fitted MOFA model can be queried for different downstream analyses, including a variance decomposition to assess the proportion of variance explained by each factor in each data modality.

By default, residuals are assumed to be normally distributed and heteroskedastic:

$$p(\epsilon_d^m) = \mathcal{N}\left(\epsilon_d^m \,|\, 0, 1/\tau_d^m\right). \tag{2}$$

Which results into the following likelihood:

$$p(\mathbf{Y}|\mathbf{W},\mathbf{Z},\mathbf{T}) = \prod_{m=1}^{M} \prod_{d=1}^{D_m} \prod_{n=1}^{N} \mathcal{N}\left(y_{nd}^m \,|\, \mathbf{z}_n^T \mathbf{w}_d^m, 1/\tau_d^m\right), \tag{3}$$

Non-Gaussian noise models are available by replacing the prior distributions from Gaussian to other distributions, such as Bernoulli (for binary data) or Poisson (for count data). The mathematical details can be found in [2]. For simplicity, unless otherwise stated, we will always assume Gaussian noise.

#### 1.1 Interpretation of the factors

Factors capture the global sources of variability in the data. They are analogous to the principal components in Principal Component Analysis (PCA). Mathematically, each factor ordinates cells along a one-dimensional axis centered at zero. Samples with different signs manifest opposite phenotypes along the inferred axis of variation, with higher absolute value indicating a stronger effect.
For example, assume that the $k$-th factor captures the variability associated with cell cycle. We could expect cells in Mitosis to be at one side of the factor axis (irrespective of the sign, only the relative positioning being of importance), whereas cells in G1 phase are expected to be at the other end of the factor axis. Cells with intermediate phenotype, or with no clear phenotype (i.e. no cell cycle genes profiled), are expected to be located around zero (because of the zero-mean prior distribution).

#### 1.2 Interpretation of the loadings

The loadings provide a score for each feature on each factor, and are interpreted in a similar way as the factors. Genes with no association with the factor are expected to have values close to zero, whereas genes with strong association with the factor are expected to have large absolute values. The sign of the loading indicates the direction of the effect: a positive loading indicates that the feature has higher levels in the cells with positive factor values, and vice-versa.
Following the cell cycle example from above, genes that are upregulated in the M phase are expected to have large positive loadings, whereas genes that are downregulated in the M phase (or, equivalently, upregulated in the G1 phase) are expected to have negative loadings.

The following figure shows a real-case example of a Factor capturing the cell cycle effect, with the corresponding loadings:

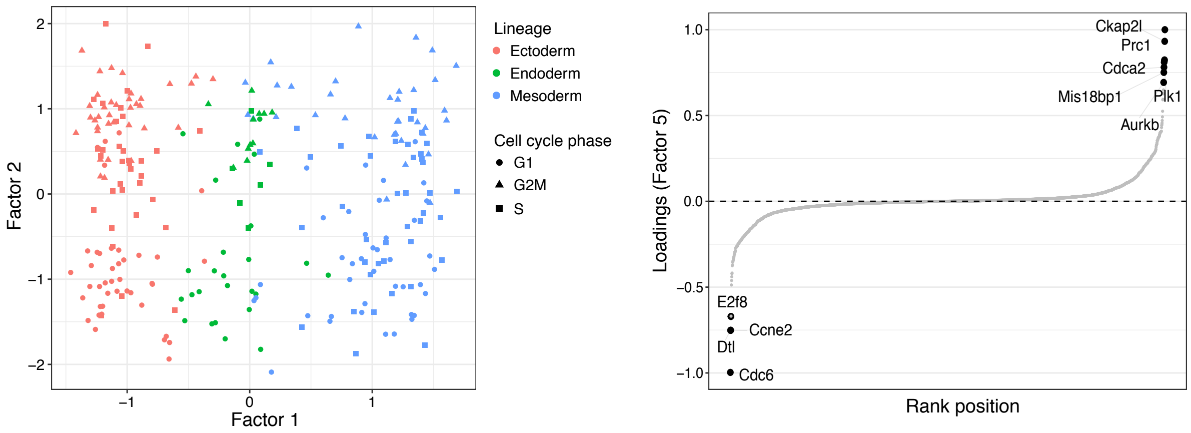

Figure 2: Example of a factor (Factor 2) that captures the cell cycle phenotype. Left shows a scatterplot of Factor 1 and Factor 2, where each dot is a single cell. The cells are colored by the inferred lineage and they are shaped by the inferred cell cycle phase. Right shows the RNA expression loadings for Factor 5. Each dot represents a gene.

Importantly, the scale of the loadings should not be compared across views, only within a view. The scale of the loadings are adjusted to match the distribution of the data modalities, so if you multiple the values of a data modality by two this will yield loadings scaled by a factor of 2. Hence, for simplicity, it is common practice to report them scaled from -1 to 1 (or from 0 to 1 if using the absolute value).

#### 1.3 Interpretation of the noise

The use of a probabilistic framework allows the model to explicitly disentangle the signal (the explained variance) from the noise (the unexplained variance). In the Gaussian framework, large values of $\tau_d^m$ indicate that the model predicts with high accuracy the observations for the feature $d$ in view $m$. In contrast, small values of $\tau_d^m$ are indicative of low predictive power.

#### 1.4 Prior distributions for the factors and the loadings

The key determinant of the model is the regularization used on the prior distributions of the factors and the weights. In the first version of MOFA we defined a standard Gaussian prior on the factors:

$$p(z_{nk}) = \mathcal{N}\left(z_{nk} \,|\, 0, 1\right) \tag{4}$$

Effectively this assumes, *a priori*, independent samples centered at 0. This zero-mean prior induces sparsity, but only to some extent. If the data does not contain meaningful factors, the posterior distributions will also be centered at 0.

In contrast, the weights are assumed to be potentially very sparse, the rationality being that the number of features is very large and that real biological factors are driven by potentially small gene regulatory networks [8]. To achieve this, MOFA encodes two levels of sparsity: (1) a view- and factor-wise sparsity and (2) an individual feature-wise sparsity. The aim of the factor- and view-wise sparsity is to identify which factors are active in which views, such that the weight vector $\mathbf{w}_{:,k}^m$ is shrunk to zero if the factor $k$ does not explain any variation in view $m$. In addition, we place a second layer of sparsity which encourages inactive weights on each individual feature.

Mathematically, we express this as a combination of an Automatic Relevance Determination (ARD) prior [14] for the view- and factor-wise sparsity and a spike-and-slab prior [16] for the feature-wise sparsity:

$$p(w_{kd}^m) = (1 - \theta_k^m)\mathbb{1}_0(w_{kd}^m) + \theta_k^m \mathcal{N}(w_{kd}^m \,|\, 0, 1/\alpha_k^m) \tag{5}$$

However, the spike-and-slab prior contains a Dirac delta function, which makes the inference procedure troublesome. To solve this, we introduce a re-parametrization of the weights $w$ as a product of a Gaussian random variable $\hat{w}$ and a Bernoulli random variable $s$ [24], resulting in the following prior for every single weight $w_{kd}$:

$$p(\hat{w}_{kd}^m, s_{kd}^m) = \mathcal{N}(\hat{w}_{kd}^m \,|\, 0, 1/\alpha_k^m)\,\mathrm{Ber}(s_{kd}^m \,|\, \theta_k^m) \tag{6}$$

In this formulation $\alpha_k^m$ controls the strength of factor $k$ in view $m$ and $\theta_k^m$ controls the fraction of non-zero (active) loadings (i.e. the sparsity levels) of factor $k$ in view $m$. Finally, we allow the model to learn the levels of sparsity and introduce the following hierarchical priors for $\theta$ and $\alpha$:

$$p(\theta_k^m) = \mathrm{Beta}\left(\theta_k^m \,|\, a_0^\theta, b_0^\theta\right) \tag{7}$$
$$p(\alpha_k^m) = \mathcal{G}\left(\alpha_k^m \,|\, a_0^\alpha, b_0^\alpha\right), \tag{8}$$

with hyper-parameters $a_0^\theta, b_0^\theta = 1$ and $a_0^\alpha, b_0^\alpha = 1e^{-3}$ to get uninformative priors. Posterior values of $\theta_k^m$ close to 0 implies that most of the weights of factor $k$ in view $m$ are shrinked to 0 (sparse factor). In contrast, a value of $\theta_k^m$ close to 1 implies that most of the weights are non-zero (non-sparse factor). A small value of $\alpha_k^m$ implies that factor $k$ is active in view $m$. In contrast, a large value of $\alpha_k^m$ implies that factor $k$ is inactive in view $m$.

This achieves the definition of the original MOFA model.

#### 2 Multi-Omics Factor Analysis v2

##### 2.1 Introducing sparsity on the latent space

The use of a view-wise sparsity prior on the loadings relies on the assumption that the features are not independent but structured into different views. Some factors may explain variability in only subsets of views, resulting in a structured sparsity as illustrated in Figure 1.
Following the same logic, the integration of multiple conditions or studies requires breaking the assumption of independent samples and introducing a prior that captures the existence of different groups, such that some factors are allowed to be active in different subsets of groups.
For example, consider the case where we try to integrate two single-cell multi-omics experiments where the first one contains a lineaging event that is not present in the second one. In this scenario, one expects to learn a factor that capture such variability, but this factor should only be active in the first data set. A real case example is shown in the following figure:

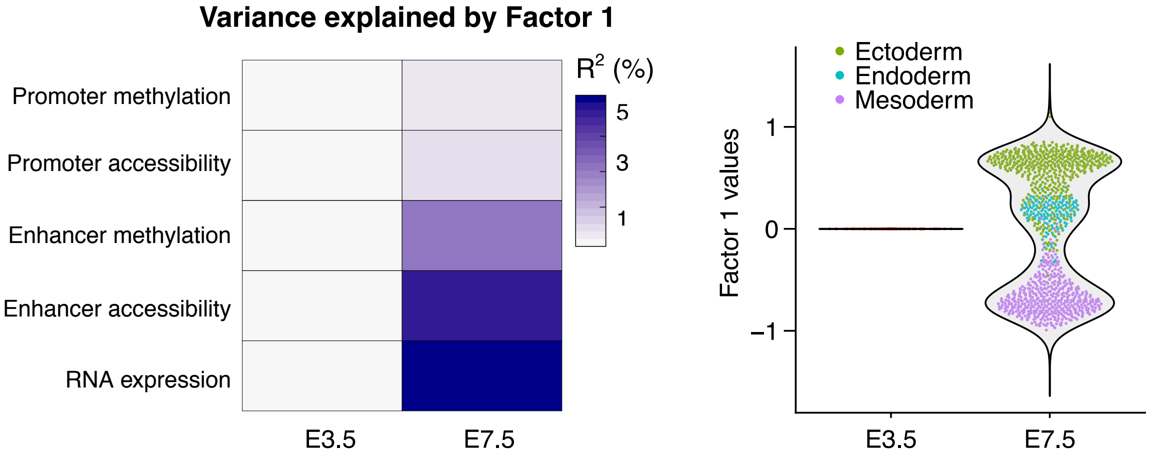

Figure 3: Example of factor that captures a lineaging event during mouse gastrulation. The data consists of six views (RNA expression, DNA methylation of promoters and enhancers, and chromatin accessibility of promoters and enhancers, respectively) and two experiments (embryonic day E3.5 and E7.5). For simplicity, a single factor (Factor 1) is shown. The left plot shows the variance explained by Factor 1 in each view and experiment. The right plot shows Factor 1 values per experiment, where each dot is a single cell.

In addition, if the data has a large number of (noisy) cells, one can expect each factor to be active in only a small fraction of cells.

To formalise the assumptions above, we mirror the double sparsity prior from the loadings:

$$p(\hat{z}_{nk}^{g}, s_{nk}^{g}) = \mathcal{N}(\hat{z}_{nk}^{g} \,|\, 0, 1/\alpha_k^g)\, \text{Ber}(s_{nk}^{g} \,|\, \theta_k^g) \tag{9}$$

$$p(\theta_k^g) = \text{Beta}\left(\theta_k^g \,|\, a_0^\theta, b_0^\theta\right) \tag{10}$$

$$p(\alpha_k^g) = \mathcal{G}\left(\alpha_k^g \,|\, a_0^\alpha, b_0^\alpha\right), \tag{11}$$

where $g$ is the index of the sample groups.

In addition, to account for the fact that different sample groups may exhibit different noise levels, we use the following noise model, where $\epsilon_d^{m,g}$ denotes the residual for a particular view, feature and sample group :

$$p(\epsilon_d^{m,g}) = \mathcal{N}\left(\epsilon_d^{m,g} \,|\, 0, 1/\tau_d^{m,g}\right) \tag{12}$$

$$p(\tau_d^{m,g}) = \mathcal{G}\left(\tau_d^{m,g} \,|\, a_0^\tau, b_0^\tau\right) \tag{13}$$

Effectively, this symmetric sparsity allows the model to integrate multiple views (at the feature level) as well as multiple groups (at the sample level). The graphical model is shown in Figure 4:

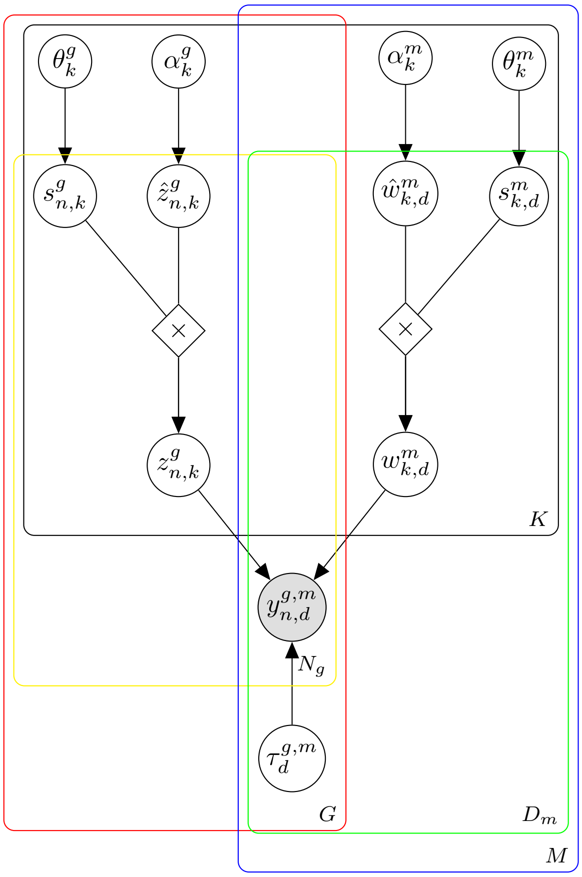

Figure 4: Graphical model for MOFA 2.0. The white circles represent hidden variables that are inferred by the model, whereas the grey circles represent the observed variables. There are a total of five plates, each one representing a dimension of the model: $M$ for the number of views, $G$ for the number of groups, $K$ for the number of factors, $D_m$ for the number of features in view $m$ and $N_g$ for the number of samples in group $g$

##### Solving the rotational invariance problem

Conventional Factor Analysis is invariant to rotation in the latent space[26]. To demonstrate this property, let us apply an arbitrary rotation to the loadings and the factors, specified by the rotation matrix $\mathbf{R} \in \mathbb{R}^{K \times K}$:

$$\tilde{\mathbf{Z}} = \mathbf{Z}\mathbf{R}^{-1}$$
$$\tilde{\mathbf{W}} = \mathbf{R}\mathbf{W}$$

First, note that the model likelihood is unchanged by this rotation, irrespective of the prior distribution used.

$$p(\mathbf{Y}|\tilde{\mathbf{Z}}\tilde{\mathbf{W}}, \tau) = p(\mathbf{Y}|\mathbf{Z}\mathbf{R}^{-1}\mathbf{R}\mathbf{W}, \tau) = p(\mathbf{Y}|\mathbf{Z}\mathbf{W}, \tau)$$

However, the prior distributions of the factors and the loadings are only invariant to rotations when using isotropic Normal priors:

$$\ln p(\mathbf{W}) \propto \sum_{k=1}^{K} \sum_{d=1}^{D} w_{d,k}^2 = \mathrm{Tr}(\mathbf{W}^T\mathbf{W}) = \mathrm{Tr}(\mathbf{W}^T\mathbf{R}^{-1}\mathbf{R}\mathbf{W}) = \mathrm{Tr}(\tilde{\mathbf{W}}^T\tilde{\mathbf{W}})$$

where we have used the property $\mathbf{R}^T = \mathbf{R}^{-1}$ that applies to rotation matrices. The same derivation follows for the factors $\mathbf{Z}$.

In practice, this property renders conventional Factor Analysis unidentifiable, hence limiting its interpretation and applicability. Sparsity assumptions, however, partially address the rotational invariance

problem [10]. When using independent identically distributed spike-and-slab priors, the proof above cannot be applied, hence making the proposed factor analysis model not rotationally invariant.
It is important to remark that the factors are nonetheless invariant to permutations. This implies that, under different initial conditions, the order of the factors is not necessarily the same in independent model fittings. To address this, we manually sort factors *a posteriori* based on total variance explained.

#### 2.2 Scaling up using stochastic variational inference with natural gradients

The size of biological datasets is rapidly increasing, particularly in the field of single cell sequencing, with some studies reporting more than a milion cells[23, 6].
In the original MOFA model, inference was performed using variational Bayes [2, 3, 11]. While this framework is typically faster than sampling-based Monte Carlo approaches, it becomes prohibitively slow with very large datasets, hence motivating the development of a more efficient inference scheme. For this purpose, we derived the stochastic version of the variational inference algorithm developed by [9, 7, 1] for our specific probabilistic model (Figure 4).

In the first two subsections, we briefly review the variational inference algorithm and its stochastic version as proposed by [9]. In the next two subsections, we show how to derive the variational inference algorithm and its stochastic version for the specific MOFA+ probabilistic model.

##### 2.2.1 Review section: Variational Inference (VI)

###### a) Main principles

First, we briefly introduce the variational Bayes framework and show how to turn it into an optimisation problem that can be solved via coordinate or gradient ascent methods. For a detailed mathematical introduction to variational methods we refer the reader to [4, 17, 5, 25, 3].

Let us define a generative model specified by some observations $\mathbf{Y}$ and some latent variables $\mathbf{X}$.
The central aim of Bayesian inference is to obtain the posterior distribution of the latent variables given the observations:

$$p(\mathbf{X}|\mathbf{Y}) = \frac{p(\mathbf{Y},\mathbf{X})}{\int_x p(\mathbf{Y},\mathbf{X})}$$

For most realistic models, the integral in the denominator is intractable and one has to resort to approximations.

In variational inference the true (but intractable) posterior distribution $p(\mathbf{X}|\mathbf{Y})$ is approximated by a simpler (variational) distribution $q(\mathbf{X})$. This distribution is found as the closest approximation to the true posterior, among a predefined family of distributions which are tractable to compute. The distance between the true posterior distribution and an approximate posterior distribution (belonging to the chosen family) is calculated using the KL divergence:

$$\mathrm{KL}(q(\mathbf{X})||p(\mathbf{X}|\mathbf{Y})) = -\int_x q(\mathbf{X}) \log \frac{p(\mathbf{X}|\mathbf{Y})}{q(\mathbf{X})}$$

Note that, if we allowed any possible choice of $q(\mathbf{X})$, then the minimum of this function would occur when $q(\mathbf{X})$ equals the true posterior distribution $p(\mathbf{X}|\mathbf{Y})$. Since the true posterior is intractable to compute, we instead look for the distribution minimizing the KL divergence among a restricted family of approximate distributions.

Doing some calculus it can be shown (see [4, 17]) that the KL divergence $\mathrm{KL}(q(\mathbf{X})||p(\mathbf{X}|\mathbf{Y}))$ is the difference between the log of the marginal probability of the observations $\log(\mathbf{Y})$ and a term $\mathcal{L}(\mathbf{X})$ that is typically called the Evidence Lower Bound (ELBO):

$$\mathrm{KL}(q(\mathbf{X})||p(\mathbf{X}|\mathbf{Y})) = \log(\mathbf{Y}) - \mathcal{L}(\mathbf{X})$$

The first term in the right-hand side is a constant. Hence, as illustrated in Figure 5, minimising the KL divergence is equivalent to maximising $\mathcal{L}(\mathbf{X})$ :

$$\begin{aligned}\mathcal{L}(\mathbf{X}) &= \int_x q(\mathbf{X})\Big(\log\frac{p(\mathbf{X}|\mathbf{Y})}{q(\mathbf{X})} + \log p(\mathbf{Y})\Big)d\mathbf{X} \\ &= \mathbb{E}_q[\log p(\mathbf{X},\mathbf{Y})] - \mathbb{E}_q[\log q(\mathbf{X})]\end{aligned} \tag{14}$$

The first term is the expectation of the log joint probability distribution with respect to the variational distribution. The second term is the entropy of the variational distribution. Importantly, given a simple parametric form of $q(\mathbf{X})$, each of the terms in Equation (14) can be computed in closed form.
In some occasions, we will use the following form for the ELBO:

$$\mathcal{L}(\mathbf{X}) = \mathbb{E}_q[\log p(\mathbf{Y}|\mathbf{X})] + (\mathbb{E}_q[\log p(\mathbf{X})] - \mathbb{E}_q[\log q(\mathbf{X})]) \tag{15}$$

where the first term is the expectation of the log likelihood and the second term is the difference in the expectations of the $p$ and $q$ distributions of each hidden variable.

In conclusion, variational learning involves minimising the KL divergence between $q(\mathbf{X})$ and $p(\mathbf{X}|\mathbf{Y})$ by instead maximising $\mathcal{L}(\mathbf{X})$. The following image summarises the general picture of variational learning:

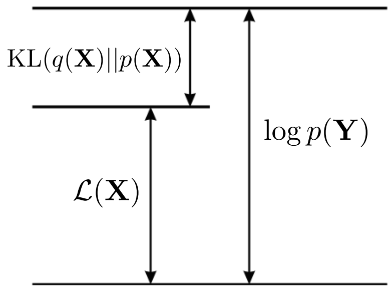

Figure 5: The quantity $\mathcal{L}(\mathbf{X})$ provides a lower bound on the true log marginal likelihood $\log p(\mathbf{Y})$, with the difference being given by the Kullback-Leibler divergence KL($q||p$) between the variational distribution $q(\mathbf{X})$ and the true posterior $p(\mathbf{X}|\mathbf{Y})$

To define the family of distributions where to find the approximate posterior $q(\mathbf{X})$, two approaches are possible : non-parametric or free-form [4], and parametric or fixed-form [25, 5]. We will present both approaches under the mean-field assumption, which assumes that the approximate posterior factorises over $M$ disjoint groups of variables[21]:

$$q(\mathbf{X}) = \prod_{i=1}^{M} q_i(\mathbf{x}_i) \tag{16}$$

**b) Free-form mean-field variational inference**

A common approach is to make no parametric assumptions on the form of the factors $q_i$.
Using calculus of variations (see [4, 17]), it can be shown that the ELBO is maximised with respect to $q_i$ when:

$$\log q_i^*(\mathbf{x}_i) = \mathbb{E}_{-i}[\log p(\mathbf{Y},\mathbf{X})] + \text{const} \tag{17}$$

where $\mathbb{E}_{-i}$ denotes an expectation with respect to the $q$ distributions over all variables $\mathbf{x}_j$ except for $\mathbf{x}_i$. The additive constant is set by normalising the distribution $q_i^*(\mathbf{x}_i)$:

$$q_i^*(\mathbf{x}_i) = \frac{\exp(\mathbb{E}_{-i}[\log p(\mathbf{Y},\mathbf{X})])}{\int \exp(\mathbb{E}_{-i}[\log p(\mathbf{Y},\mathbf{X})])d\mathbf{X}}$$

Equation 17 consists in a set of one equation per model variable $\mathbf{x}_i$, with dependencies in the first and second moment of the other distributions $q_{j\neq i}$ (via the expectation $\mathbb{E}_{-i}$). In practice, this is solved using coordinate ascent, where $q_i$ distributions are updated iteratively while keeping the $q_{j\neq i}$ distributions fixed, until convergence of the ELBO. When conjugate priors are used, the $q_i$ distributions have the same functional forms as the priors $P(\mathbf{x}_i)$, and their parameters can easily be identified from the functional form of $\mathbb{E}_{-i}\left[\ln P(\mathbf{Y},\mathbf{X})\right]$. Again, please refer to [4, 17, 26] for mathematical details.

**c) Fixed-form mean-field variational inference**

An alternative and straightforward choice is to directly define the distribution $q(\mathbf{X})$ to be of the same form as the prior distribution $p(\mathbf{X})$, with (variational) parameters $\mathbf{\Theta}$. Thus, each factor $q_i$ belongs to the same family as the prior over the variables $x_i$, and can be parametrized with its own parameters $\theta_i$ to be tuned.

Importantly, this approach introduces parametric assumptions which only match the free-form mean-field derivation when using conjugate priors. However, for generic models with arbitrary families of distributions, no closed-form variational distributions exist via the free-form mean-field approximation [25, 5].

In the fixed-form approach, the ELBO can be maximised by optimising the parameters $\mathbf{\Theta}$ via conventional numeric optimisation methods. Hence, while the parametric assumption certainly limits the flexibility of variational distributions, the advantage of this formulation is that it opens up the possibility to use (potentially fast) gradient-based methods for the inference procedure [9, 18, 12].

##### 2.2.2 Review section: Stochastic Variational Inference (SVI)

We have seen in the previous section that mean-field variational inference amounts to a coordinate ascent procedure to maximise the evidence lower bound.
A faster alternative to maximise the ELBO with respect to the variational distribution $q(\mathbf{X})$ is to use a stochastic gradient ascent procedure. In the context of variational inference, this approach was proposed by [9] by the name of Stochastic Variational Inference (SVI).
In the followings, we first review briefly the two ingredients required to understand SVI - stochastic gradient ascent and natural gradients - to finally introduce the SVI algorithm.

**a) Stochastic gradient ascent**

Gradient ascent is a common first-order optimization algorithm for finding the maximum of a function [4, 17]. It works iteratively by taking steps proportional to the gradient of the function evaluated at each iteration. Formally, for a differentiable function $F(x)$, the iterative scheme of gradient ascent is:

$$\mathbf{x}^{(t+1)} = \mathbf{x}^{(t)} + \rho^{(t)} \nabla F(\mathbf{x}^{(t)}) \tag{18}$$

At each iteration, the gradient $\nabla F$ is re-evaluated and a step is performed towards its direction. The step size is controlled by $\rho^{(t)}$, a parameter called the learning rate, which is typically adjusted at each iteration $t$ [19].

Gradient ascent is appealing because of its simplicity, but it becomes prohibitively slow with large datasets, mainly because of the computational cost (both in terms of time and memory) associated with the iterative calculation of gradients [22].

Assuming the existence of redundancy in the dataset, a fast approximation of the gradient $\hat{\nabla} F$ can be calculated using a random subset of the data (minibatch). Formally, as in standard gradient ascent, the iterative training schedule proceeds by taking steps of size $\rho$ in the direction of the approximate gradient $\hat{\nabla} F$:

$$\mathbf{x}^{(t+1)} = \mathbf{x}^{(t)} + \rho^{(t)} \hat{\nabla} F(\mathbf{x}^{(t)}) \tag{19}$$

The step size $\rho^{(t)}$ can also be adjusted at each iteration $t$. When the series $\rho(t)$ satisfies the Robbins-Monro conditions: $\sum_t \rho^{(t)} = \infty$ and $\sum_t (\rho^{(t)})^2 < \infty$, $F$ is guaranteed to converge to a local maximum [20]. If $F$ is not convex, the algorithm is sensible to the initialisation $\mathbf{x}^{t=0}$.

**b) Natural gradient ascent**

Let us consider a model with a single hidden variable $x$ and corresponding variational parameter $\theta$. The objective function is the ELBO $\mathcal{L}(\theta)$. From the definition of the gradient:

$$\nabla \mathcal{L}(\theta) = \lim_{||h|| \to 0} \frac{\mathcal{L}(\theta + h) - \mathcal{L}(\theta)}{||h||}$$

where $h$ represents an infinitesimally small positive step in the space of $\theta$.
To find the steepest gradient, one would need to search over all possible directions $d$ in an infinitely small distance $h$, and select the $\hat{d}$ with the largest gradient:

$$\nabla \mathcal{L}(\theta) = \lim_{h \to 0} \frac{1}{h} \underset{d\, s.t. \|d\|=h}{\arg\max}\ \mathcal{L}(\theta + d) - \mathcal{L}(\theta)$$

Notice that the neighborhood of $\theta$ is measured in terms of its Euclidean norm, and the direction of steepest ascent is hence dependent on the Euclidean geometry of the $\theta$ space. This is the key problematic issue when working with probability distributions. A small step from $\theta^{(t)}$ to $\theta^{(t+1)}$ does not guarantee an equivalently small change from $\mathcal{L}(\theta^{(t)})$ to $\mathcal{L}(\theta^{(t+1)})$. As an example, consider four random variables:

$$\begin{aligned} \psi_1 &\sim \mathcal{N}(0 \,|\, 5) \qquad & \psi_3 &\sim \mathcal{N}(0 \,|\, 1) \\ \psi_2 &\sim \mathcal{N}(10 \,|\, 5) \qquad & \psi_4 &\sim \mathcal{N}(10 \,|\, 1) \end{aligned} \tag{20}$$

Using the Euclidean metric, the distance between $\psi_1$ and $\psi_2$ is the same as the distance between $\psi_3$ and $\psi_4$. However, the distance in distribution space (measured for example by the KL divergence) is clearly much larger between $\psi_1$ and $\psi_2$ than between $\psi_3$ and $\psi_4$ (Figure 6).

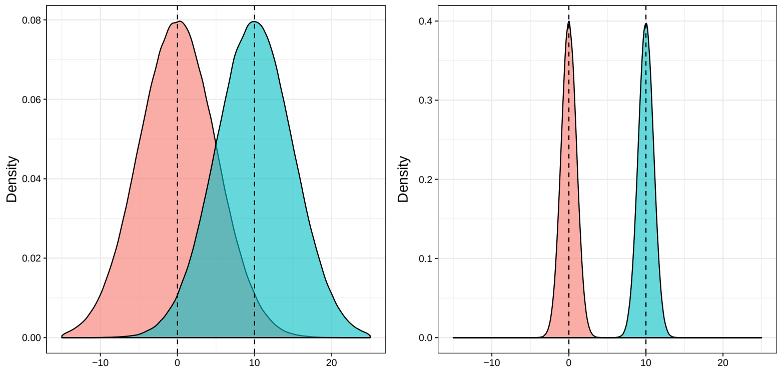

Figure 6: Illustration of the problem of using Euclidean distances to measure distances between parameters of distributions. In both plots, the red and blue distributions are separated by the same Euclidean distance of 100. Yet, the distance in probability space between the two distributions is intuitively much higher in the right plot.

Hence, rather than using the Euclidean distance, it is more appropriate to use the KL divergence as a distance metric:

$$\nabla_{KL} \mathcal{L}(\theta) = \lim_{h \to 0} \frac{1}{h} \underset{d\, s.t. KL[p_\theta || p_{\theta+d}]=h}{\arg\max}\ \mathcal{L}(\theta + d) - \mathcal{L}(\theta)$$

The direction of steepest ascent measured by the KL divergence is called the natural gradient [1, 15]. By introducing Lagrange multipliers and Taylor expansions, one can solve the optimisation problem to obtain the direction of the steepest natural gradient (see [1, 13]). The solution corresponds to the standard (Euclidean) gradient pre-multiplied by the inverse of the Fisher Information Matrix of $q(x|\theta)$:

$$\hat{d}_{KL} \propto \mathbf{F}^{-1}(\theta) \nabla_\theta \mathcal{L}(\theta) \tag{21}$$

where $\mathbf{F}(\theta)$ is defined as

$$\mathbf{F}(\theta) = \mathbb{E}_{q(x|\theta)}[(\nabla_\theta \log q(x|\theta))(\nabla_\theta \log q(x|\theta))^T]$$

Effectively, the premultiplication by $\mathbf{F}^{-1}$ takes into account the local curvate of $q(\theta)$ in distribution space. Importantly, when $q(x|\theta)$ belongs to the exponential family, the Fisher Information matrix is simply the Hessian of the log normalizer.

In conclusion, while the standard gradient points to the direction of steepest ascent in Euclidean space, the natural gradient points to the direction of steepest ascent in a space where distances are defined by the KL divergence [13, 1, 9].

##### c) SVI is a stochastic natural gradient ascent

We can finally introduce the stochastic variational inference (SVI) algorithm, which may indeed be seen as a stochastic natural gradient ascent. This approach was proposed by [9] to solve the fixed-form mean-field variational inference introduced in the previous section.
The stochastic nature of the approach is interesting when one dimension of the matrix of observed variables is much larger than the others. In our case, it corresponds to $N$, the number of samples (or cells).

In this section we will summarise the key principles to understand the SVI algorithm. For a complete mathematical derivation we refer the readers to [9]
As a starting point, we classify the variables of the probabilistic model into four different types:

- observations ($\mathbf{Y}$): $N$ different vectors $\mathbf{y}_n$ which contain the observed variables for the $n$-th sample.
- local (hidden) variables ($\mathbf{Z}$): $N$ different vectors $\mathbf{z}_n$ which contain all $K$ hidden variables associated with each sample $n$.
- global (hidden) variables ($\boldsymbol{\beta}$): one vector that contains all $B$ hidden variables not indexed by $n$.
- parameters ($\boldsymbol{\alpha}$): a vector that contains all fixed parameters for the global variables.

The distinction between local and global variables lies on the conditional dependencies. Given the global variables $\beta$, the $n$th local variable $\mathbf{z}_n$ is conditionally independent from any other observation $\mathbf{y}_j$ or local variable $\mathbf{z}_j$ (where $j \neq n$):

$$p(\mathbf{y}_n, \mathbf{z}_n | \mathbf{y}_j, \mathbf{z}_{nj}, \beta, \alpha) = p(\mathbf{y}_n, \mathbf{z}_n | \beta, \alpha)$$

Under the assumptions that the complete conditionals in the model are in the exponential family [1] — which notably imply that $p(\boldsymbol{\beta})$ and $p(\mathbf{y}_n, \mathbf{z}_n|\boldsymbol{\beta})$ are also in the exponential family —, [9] showed that:

- The update equations of the classic variational inference algorithm (which is a coordinate ascent using the classic gradient of the ELBO) are also satisfied when following the natural gradient of the ELBO
- The ELBO decomposes into a global term and a sum of local terms — each local term specific to a particular sample —, and so does its natural gradient, which makes the stochastic gradient algorithm applicable to the natural gradient.
- The natural gradient of the ELBO is cheaper to compute than the classic gradient.

To illustrate the key points above, let us introduce a general formula for the ELBO (the objective function) in terms of the four types of variables defined above. For this purpose, we first introduce the following notations:

- the prior distributions are members of the exponential family :

$$\begin{aligned} p(\beta|\alpha_\beta) &= h(\beta)\exp\{\eta_g(\alpha_\beta)t(\beta) - a_g(\alpha_\beta)\} \\ p(z_{nk}|\alpha_z) &= h(z_{nk})\exp\{\eta_l(\alpha_z)t(z_{nk}) - a_l(\alpha_z)\} \end{aligned} \tag{22}$$

- the complete conditionals are members of the exponential family:

$$\begin{aligned} p(\beta|\mathbf{Y}, \mathbf{Z}, \boldsymbol{\alpha}) &= h(\beta)\exp\{\eta_g(\mathbf{Y}, \mathbf{Z}, \boldsymbol{\alpha})^T t(\beta) - a_g(\eta_g(\mathbf{Y}, \mathbf{Z}, \boldsymbol{\alpha}))\} \\ p(\mathbf{z}_n|\mathbf{y}_{nj}, \mathbf{z}_{nj}, \beta) &= h(\mathbf{z}_n)\exp\{\eta_l(\mathbf{y}_{nj}, \mathbf{z}_{nj}, \beta)^T t(\mathbf{z}_n) - a_l(\eta_l(\mathbf{y}_{nj}, \mathbf{z}_{nj}, \beta))\} \end{aligned} \tag{23}$$

- the variational distributions in the fixed-form mean-field assumption belong to the same exponential family as their priors:

$$q(\mathbf{z}, \beta) = q(\beta|\lambda)\prod_{n=1}^{N}\prod_{k=1}^{K} p(z_{nk}|\phi_{nk}) \tag{24}$$

$$q(\beta|\lambda) = h(\beta)\exp\{\eta_g(\lambda)t(\beta) - a_g(\lambda)\} \tag{25}$$

$$q(z_{nk}|\phi_{nk}) = h(z_{nk})\exp\{\eta_l(\phi_n)t(z_{nk}) - a_l(z_{nk})\} \tag{26}$$

where $\lambda$ are the parameters governing the global variable and $\phi_{nk}$ are the parameters governing the $k$-th local variable for the $n$-th sample.

[1] a complete conditional is the distribution of a hidden variable given all other variables, hidden or observed

Following this, the ELBO (Equation (14)), factorises as:

$$\begin{aligned}\mathcal{L} &= \mathbb{E}_{q(\mathbf{Z},\beta)}[\log p(\mathbf{Y}, \mathbf{Z}, \beta)] - \mathbb{E}_{q(\mathbf{Z})}[\log q(\mathbf{Z})] - \mathbb{E}_{q(\beta)}[\log q(\beta)] \\ &= \sum_{n=1}^{N} \mathbb{E}_{q(\mathbf{z}_n,\beta)}[\log p(\mathbf{y}_n, \mathbf{z}_n, \beta)] - \sum_{n=1}^{N}\sum_{k=1}^{K} \mathbb{E}_{q(z_{nk})}[\log q(z_{nk})] - \mathbb{E}_{q(\beta)}[\log q(\beta)]\end{aligned} \tag{27}$$

Notice the existence of global terms (which do not depend on $n$) and local terms (which do depend on $n$). We can further derive the corresponding noisy estimate by using a randomly sampled mini-batch of size $S$:

$$\mathcal{L}^S = \frac{N}{S}\sum_{n=1}^{S} \mathbb{E}_{q(\mathbf{z}_n,\beta)}[\log p(\mathbf{y}_n, \mathbf{z}_n, \beta)] - \frac{N}{S}\sum_{s=1}^{S}\sum_{k=1}^{K} \mathbb{E}_{q(z_{nk})}[\log q(z_{nk})] - \mathbb{E}_{q(\beta)}[\log q(\beta)]$$

where the factor $N/S$ ensures that the estimate is unbiased.

To derive the variational updates, we need to compute the natural gradient of $\mathcal{L}^S$ with respect to the global and the local parameters at a turn.

Taking the natural gradient of $\mathcal{L}^S$ with respect to the local parameter $\phi_{nk}$ leads to :

$$\hat{\nabla}_\phi \mathcal{L}(\phi_{nk}) = \mathbb{E}_{q(\beta,\mathbf{z}_{nj})}[\eta_l(\mathbf{y}_n, \mathbf{z}_{nj}, \beta)] - \phi_{nk}$$

Similarly, taking the natural gradient of $\mathcal{L}^S$ with respect to the global parameter $\lambda$ leads to :

$$\hat{\nabla}_\lambda \mathcal{L}^S(\lambda) = \mathbb{E}_{q(z)}\left[\frac{N}{S}\eta_g(\mathbf{Y}_{1:S}, \mathbf{Z}_{1:S}, \boldsymbol{\alpha})\right] - \lambda \tag{28}$$

If we had taken the natural gradient of $\mathcal{L}$ instead of $\mathcal{L}^S$, we would have obtained:

$$\hat{\nabla}_\lambda \mathcal{L}(\lambda) = \mathbb{E}_{q(z)}\left[\eta_g(\mathbf{Y}, \mathbf{Z}, \boldsymbol{\alpha})\right] - \lambda \tag{29}$$

which we can compare to the equation obtained using the standard (Euclidean) gradient:

$$\nabla_\lambda \mathcal{L}(\lambda) = \nabla^2_\lambda a_g(\lambda)(\mathbb{E}_{q(z)}[\eta_g(\mathbf{Y}, \mathbf{Z}, \boldsymbol{\alpha})] - \lambda) \tag{30}$$

The standard gradient requires the Hessian of the log normalizer (i.e. the Fisher Information Matrix) to be explicitly computed at each iteration. Remarkably, in the natural gradient this term has canceled out (see Equation (21)), which leads to a much cheaper computation.

#### d) SVI algorithm vs. VI algorithm

We can notice that the SVI algorithm (with natural gradients) is equivalent to the non-stochastic VI algorithm when using a step-size 1 and when each mini-batch is equal to the full dataset:

**Algorithm 1** Mean-field variational inference

Initialise the global parameters $\boldsymbol{\lambda}^{(t=0)}$ randomly
**repeat**
  **for** each local variational parameter $\phi_{nk}$ **do**

$$\phi_{nk}^{(t+1)} = \mathbb{E}_{q^{(t)}(\boldsymbol{\beta},\mathbf{z}_{nj})}[\eta_l(\mathbf{y}_n, \mathbf{z}_{nj}, \boldsymbol{\beta})]$$

  **end for**
  **for** each global variational parameter $\lambda$ **do**

$$\lambda^{(t+1)} = \mathbb{E}_{q^{(t+1)}(z)}[\eta_g(\mathbf{Y}, \mathbf{Z}, \boldsymbol{\alpha})]$$

  **end for**
**until** ELBO convergence

**Algorithm 2** Stochastic mean-field variational inference

Initialise the global parameters $\boldsymbol{\lambda}^{(t=0)}$ randomly.
Initialise step size $\rho^{(t=0)}$
**repeat**
  sample $\mathcal{B}$ a mini-batch of samples of size $S$
  **for** each local variational parameter $\phi_{nk}$ such that $n$ is in batch $\mathcal{B}$ **do**

$$\phi_{nk}^{(t+1)} = \mathbb{E}_{q^{(t)}(\boldsymbol{\beta}, \mathbf{z}_{nj})}[\eta_l(\mathbf{y}_n, \mathbf{z}_{nj}, \boldsymbol{\beta})]$$

  **end for**
  **for** each global variational parameter $\lambda$ **do**

$$\begin{aligned} \lambda^{(t+1)} &= \lambda^{(t)} + \rho^{(t)} \hat{\nabla}_\lambda \mathcal{L}^S(\lambda) \\ &= (1 - \rho^{(t)})\lambda^{(t)} + \rho^{(t)} \mathbb{E}_{q^{(t+1)}(z)} \left[ \frac{N}{S} \eta_g(\mathbf{Y}_{[n\in\mathcal{B}],:}, \mathbf{Z}_{[n\in\mathcal{B}],:}, \boldsymbol{\alpha}) \right] \end{aligned} \tag{31}$$

    where $[n \in \mathcal{B}]$ denotes the subset of indices corresponding to the samples in $\mathcal{B}$
  **end for**
**until** ELBO convergence

##### e) Hyperparameters

Stochastic variational inference algorithm has two critical hyperparameters:

- The **batch size** $S$ controls the number of samples that are used to compute the gradients at each iteration. A trade-off exists where high batch sizes lead to a slower computation of the gradient but to a less noisy estimate of the gradient.
- The **learning rate** $\rho^{(t)}$ controls the step size in the direction of the natural gradient, with high learning rates leading to higher step sizes. Also, in the natural gradient setting, the learning rate controls how much memory from previous iterations is translated to the current updates. The particular case of a constant $\rho = 1$ yields no memory from previous iterations, which is the particular case of the standard gradient ascent. To ensure proper convergence, the learning rate is usually decayed during training by a pre-defined function. How to adapt the learning rate is an extensive area of research [19].

###### 2.2.3 Deriving the variational inference algorithm

We applied the variational inference principle previously introduced. We looked for an approximate distribution of the true posterior which is the closest to this true posterior (according to the KL divergence distance) among the following family of factorized distributions:

$$\begin{aligned}
q(\mathbf{X}) &= q\left(\{\widehat{\mathbf{Z}^{\mathbf{g}}}, \mathbf{S}^{\mathbf{g}}, \alpha^{\mathbf{g}}, \theta^{\mathbf{g}}\}_{\mathbf{g}\in[\![\mathbf{1};\mathbf{G}]\!]}, \{\widehat{\mathbf{W}^{\mathbf{m}}}, \mathbf{S}^{\mathbf{m}}, \alpha^{\mathbf{m}}, \theta^{\mathbf{m}}\}_{\mathbf{m}\in[\![\mathbf{1};\mathbf{M}]\!]}, \{\tau^{\mathbf{gm}}\}_{\mathbf{g}\in[\![\mathbf{1};\mathbf{G}]\!],\ \mathbf{m}\in[\![\mathbf{1};\mathbf{M}]\!]}\right) \\
&= \prod_{g=1}^{G}\prod_{n=1}^{N_g}\prod_{k=1}^{K} q(\hat{z}^g_{nk}, s^g_{nk}) \prod_{g=1}^{G}\prod_{k=1}^{K} q(\alpha^g_k) \prod_{g=1}^{G}\prod_{k=1}^{K} q(\theta^g_k) \\
&\times \prod_{m=1}^{M}\prod_{d=1}^{D_m}\prod_{k=1}^{K} q(\hat{w}^m_{kd}, s^m_{kd}) \prod_{m=1}^{M}\prod_{k=1}^{K} q(\alpha^m_k) \prod_{m=1}^{M}\prod_{k=1}^{K} q(\theta^m_k) \\
&\times \prod_{g=1}^{G}\prod_{m=1}^{M}\prod_{d=1}^{D_m} q(\tau^{gm}_d)
\end{aligned}$$

Inspired by [24], we did not look for fully factorized distributions like in the mean-field distribution, since we can hardly assume $\hat{w}^m_k$ and $s^m_k$ to be independent in the posterior ($\theta_k$ and $\alpha_k$ correspond respectively to the amount and the variability of the non-zero loadings for factor $k$).

Since the posterior is fully factorized in terms of the following variables $\{(\hat{z}^g_{nk}, s^g_{nk}), \alpha^g_k, \theta^g_k, (\hat{w}^m_{kd}, s^m_{kd}), \alpha^m_k, \theta^m_k, \tau^{gm}_d\}$, we could derive from equation 17 update equations for the parameters of the following factors : $q(\hat{z}^g_{nk})$, $q(\hat{z}^g_{nk}|s^g_{nk}=0)$, $q(\hat{z}^g_{nk}|s^g_{nk}=1)$, $q(\alpha^g_k)$, $q(\theta^g_k)$, $q(\hat{w}^m_{kd}|s^m_{kd}=0)$, $q(\hat{w}^m_{kd}|s^m_{kd}=1)$, $q(\alpha^m_k)$, $q(\theta^m_k)$, and $q(\tau^{gm}_d)$. In order to derive update equations for the parameters of $q(s^g_{nk})$ (and similarly for $q(s^m_{kd})$), we adapted the reasoning done in [4] to show that, when the functional $\mathcal{L}(q)$ is maximized, then we get the following equation (where we denote $X' = X \setminus \{\hat{z}^g_{nk}, s^g_{nk}\}$) and which slightly differs from equation 17 :

$$\begin{aligned}
\log q(s^g_{nk}) =& \mathbb{E}_{X',\hat{z}^g_{nk}|s^g_{nk}}\left[\log \frac{p(Y,X)}{q(\hat{z}^g_{nk}|s^g_{nk})}\right] \\
=& \mathbb{E}_{X',\hat{z}^g_{nk}|s^g_{nk}} \log p(Y|X) + \mathbb{E}_{X',\hat{z}^g_{nk}|s^g_{nk}} \log p(\hat{z}^g_{nk}|\alpha^g_k) + \mathbb{E}_{X',\hat{z}^g_{nk}|s^g_{nk}} \log p(s^g_{nk}) \\
&- \mathbb{E}_{X',\hat{z}^g_{nk}|s^g_{nk}} \log q(\hat{z}^g_{nk}|s^g_{nk})
\end{aligned} \tag{32}$$

Below we give the explicit update equations for every hidden variable of MOFA 2.0 model which are applied at each iteration of the classic variational inference algorithm.

**a) Update equations**

**Non-sparse factors**

For every group $g$, sample $n$ and factor $k$:

Prior distribution $p(z^g_{nk})$:

$$p(z^g_{nk}) = \mathcal{N}(z^g_{nk}|0,1)$$

Variational distribution $q(z^g_{nk})$:

$$q(z^g_{nk}) = \mathcal{N}(z^g_{nk}|\mu_{z^g_{nk}}, \sigma^2_{z^g_{nk}}) \tag{33}$$

where

$$\begin{aligned}
\sigma^2_{z^g_{nk}} &= \left(\sum_{m=1}^{M}\sum_{d=1}^{D_m}\langle\tau^{gm}_d\rangle\langle(w^m_{kd})^2\rangle + 1\right)^{-1} \\
\mu_{z^g_{nk}} &= \sigma^2_{z^g_{nk}} \sum_{m=1}^{M}\sum_{d=1}^{D_m}\langle\tau^{gm}_d\rangle\langle w^m_{kd}\rangle\left(\sum_{g=1}^{G} y^{gm}_{nd} - \sum_{j\neq k}\langle w^m_{jd}\rangle\langle z^g_{nj}\rangle\right)
\end{aligned} \tag{34}$$

**Sparse factors (with spike-and-slab prior)**

For every group $g$, sample $n$ and factor $k$:

Prior distribution $p(\hat{z}_{nk}^g, s_{nk}^g)$:

$$p(\hat{z}_{nk}^g, s_{nk}^g) = \mathcal{N}(\hat{z}_{nk}^g \,|\, 0,\ 1/\alpha_k^g)\,\text{Ber}(s_{nk}^g \,|\, \theta_k^g) \tag{35}$$

Variational distribution $q(\hat{z}_{nk}^g, s_{nk}^g)$:

Update for $q(s_{nk}^g)$:

$$q(s_{nk}^g) = \text{Ber}(s_{nk}^g|\gamma_{nk}^g) \tag{36}$$

with

$$\begin{aligned}
\gamma_{nk}^g &= \frac{1}{1+\exp(-\lambda_{nk}^g)} \\
\lambda_{nk}^g &= \langle \ln \frac{\theta}{1-\theta} \rangle + 0.5 \ln \frac{\langle \alpha_k^g \rangle}{\langle \tau_d^{gm} \rangle} - 0.5 \ln \left( \sum_{m=1}^{M} \sum_{d=1}^{D_m} \langle (w_{kd}^m)^2 \rangle + \frac{\langle \alpha_k^g \rangle}{\langle \tau_d^{gm} \rangle} \right) \\
&+ \frac{\langle \tau_d^{gm} \rangle}{2} \frac{\left( \sum_{m=1}^{M} \sum_{d=1}^{D_m} y_{nd}^{gm} \langle w_{kd}^m \rangle - \sum_{j \neq k} \langle s_{nj}^g \hat{z}_{nj}^g \rangle \sum_{m=1}^{M} \sum_{d=1}^{D_m} \langle w_{kd}^m \rangle \langle w_{jd}^m \rangle \right)^2}{\sum_{m=1}^{M} \sum_{d=1}^{D_m} \langle (w_{kd}^m)^2 \rangle + \frac{\langle \alpha_k^g \rangle}{\langle \tau_d^{gm} \rangle}}
\end{aligned} \tag{37}$$

Update for $q(\hat{z}_{nk}^g)$:

$$\begin{aligned}
q(\hat{z}_{nk}^g | s_{nk}^g = 0) &= \mathcal{N}\left(\hat{z}_{nk}^g | 0, 1/\alpha_k^g\right) \\
q(\hat{z}_{nk}^g | s_{nk}^g = 1) &= \mathcal{N}\left(\hat{z}_{nk}^g \middle| \mu_{z_{nk}^g}, \sigma_{z_{nk}^g}^2\right)
\end{aligned} \tag{38}$$

with

$$\begin{aligned}
\mu_{z_{nk}^g} &= \frac{\sum_{m=1}^{M} \sum_{d=1}^{D_m} y_{nd}^{m,g} \langle w_{kd}^m \rangle - \sum_{j \neq k} \langle s_{nj}^g \hat{z}_{nj}^g \rangle \sum_{m=1}^{M} \sum_{d=1}^{D_m} \langle w_{kd}^m \rangle \langle w_{jd}^m \rangle}{\sum_{m=1}^{M} \sum_{d=1}^{D_m} \langle (w_{kd}^m)^2 \rangle + \frac{\langle \alpha_k^g \rangle}{\langle \tau_d^{gm} \rangle}} \\
\sigma_{z_{nk}^g}^2 &= \frac{\langle \tau_d^{gm} \rangle^{-1}}{\sum_{m=1}^{M} \sum_{d=1}^{D_m} \langle (w_{kd}^m)^2 \rangle + \frac{\langle \alpha_k^g \rangle}{\langle \tau_d^{gm} \rangle}}
\end{aligned} \tag{39}$$

**ARD precision of the factors**

For every group $g$ and factor $k$:

Prior distribution:

$$p(\alpha_k^g) = \mathcal{G}\left(\alpha_k^g \,|\, a_0^\alpha, b_0^\alpha\right) \tag{40}$$

Variational distribution $q(\alpha_k^g)$:

$$q(\alpha_k^g) = \mathcal{G}\left(\alpha_k^g \,|\, \hat{a}_{gk}^\alpha, \hat{b}_{gk}^\alpha\right) \tag{41}$$

where:

$$\begin{aligned}
\hat{a}_{gk}^\alpha &= a_0^\alpha + \frac{N_g}{2} \\
\hat{b}_{gk}^\alpha &= b_0^\alpha + \frac{\sum_{n=1}^{N_g} \langle (\hat{z}_{nk}^g)^2 \rangle}{2}
\end{aligned} \tag{42}$$

**Sparsity parameter of the factors**

For every group $g$ and factor $k$:

Prior distribution:

$$p(\theta_k^g) = \text{Beta}\left(\theta_k^g \,|\, a_0^\theta, b_0^\theta\right) \tag{43}$$

Variational distribution:

$$q(\theta_k^g) = \text{Beta}\left(\theta_k^g \,|\, \hat{a}_{gk}^\theta, \hat{b}_{gk}^\theta\right) \tag{44}$$

where

$$\begin{aligned} \hat{a}_{gk}^\theta &= \sum_{n=1}^{N_g} \langle s_{nk}^g \rangle + a_0^\theta \\ \hat{b}_{gk}^\theta &= b_0^\theta - \sum_{n=1}^{N_g} \langle s_{nk}^g \rangle + N_g \end{aligned} \tag{45}$$

**Non-sparse weights**

For every view $m$, feature $d$ and factor $k$:

Prior distribution $p(w_{kd}^m)$:

$$p(w_{kd}^m) = \mathcal{N}(w_{kd}^m | 0, 1)$$

Variational distribution $q(w_{kd}^m)$:

$$q(w_{kd}^m) = \mathcal{N}(w_{kd}^m | \mu_{w_{kd}^m}, \sigma^2_{w_{kd}^m}) \tag{46}$$

where

$$\begin{aligned} \sigma^2_{w_{kd}^m} &= \left( \sum_{g=1}^{G} \sum_{n=1}^{N_g} \langle \tau_d^{gm} \rangle \langle (z_{nk}^g)^2 \rangle + 1 \right)^{-1} \\ \mu_{w_{kd}^m} &= \sigma^2_{w_{kd}^m} \sum_{g=1}^{G} \sum_{n=1}^{N_g} \langle \tau_d^{gm} \rangle \langle z_{nk}^g \rangle \left( \sum_{m=1}^{M} y_{nd}^{gm} - \sum_{j \neq k} \langle z_{nj}^g \rangle \langle w_{jd}^m \rangle \right) \end{aligned} \tag{47}$$

**Sparse weights (with spike-and-slab prior)**

For every view $m$, feature $d$ and factor $k$:

Prior distribution $p(\hat{w}_{kd}^m, s_{kd}^m)$:

$$p(\hat{w}_{kd}^m, s_{kd}^m) = \mathcal{N}(\hat{w}_{kd}^m \,|\, 0,\ 1/\alpha_k^m)\, \text{Ber}(s_{kd}^m \,|\, \theta_k^m) \tag{48}$$

Variational distribution $q(\hat{w}_{kd}^m, s_{kd}^m)$:

Update for $q(s_{kd}^m)$:

$$q(s_{kd}^m) = \text{Ber}(s_{kd}^m | \gamma_{kd}^m) \tag{49}$$

with

$$\gamma_{kd}^m = \frac{1}{1+\exp(-\lambda_{kd}^m)}$$

$$\lambda_{kd}^m = \langle \ln \frac{\theta}{1-\theta} \rangle + 0.5 \ln \frac{\langle \alpha_k^m \rangle}{\langle \tau_d^{gm} \rangle} - 0.5 \ln \left( \sum_{g=1}^{G} \sum_{n=1}^{N_g} \langle (z_{nk}^g)^2 \rangle + \frac{\langle \alpha_k^m \rangle}{\langle \tau_d^{gm} \rangle} \right) \tag{50}$$

$$+ \frac{\langle \tau_d^{gm} \rangle}{2} \frac{\left( \sum_{g=1}^{G} \sum_{n=1}^{N_g} y_{nd}^{gm} \langle z_{nk}^g \rangle - \sum_{j \neq k} \langle s_{jd}^m \hat{w}_{jd}^m \rangle \sum_{g=1}^{G} \sum_{n=1}^{N_g} \langle z_{nk}^g \rangle \langle z_{nj}^g \rangle \right)^2}{\sum_{g=1}^{G} \sum_{n=1}^{N_g} \langle (z_{nk}^g)^2 \rangle + \frac{\langle \alpha_k^m \rangle}{\langle \tau_d^{gm} \rangle}}$$

Update for $q(\hat{w}_{kd}^m)$:

$$q(\hat{w}_{kd}^m | s_{kd}^m = 0) = \mathcal{N}\left(\hat{w}_{kd}^m | 0, 1/\alpha_k^m\right)$$
$$q(\hat{w}_{kd}^m | s_{kd}^m = 1) = \mathcal{N}\left(\hat{w}_{kd}^m \middle| \mu_{w_{kd}^m}, \sigma_{w_{kd}^m}^2\right) \tag{51}$$

with

$$\mu_{w_{kd}^m} = \frac{\sum_{g=1}^{G} \sum_{n=1}^{N_g} y_{nd}^{gm} \langle z_{nk}^g \rangle - \sum_{j \neq k} \langle s_{jd}^m \hat{w}_{jd}^m \rangle \sum_{g=1}^{G} \sum_{n=1}^{N_g} \langle z_{nk}^g \rangle \langle z_{nj}^g \rangle}{\sum_{g=1}^{G} \sum_{n=1}^{N_g} \langle (z_{nk}^g)^2 \rangle + \frac{\langle \alpha_k^m \rangle}{\langle \tau_d^{gm} \rangle}}$$
$$\sigma_{w_{kd}^m}^2 = \frac{\langle \tau_d^{gm} \rangle^{-1}}{\sum_{g=1}^{G} \sum_{n=1}^{N_g} \langle (z_{nk}^g)^2 \rangle + \frac{\langle \alpha_k^m \rangle}{\langle \tau_d^{gm} \rangle}} \tag{52}$$

**ARD precision of the weights**

For every view $m$ and factor $k$:

Prior distribution $p(\alpha_k^m)$:

$$p(\alpha_k^m) = \mathcal{G}\left(\alpha_k^m \mid a_0^\alpha, b_0^\alpha\right)$$

Variational distribution $q(\alpha_k^m)$:

$$q(\alpha_k^m) = \mathcal{G}\left(\alpha_k^m \mid \hat{a}_{mk}^\alpha, \hat{b}_{mk}^\alpha\right) \tag{53}$$

where:

$$\hat{a}_{mk}^\alpha = a_0^\alpha + \frac{D_m}{2}$$
$$\hat{b}_{mk}^\alpha = b_0^\alpha + \frac{\sum_{d=1}^{D_m} \langle (\hat{w}_{kd}^m)^2 \rangle}{2} \tag{54}$$

**Sparsity parameter of the weights**

For every view $m$ and factor $k$:

Prior distribution:

$$p(\theta_k^m) = \text{Beta}\left(\theta_k^m \mid a_0^\theta, b_0^\theta\right)$$

Variational distribution:

$$q(\theta_k^m) = \text{Beta}\left(\theta_k^m \mid \hat{a}_{mk}^\theta, \hat{b}_{mk}^\theta\right) \tag{55}$$

where

$$\hat{a}_{mk}^\theta = \sum_{d=1}^{D_m} \langle s_{kd}^m \rangle + a_0^\theta$$
$$\hat{b}_{mk}^\theta = b_0^\theta - \sum_{d=1}^{D_m} \langle s_{kd}^m \rangle + D_m \tag{56}$$

**Noise (Gaussian)**

For every view $m$, group $g$ and feature $d$:

Prior distribution $p(\tau_d^{gm})$:

$$p(\tau_d^{gm}) = \mathcal{G}\left(\tau_{dg}^{m} \,|\, a_0^\tau, b_0^\tau\right),$$

Variational distribution $q(\tau_d^{gm})$:

$$q(\tau_d^{gm}) = \mathcal{G}\left(\tau_d^{gm} \,|\, \hat{a}_d^{gm}, \hat{b}_d^{gm}\right) \tag{57}$$

where:

$$\begin{aligned} \hat{a}_d^{gm} &= a_0^\tau + \frac{N_g}{2} \\ \hat{b}_d^{gm} &= b_0^\tau + \frac{1}{2}\sum_{n=1}^{N_g}\langle\left(y_{nd}^{gm} - \sum_{k}^{K} w_{kd}^{m} z_{nk}^{g}\right)^2\rangle \end{aligned} \tag{58}$$

**b) Evidence Lower Bound**

Although computing the ELBO is not necessary in order to estimate the posterior distribution of the parameters, it is used to monitor the convergence of the algorithm. As shown in Equation (15), the ELBO can be decomposed into a sum of two terms: (1) the expected log likelihood under the current estimate of the posterior distribution of the parameters and (2) the KL divergence between the prior and the variational distributions of the parameters:

$$\mathcal{L} = \mathbb{E}_{q(X)} \ln P(Y|X) - \text{KL}\left(q(X)||p(X)\right) \tag{59}$$

**Log likelihood term** Assuming a Gaussian likelihood:

$$\begin{aligned} \mathbb{E}_{q(X)} \ln P(Y|X) = &-\sum_{m=1}^{M} \frac{ND_m}{2}\ln(2\pi) + \sum_{g=1}^{G}\frac{N_g}{2}\sum_{m=1}^{M}\sum_{d=1}^{D_m}\langle\ln(\tau_d^{gm})\rangle \\ &-\sum_{g=1}^{G}\sum_{m=1}^{M}\sum_{d=1}^{D_m}\frac{\langle\tau_d^{gm}\rangle}{2}\sum_{n=1}^{N_g}\left(y_{nd}^{m,g} - \sum_{k=1}^{K}\langle s_{kd}^m \hat{w}_{kd}^m\rangle\langle z_{nk}^g\rangle\right)^2 \end{aligned} \tag{60}$$

**KL divergence terms** Note that $\text{KL}\left(q(X)||P(X)\right) = \mathbb{E}_q(q(X)) - \mathbb{E}_q(P(X))$.
Below, we will write the analytical form for these two expectations.

**Sparse weights**

$$\begin{aligned} \mathbb{E}_q[\ln p(\hat{W}, S)] = &-\sum_{m=1}^{M}\frac{KD_m}{2}\ln(2\pi) + \sum_{m=1}^{M}\frac{D_m}{2}\sum_{k=1}^{K}\ln(\alpha_k^m) - \sum_{m=1}^{M}\frac{\alpha_k^m}{2}\sum_{d=1}^{D_m}\sum_{k=1}^{K}\langle(\hat{w}_{kd}^m)^2\rangle \\ &+ \langle\ln(\theta)\rangle\sum_{m=1}^{M}\sum_{d=1}^{D_m}\sum_{k=1}^{K}\langle s_{kd}^m\rangle + \langle\ln(1-\theta)\rangle\sum_{m=1}^{M}\sum_{d=1}^{D_m}\sum_{k=1}^{K}(1-\langle s_{kd}^m\rangle) \end{aligned} \tag{61}$$

$$\begin{aligned} \mathbb{E}_q[\ln q(\hat{W}, S)] = &-\sum_{m=1}^{M}\frac{KD_m}{2}\ln(2\pi) + \frac{1}{2}\sum_{m=1}^{M}\sum_{d=1}^{D_m}\sum_{k=1}^{K}\ln(\langle s_{kd}^m\rangle\sigma_{w_{kd}^m}^2 + (1-\langle s_{kd}^m\rangle)/\alpha_k^m) \\ &+\sum_{m=1}^{M}\sum_{d=1}^{D_m}\sum_{k=1}^{K}(1-\langle s_{kd}^m\rangle)\ln(1-\langle s_{kd}^m\rangle) - \langle s_{kd}^m\rangle\ln\langle s_{kd}^m\rangle \end{aligned} \tag{62}$$

**Non-sparse weights**

$$\mathbb{E}_q[\ln p(W)] = -\frac{DK}{2}\ln(2\pi) - \frac{1}{2}\sum_{m=1}^{M}\sum_{d=1}^{D_m}\sum_{k=1}^{K}\langle (w_{kd}^{m})^2\rangle$$
$$\mathbb{E}_q[\ln q(W)] = -\frac{DK}{2}(1+\ln(2\pi)) - \frac{1}{2}\sum_{m=1}^{M}\sum_{d=1}^{D_m}\sum_{k=1}^{K}\ln(\sigma^2_{w_{kd}}) \tag{63}$$

**Sparse factors**

$$\mathbb{E}_q[\ln p(\hat{Z},S)] = -\sum_{g=1}^{G}\frac{N_g K}{2}\ln(2\pi) + \sum_{g=1}^{G}\frac{N_g}{2}\sum_{k=1}^{K}\ln(\alpha_k^g) - \sum_{g=1}^{G}\frac{\alpha_k^g}{2}\sum_{n=1}^{N_g}\sum_{k=1}^{K}\langle(\hat{z}_{nk}^g)^2\rangle$$
$$+ \langle\ln(\theta)\rangle\sum_{g=1}^{G}\sum_{n=1}^{N_g}\sum_{k=1}^{K}\langle s_{nk}^g\rangle + \langle\ln(1-\theta)\rangle\sum_{g=1}^{G}\sum_{n=1}^{N_g}\sum_{k=1}^{K}(1-\langle s_{nk}^g\rangle) \tag{64}$$

$$\mathbb{E}_q[\ln q(\hat{Z},S)] = -\sum_{g=1}^{G}\frac{N_g K}{2}\ln(2\pi) + \frac{1}{2}\sum_{g=1}^{G}\sum_{n=1}^{N_g}\sum_{k=1}^{K}\ln(\langle s_{nk}^g\rangle\sigma^2_{z_{nk}^g} + (1-\langle s_{nk}^g\rangle)/\alpha_k^g)$$
$$+\sum_{g=1}^{G}\sum_{n=1}^{N_g}\sum_{k=1}^{K}(1-\langle s_{nk}^g\rangle)\ln(1-\langle s_{nk}^g\rangle) - \langle s_{nk}^g\rangle\ln\langle s_{nk}^g\rangle \tag{65}$$

**Non-sparse factors**

$$\mathbb{E}_q[\ln p(Z)] = -\frac{NK}{2}\ln(2\pi) - \frac{1}{2}\sum_{g=1}^{G}\sum_{n=1}^{N_g}\sum_{k=1}^{K}\langle(z_{nk}^g)^2\rangle$$
$$\mathbb{E}_q[\ln q(Z)] = -\frac{NK}{2}(1+\ln(2\pi)) - \frac{1}{2}\sum_{g=1}^{G}\sum_{n=1}^{N_g}\sum_{k=1}^{K}\ln(\sigma^2_{z_{nk}}) \tag{66}$$

**ARD precision for the weights**

$$\mathbb{E}_q[\ln p(\boldsymbol{\alpha})] = \sum_{m=1}^{M}\sum_{k=1}^{K}\Big(a_0^\alpha\ln b_0^\alpha + (a_0^\alpha - 1)\langle\ln\alpha_k\rangle - b_0^\alpha\langle\alpha_k\rangle - \ln\Gamma(a_0^\alpha)\Big)$$
$$\mathbb{E}_q[\ln q(\boldsymbol{\alpha})] = \sum_{m=1}^{M}\sum_{k=1}^{K}\Big(\hat{a}_k^\alpha\ln\hat{b}_k^\alpha + (\hat{a}_k^\alpha - 1)\langle\ln\alpha_k\rangle - \hat{b}_k^\alpha\langle\alpha_k\rangle - \ln\Gamma(\hat{a}_k^\alpha)\Big) \tag{67}$$

**Sparsity parameter of the weights**

$$\mathbb{E}_q\left[\ln p(\boldsymbol{\theta})\right] = \sum_{m=1}^{M}\sum_{k=1}^{K}\sum_{d=1}^{D_m}\big((a_0-1)\times\langle\ln(\pi_{d,k}^m)\rangle + (b_0-1)\langle\ln(1-\pi_{d,k}^m)\rangle - \ln(\mathrm{B}(a_0,b_0))\big)$$
$$\mathbb{E}_q\left[\ln q(\boldsymbol{\theta})\right] = \sum_{m=1}^{M}\sum_{k=1}^{K}\sum_{d=1}^{D_m}\big((a_{k,d}^m-1)\times\langle\ln(\pi_{d,k}^m)\rangle + (b_{k,d}^m-1)\langle\ln(1-\pi_{d,k}^m)\rangle - \ln(\mathrm{B}(a_{k,d}^m,b_{k,d}^m))\big) \tag{68}$$

**ARD precision for the factors**

$$\mathbb{E}_q[\ln p(\boldsymbol{\alpha})] = \sum_{g=1}^{G}\sum_{k=1}^{K}\Big(a_0^\alpha\ln b_0^\alpha + (a_0^\alpha - 1)\langle\ln\alpha_k\rangle - b_0^\alpha\langle\alpha_k\rangle - \ln\Gamma(a_0^\alpha)\Big)$$
$$\mathbb{E}_q[\ln q(\boldsymbol{\alpha})] = \sum_{g=1}^{G}\sum_{k=1}^{K}\Big(\hat{a}_k^\alpha\ln\hat{b}_k^\alpha + (\hat{a}_k^\alpha - 1)\langle\ln\alpha_k\rangle - \hat{b}_k^\alpha\langle\alpha_k\rangle - \ln\Gamma(\hat{a}_k^\alpha)\Big) \tag{69}$$

**Sparsity parameter of the factors**

$$\begin{aligned}\mathbb{E}_q\left[\ln p(\boldsymbol{\theta})\right] &= \sum_{g=1}^{G}\sum_{k=1}^{K}\sum_{n=1}^{N_g}\Big((a_0-1)\times\langle\ln(\pi^g_{n,k})\rangle+(b_0-1)\langle\ln(1-\pi^g_{n,k})\rangle-\ln(\mathrm{B}(a_0,b_0))\Big)\\ \mathbb{E}_q\left[\ln q(\boldsymbol{\theta})\right] &= \sum_{g=1}^{G}\sum_{k=1}^{K}\sum_{n=1}^{N_g}\Big((a^g_{k,n}-1)\times\langle\ln(\pi^g_{n,k})\rangle+(b^g_{k,n}-1)\langle\ln(1-\pi^g_{n,k})\rangle-\ln(\mathrm{B}(a^g_{k,n},b^g_{k,n}))\Big)\end{aligned} \tag{70}$$

**Noise**

$$\begin{aligned}\mathbb{E}_q[\ln p(\boldsymbol{\tau})] &= \sum_{m=1}^{M}D_m a_0^\tau\ln b_0^\tau+\sum_{g=1}^{G}\sum_{m=1}^{M}\sum_{d=1}^{Dm}(a_0^\tau-1)\langle\ln\tau_d^{gm}\rangle-\sum_{g=1}^{G}\sum_{m=1}^{M}\sum_{d=1}^{Dm}b_0^\tau\langle\tau_d^{gm}\rangle-\sum_{m=1}^{M}D_m\ln\Gamma(a_0^\tau)\\ \mathbb{E}_q[\ln q(\boldsymbol{\tau})] &= \sum_{g=1}^{G}\sum_{m=1}^{M}\sum_{d=1}^{D_m}\Big(\hat{a}^\tau_{dgm}\ln\hat{b}^\tau_{dgm}+(\hat{a}^\tau_{dgm}-1)\langle\ln\tau_d^{gm}\rangle-\hat{b}^\tau_{dgm}\langle\tau_d^{gm}\rangle-\ln\Gamma(\hat{a}^\tau_{dgm})\Big)\end{aligned} \tag{71}$$

##### 2.2.4 Applying the stochastic variational inference algorithm

###### a) The algorithm

The local dimension we chose to apply the Stochastic Variational Inference algorithm is the axis of the samples. Indeed, in large single-cells datasets, there is often more single cells than variables measured : $N = \sum_{g=1}^{G} N_g$ is often larger than $D = \sum_{m=1}^{M} D_m$.

Consequently, the only local hidden variables are the factors matrices $\mathbf{Z^g} = (z^g_{nk})_{1 \leq n \leq N_g,\ \ 1 \leq k \leq K}$ which contains the factors expressions for each sample $n$ belonging to group $g \in [1, G]$. When adding the spike-and-slab prior over the factors matrices $\mathbf{Z^g}$ (sparse factors), the only local hidden variables are the two matrices $\hat{\mathbf{Z}}^\mathbf{g}$ and $\mathbf{S^g}$ (whose term-wise product gives $\mathbf{Z^g}$) for each $g \in [1, G]$.

All other hidden variables are global : $\tau^\mathbf{gm}$, $\hat{\mathbf{W}}^\mathbf{m}$ and $\mathbf{S^m}$ (whose term-wise product gives $\mathbf{W}^m$), $\alpha^\mathbf{m}, \theta^\mathbf{m}$, as well as $\alpha^\mathbf{g}$ and $\theta^\mathbf{g}$ when adding the spike-and-slab prior over the factor matrices $\mathbf{Z^g}$.

Hence, we chose to apply the following SVI algorithm to speed-up the inference of the MOFA model on datasets with a large number of samples $N$:

**Algorithm 3** Stochastic mean-field variational inference for MOFA 2.0 with sparse factors

Initialise randomly the parameters of the global variables $\{\tau^\mathbf{gm}, \hat{\mathbf{W}}^\mathbf{m}, \mathbf{S^m}, \alpha^\mathbf{m}, \theta^\mathbf{m}, \alpha^\mathbf{g}, \theta^\mathbf{g}\}$.
Initialise the step size $\rho^{(t=0)}$
**repeat**
  sample $\mathcal{B}$ a mini-batch of samples of size $S << N$
  **for** each local variational parameter $\phi^g_{nk}$ of nodes $\{z^{\hat{g}}_{nk}, s^g_{nk}\}$ such that $n$ is in batch $\mathcal{B}$ **do**
    $\phi^{(t+1)}_{nk}$ is the updated parameter $\phi_{nk}$ following the classic VI update equation
  **end for**
  **for** each global variational parameter $\lambda$ of nodes $\{\tau^\mathbf{gm}, \hat{\mathbf{W}}^\mathbf{m}, \mathbf{S}^m, \alpha^\mathbf{m}, \theta^\mathbf{m}, \alpha^\mathbf{g}, \theta^\mathbf{g}\}$ **do**

$$\lambda^{(t+1)} = (1 - \rho^{(t)})\lambda^{(t)} + \rho^{(t)}\lambda^{(t+1)}_{\mathcal{B}} \tag{72}$$

    where $\lambda^{(t+1)}_{\mathcal{B}}$ is the updated parameter $\lambda$ following the classic VI update equation,
    but considering the selected batch $\mathcal{B}$ repeated $N/S$ times instead of the full dataset.
  **end for**
**until** ELBO convergence

###### b) Choosing the hyperparameters

**Batch size**: Using small batch sizes for inference on large datasets is often interesting : the number of iterations required to converge is multiplied by less than $n$ while the time per iteration is divided by $n$. We may explain this fact by a faster improvement in the very first iterations of the guess of the global parameters : indeed, we do not need to know the observations for each cell of the dataset to improve the guess of the global variables from their random initialization. We did our experiments with a batch size varying from 10% to 50% of the full dataset.

**Learning rate**: We used the following function to iteratively decrease the learning rate:

$$\rho^{(t)} = \frac{\tau}{(1 + \kappa t)^{3/4}}$$

This function has two hyperparameters:

- The forgetting rate $\kappa$: controls the decay of the learning rate, with large $\kappa$ leading to faster decays. In the natural gradient setting, $\kappa$ also controls how quickly information from previous iterations is forgotten.

- The delay $\tau$: determines the initial learning rate. In the natural gradient setting, the larger $\tau$ the more early iterations are down-weighted.

Figure 7 shows the effect on the learning rate of varying the two hyperparameters.

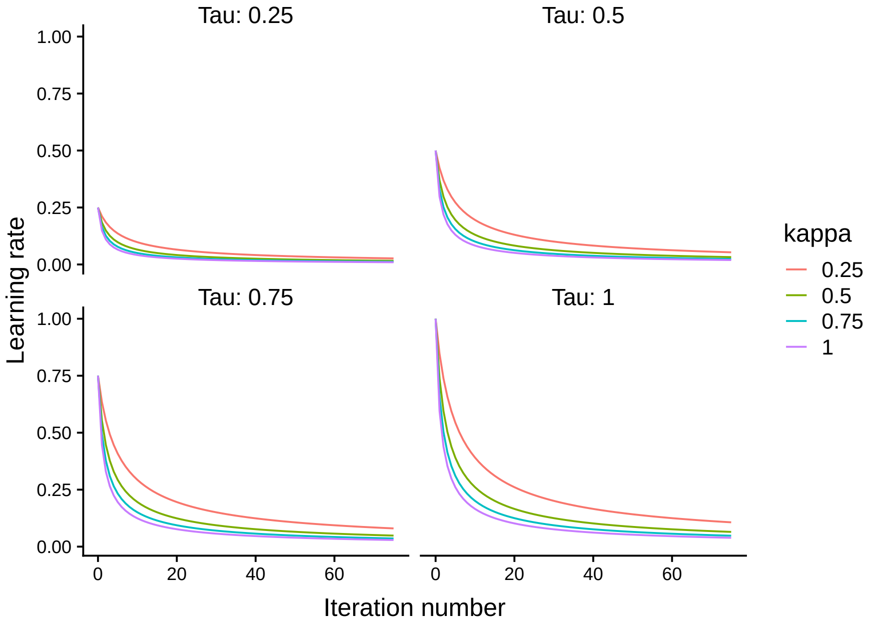

Figure 7

### References

[1] S.-I. Amari. "Natural Gradient Works Efficiently in Learning". In: *Neural Comput.* 10.2 (Feb. 1998), pp. 251–276.

[2] R. Argelaguet et al. "Multi-Omics Factor Analysis-a framework for unsupervised integration of multi-omics data sets". In: *Mol Syst Biol* 14.6 (2018), e8124. ISSN: 1744-4292 (Electronic) 1744-4292 (Linking). DOI: 10.15252/msb.20178124.

[3] J. Beal. "Variational algorithms for approximate bayesian inference". University College London, 2003.

[4] C. M. Bishop. "Pattern recognition". In: *Machine Learning* 128 (2006), pp. 1–58.

[5] D. M. Blei, A. Kucukelbir, and J. D. McAuliffe. "Variational Inference: A Review for Statisticians". In: *arXiv e-prints*, arXiv:1601.00670 (Jan. 2016), arXiv:1601.00670. arXiv: 1601.00670 [stat.CO].

[6] J. Cao et al. "The single-cell transcriptional landscape of mammalian organogenesis". In: *Nature* 566.7745 (2019), pp. 496–502. DOI: 10.1038/s41586-019-0969-x.

[7] M. Emtiyaz Khan and D. Nielsen. "Fast yet Simple Natural-Gradient Descent for Variational Inference in Complex Models". In: *arXiv e-prints*, arXiv:1807.04489 (July 2018), arXiv:1807.04489. arXiv: 1807.04489 [stat.ML].

[8] C. Gao, C. D. Brown, and B. E. Engelhardt. "A latent factor model with a mixture of sparse and dense factors to model gene expression data with confounding effects". In: *arXiv e-prints*, arXiv:1310.4792 (2013), arXiv:1310.4792. arXiv: 1310.4792 [stat.AP].

[9] M. Hoffman et al. "Stochastic Variational Inference". In: *arXiv e-prints*, arXiv:1206.7051 (June 2012), arXiv:1206.7051. eprint: 1206.7051.

[10] V. Hore. "Latent Variable Models for Analysing Multidimensional Gene Expression Data". PhD thesis. University of Oxford, 2015.

[11] M. I. Jordan et al. "An Introduction to Variational Methods for Graphical Models". In: *Mach. Learn.* 37.2 (Nov. 1999), pp. 183–233.

[12] E. Kiciman, D. Maltz, and J. C. Platt. "Fast Variational Inference for Large-scale Internet Diagnosis". In: *Advances in Neural Information Processing Systems 20.* Ed. by J. C. Platt et al. Curran Associates, Inc., 2008, pp. 1169–1176.

[13] A. Kristiadi. *Natural Gradient Descent.* https://wiseodd.github.io/techblog/2018/03/14/natural-gradient/. Blog. 2019.

[14] D. J. MacKay. "Bayesian methods for backpropagation networks". In: *Models of neural networks III.* Springer, 1996, pp. 211–254.

[15] J. Martens. "New insights and perspectives on the natural gradient method". In: *arXiv e-prints*, arXiv:1412.1193 (Dec. 2014), arXiv:1412.1193. arXiv: 1412.1193 [cs.LG].

[16] T. J. Mitchell and J. J. Beauchamp. "Bayesian variable selection in linear regression". In: *Journal of the American Statistical Association* 83.404 (1988), pp. 1023–1032.

[17] K. P. Murphy. *Machine Learning: A Probabilistic Perspective.* The MIT Press, 2012. ISBN: 0262018020, 9780262018029.

[18] R. Ranganath, S. Gerrish, and D. M. Blei. "Black Box Variational Inference". In: *arXiv e-prints*, arXiv:1401.0118 (Dec. 2013), arXiv:1401.0118. arXiv: 1401.0118 [stat.ML].

[19] R. Ranganath et al. "An Adaptive Learning Rate for Stochastic Variational Inference". In: *Proceedings of the 30th International Conference on International Conference on Machine Learning - Volume 28.* ICML'13. JMLR.org, 2013, pp. II-298–II-306.

[20] H. Robbins and S. Monro. "A stochastic approximation method". In: *The Annals of Mathematical Statistics* 22.3 (1951), pp. 400–407.

[21] L. K. Saul, T. Jaakkola, and M. I. Jordan. "Mean Field Theory for Sigmoid Belief Networks". In: *arXiv e-prints*, cs/9603102 (Feb. 1996), cs/9603102. arXiv: cs/9603102.

[22] J. C. Spall. *Introduction to stochastic search and optimization: estimation, simulation, and control.* Hoboken, N.J: J. Wiley, 2003.

[23] V. Svensson, R. Vento-Tormo, and S. A. Teichmann. "Exponential scaling of single-cell RNA-seq in the past decade". In: *Nature Protocols* 13 (Mar. 2018),

[24] M. K. Titsias and M. Lázaro-Gredilla. “Spike and slab variational inference for multi-task and multiple kernel learning”. In: *Advances in neural information processing systems*. 2011, pp. 2339–2347.

[25] C. Zhang et al. “Advances in Variational Inference”. In: *arXiv e-prints*, arXiv:1711.05597 (Nov. 2017), arXiv:1711.05597. arXiv: 1711.05597.

[26] J.-h. Zhao and P. L. Yu. “A note on variational Bayesian factor analysis”. In: *Neural Networks* 22.7 (2009), pp. 988–997. ISSN: 0893-6080. DOI: https://doi.org/10.1016/j.neunet.2008.11.002.
